# Investigation of the threonine metabolism of *Echinococcus multilocularis*: the threonine dehydrogenase as a potential drug target in alveolar echinococcosis

**DOI:** 10.1101/2024.07.27.605433

**Authors:** Marc Kaethner, Pascal Zumstein, Matías Preza, Philipp Grossenbacher, Anissa Bartetzko, Martin Lochner, Stefan Schürch, Clement Regnault, Daniel Villalobos Ramírez, Britta Lundström-Stadelmann

**Affiliations:** Institute of Parasitology, Vetsuisse Faculty, University of Bern, Bern, Switzerland; Graduate School for Cellular and Biomedical Sciences, University of Bern, Bern, Switzerland; Institute of Biochemistry and Molecular Medicine, University of Bern, Bern, Switzerland; Department of Chemistry, Biochemistry and Pharmaceutical Sciences, University of Bern, Bern, Switzerland; Integrated Protein Analysis - Mass Spectrometry unit, MVLS Shared Research Facilities, College of Medical, Veterinary and Life Sciences, University of Glasgow, Glasgow, United Kingdom; Department of Bioinformatics, University of Würzburg, Würzburg, Germany; Multidisciplinary Center for Infectious Diseases, University of Bern, Bern, Switzerland

**Keywords:** Echinococcus multilocularis, cestode, threonine metabolism, target-based screening, TDH, disulfiram, sanguinarine

## Abstract

Alveolar echinococcosis (AE) is a severe zoonotic disease caused by the metacestode stage of the fox tapeworm *Echinococcus multilocularis*. We recently showed that *E. multilocularis* metacestode vesicles scavenge large amounts of L-threonine from the culture medium that were neither stored nor overused for protein synthesis. This motivated us to study the effect of L-threonine on the parasite and how it is metabolized. We established a novel metacestode vesicle growth assay with an automated readout, which showed that L-threonine treatment led to significantly increased parasite growth. In addition, L-threonine increased the formation of novel metacestode vesicles from primary parasite cell cultures in contrast to the non-proteinogenic threonine analog 3-hydroxynorvaline. Tracing of [U-^13^C]-L-threonine and metabolites in metacestode vesicles and culture medium resulted in the detection of [U-^13^C]-labeling in aminoacetone and glycine, indicating that L-threonine was metabolized by threonine dehydrogenase (TDH). In addition, the detection of [^13^C_2_]-glutathione, suggested that *E. multilocularis* metacestode vesicles synthesize glutathione via L-threonine-derived glycine. EmTDH-mediated threonine metabolism in the *E. multilocularis* metacestode stage was further confirmed by quantitative real-time PCR, which demonstrated high expression of *emtdh* in *in vitro* cultured metacestode vesicles and also in metacestode samples obtained from infected animals. EmTDH was enzymatically active in metacestode vesicle extracts. Thus, the drugs disulfiram, myricetin, quercetin, sanguinarine and seven quinazoline carboxamides were assessed for inhibition of recombinantly expressed EmTDH, and the most potent inhibitors disulfiram, myricetin and sanguinarine were further tested for activity against *E. multilocularis* metacestode vesicles and primary parasite cells. Sanguinarine exhibited significant *in vitro* activity and IC_50_-values for metacestode vesicles, primary parasite cells, as well as mammalian cells were determined. Our results suggest that sanguinarine treatment should be further assessed *in vivo* employing suitable AE mouse models. Furthermore, the EmTDH assay could serve as high-throughput target-based discovery platform for novel anti-echinococcal compounds.

## 1. Introduction

Platyhelminth parasites pose major burdens on human and veterinary health worldwide. The class Cestoda includes the fox tapeworm *Echinococcus multilocularis* which causes the severe zoonotic disease alveolar echinococcosis (AE) in humans and other mammals such as various species of simians and dogs (1–3). Worldwide, approximately 18,000 new human AE cases occur annually which correspond to 688,000 disability adjusted life years (4). The distribution of *E. multilocularis* is restricted to the Northern Hemisphere and more than 90% of the cases occur in China (5). The infection is acquired via oral uptake of *E. multilocularis* eggs, from which infective oncospheres hatch and establish themselves in the liver as metacestodes (6). Metacestodes grow infiltratively into the liver and surrounding organs, and may form metastases to more distant body locations (6). AE is fatal if left untreated and curative surgery is applicable in 20 to 50% of cases in countries with well-developed and -accessible health infrastructure (7). Nonsurgical interventions consist of lifelong therapy with daily intake of either albendazole (10 to 15 mg/kg/day divided in two doses) or mebendazole (40 to 50 mg/kg/day divided in three doses) (8). However, treatment with these benzimidazoles can induce adverse effects including severe liver toxicity affecting up to 6.9 % of patients (9). The resulting treatment discontinuation can lead to recurrence of parasite growth (9,10), which has been proposed to be caused by the undifferentiated stem cells of metacestodes that are not affected by albendazole or mebendazole (11,12). The undifferentiated stem cells are an integral part of the germinal layer (GL), which constitutes the actual parasite tissue and forms the inner layer of the fluid-filled metacestode vesicles (13). In addition, the GL contains differentiated cell types such as muscle cells, nerve cells, glycogen storage cells and subtegumentary cytons (13,14). The inner fluid of metacestodes, also called vesicle fluid (VF) for *in vitro* grown metacestode vesicles (15), stores nutrients such as glucose, as well as various amino acids (Ritler et al., 2019) and proteins (17). The GL is further surrounded by a syncytial tegument and an outer acellular and carbohydrate-rich laminated layer (18,19). The metacestode stem cells are the only cells of the metacestode tissue that undergo continuous proliferation (13) and, in order to be effective, new treatment options must target these stem cells (20).

In the search for new anthelmintics, efforts have been made to develop new assays for whole-organism-based drug screening that allow for efficient *in vitro* screening of drug libraries, either applying novel compounds, or repurposed drugs (21). For platyhelminths many different *in vitro* assays have been established, which allow for objective readout methods to evaluate activities of compounds against the trematodes *Schistosoma mansoni* (22–25) or *Fasciola hepatica* (26,27). Regarding cestodes, objective assays have been developed for *Taenia crassiceps* cysticerci and *Mesocestoides corti* tetrathyridia (28,29). In the case of *E. multilocularis* and the closely related *E. granulosus sensu stricto* the development of drug screening assays led to the establishment of well-defined drug screening cascades (20,30). These include *in vitro* assays on the disease-causing metacestode stage and drug efficacy is assessed via damage marker and viability assays on metacestodes (30–32), protoscoleces and isolated GL cells (31,33), which consist of up to 83% of undifferentiated stem cells (13). While these assays can help in the discovery of novel anti-echinococcal compounds, the translation of these discoveries into a new drug treatment options against AE faces very high hurdles: investigations on absorption, pharmacokinetics, biodistribution, toxicity, metabolism and other preclinical and clinical studies are highly cost intensive undertakings, and the expected market return of new drugs against AE is relatively low (20,34). This challenge can be addressed by drug repurposing (35) and therefore, the screening of compounds with already known pharmacological profiles against *E. multilocularis* represents an important approach towards the discovery of potential new treatment options (36). However, the identification of novel compounds via whole-organism-based screens is costly, requires much labor and resources (37,38). Another approach is the focus on target-based drug screening for which a profound understanding of parasite biology and host-parasite interaction is required (38).

Studies on the genome of *E. multilocularis* have shown that this parasite has adapted to its life within a host and this has rendered the parasite dependant on scavenging nutrients from its host (39). Examples are the loss of pathways for the *de novo* synthesis of fatty acids, purines and pyrimidines, cholesterol and amino acids (39,40). In a previous study, we investigated the uptake of nutrients and secretion of metabolites by *E. multilocularis* metacestode vesicles in a controlled *in vitro* setting (Ritler et al., 2019). We found that metacestode vesicles take up high amounts of L-threonine from the culture medium and secrete glycine (Ritler et al., 2019). L-threonine was not overrepresented in VF or GL cells and thus the uptake could not be explained by simple storage within these compartments (Ritler et al., 2019). Albeit the laminated layer antigen Em2 is rich in threonine (41), threonine was not overrepresented in proteins of the GL or laminated layer of *in vitro* cultured metacestode vesicles (Ritler et al., 2019). Thus, there must be another reason for the high threonine consumption of *E. multilocularis* metacestode vesicles *in vitro*.

In the model helminth *Caenorhabditis elegans*, L-threonine can be metabolized by three different enzymes, threonine deaminase (TD), threonine dehydrogenase (TDH) and threonine aldolase (TA) (42,43) (Figure 1). TD metabolizes threonine to α-ketobutyrate and ammonia (44). TDH catabolizes threonine to 2-amino-3-ketobutyrate, which is later metabolized via the 2-amino-3-ketobutyrate coenzyme A ligase (KBL) to glycine and acetyl-coenzyme A (45), or can decarboxylate non-enzymatically to aminoacetone (46,47). TA-mediated threonine catabolism generates glycine and acetaldehyde (48). In contrast to *C. elegans*, little is known about the metabolism of L-threonine concerning parasites. Threonine catabolism via TD was investigated in the protozoan *Entamoeba histolytica* (49), the nematodes *Heligmosomoides polygyrus* and *Nippostrongylus brasiliensis* (50,51) and the trematode *Fasciola indica* (52). TDH-mediated threonine catabolism has only been characterized in the protozoan parasite *Trypanosoma brucei* (53,54). Here, TbTDH has been proposed as potential drug target since human *tdh* is a nonfunctional pseudogene (55,56). To the best of our knowledge, no studies have been done on TA-mediated threonine catabolism in parasites and also studies regarding threonine metabolism in cestodes are lacking.

**Fig 1.**
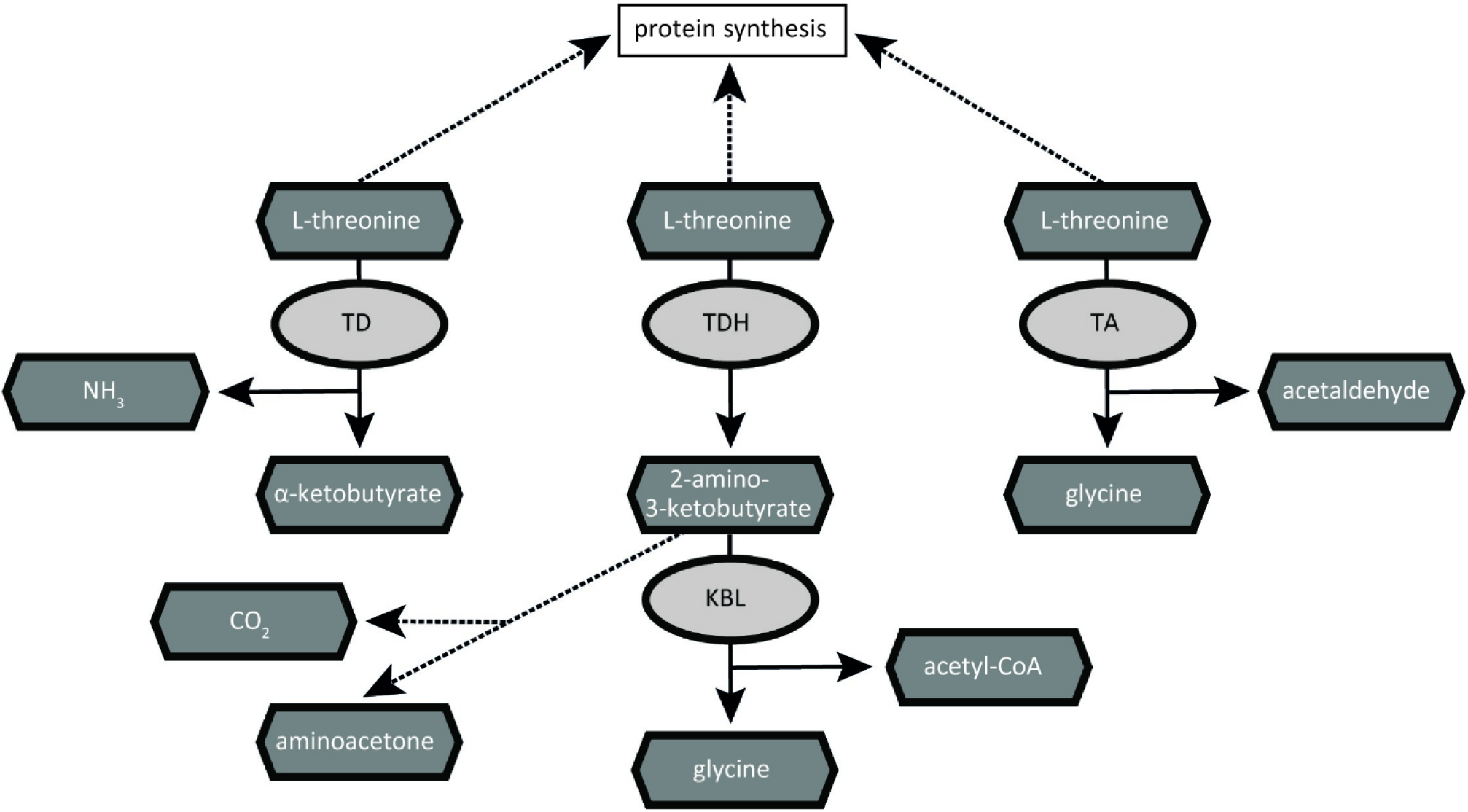
Pathways of threonine catabolism in the model helminth *C. elegans*. L-threonine can be used as substrate by the three different enzymes threonine deaminase (TD), threonine dehydrogenase (TDH) and threonine aldolase (TA) (43). Upon metabolization of threonine by TD, α-ketobutyrate and ammonia are generated. TDH metabolizes threonine to 2-amino-3-ketobutyrate, which is further metabolized by the 2-amino-3-ketobutyrate coenzyme A ligase (KBL) to glycine and acetyl-coenzyme A. TA generates glycine and acetaldehyde upon degradation of threonine. Enzymes are depicted in light grey and metabolites in dark grey. Reactions are represented by arrows (enzymatically) or dashed arrows (non-enzymatically).

The aim of this study was to unravel the presence and relevance of an active threonine metabolism for *E. multilocularis in vitro*. In the here presented work, we showed that an active threonine metabolism positively affects growth and development of *E. multilocularis in vitro*. Furthermore, we identified the enzymes responsible for threonine catabolism of *E. multilocularis in vitro* and established an enzymatic assay to test potential inhibitors that could be employed for drug-mediated treatment options against AE in the future.

## 2. Material and methods

### 2.1. Chemicals and reagents

If not stated otherwise, all chemicals were purchased from Sigma-Aldrich (Buchs, Switzerland) and all plastic ware was purchased from Sarstedt (Sevelen, Switzerland). Dulbeccos’s modified Eagle medium (DMEM) and penicillin and streptomycin (10,000 Units/mL penicillin, 10,000 μg/mL streptomycin) were from Gibco (Fisher Scientific AG, Reinach, Switzerland). DMEM without threonine and glucose was purchased from Teknova (Hollister, California, USA). Fetal bovine serum (FBS) and Trypsin/EDTA (0.05% Trypsin/0.02% EDTA) were from Bioswisstec (Schaffhausen, Switzerland). Quinazoline carboxamides (QCs) were synthesized as described in S1 File. Reuber rat hepatoma cells (RH, H-4-II-E) and human foreskin fibroblasts (HFFs) were purchased from ATCC (Molsheim Cedex, France). Murine Hepa 1-6 cells were kindly provided by Magali Roques (Institute of Cell Biology, University of Bern).

### 2.2. Mice and ethics statement

*E. multilocularis* strain H95 was maintained in female BALB/c mice (Charles River Laboratories, Sulzheim, Germany). Mice were kept under controlled conditions with a twelve hours light/dark cycle, a temperature of 21 – 23°C and a relative humidity of 45 – 55%. Food and water were provided *ad libitum,* and cages were enriched with mouse houses (Tecniplast, Gams, Switzerland), tunnels (Zoonlab, Castrop-Rauxel, Germany) and nestlets (Plexx, Elst, Netherlands). All animals were treated in compliance with the Swiss Federal Protection of Animals Act (TSchV, SR455), and experiments were approved by the Animal Welfare Committee of the canton of Bern under the license numbers BE30/19 and BE2/22.

### 2.3. Culture of *E. multilocularis* metacestode vesicles

*E. multilocularis* metacestode vesicles (strain H95) were cultured as previously described (30). Metacestode material was aseptically collected from intraperitoneally infected BALB/c mice and pressed through a conventional tea strainer (Migros, Bern, Switzerland). The material was incubated overnight at 4°C in PBS containing penicillin (100 U/mL), streptomycin (100 µg/mL) and tetracycline (10 µg/mL) and the next day, 1.5 mL of pure parasite material was co-cultured with RH cells in DMEM supplemented with 10% FBS, penicillin (100 U/mL), streptomycin (100 µg/mL) and tetracycline (5 µg/mL) at 37°C under humid, 5% CO_2_ atmosphere.

### 2.4. Effect of threonine on *E. multilocularis*

#### 2.4.1. Development of an *E. multilocularis* metacestode vesicle growth assay

Metacestode vesicle growth was analyzed via a newly developed growth assay using automated image-based analysis in ImageJ via scripts that enables fast, precise, and objective measurements of *E. multilocularis* metacestode vesicles by providing a mean diameter of 360 diameter measurements. The pre-processing of the images was performed via a cleanup algorithm adapted from a code for the measurement of tumor spheroids (57). The scripts were validated with n=50 metacestode vesicles placed individually in 24-well plates and photographed using a Nikon SMZ18 stereo microscope (Nikon, Basel, Switzerland) at 1X magnification. The metacestode vesicles were moved within the well by circular movement of the plate to get three different images of the same metacestode vesicle. The resulting 150 images were randomly numbered and measured in a blinded manner by three different methods: a) manually in ImageJ version 1.54g with two diameters and calculations of mean values; b) by an automated macro measuring 360 diameters giving mean values as a result (S2 File); c) by a semi-automated version consisting of the automated script from b) and additionally one step prior to the measurement in which the metacestode vesicle is encircled by the user (S3 File). Thus, based on these three approaches, the mean diameter and SD was calculated for each of the 50 metacestode vesicles. The scripts were validated by comparing the mean values of the metacestode vesicle diameters measured with the automated script (b) or the semi-automated script (c) to the measurements performed manually (a) in ImageJ via two-sample two-tailed students t-tests with equal variance in R version 4.3.0 and Bonferroni-correction. *p*-values of *p*<0.05 were considered to be significant. Additionally, the internal variation between the three photos of the same metacestode vesicle was calculated for all individual metacestode vesicles based on each of the three measurement methods. Given are the diameter variation for the same metacestode vesicle with mean and SD values for each of the measurement methods.

#### 2.4.2. Effect of L-threonine on *E. multilocularis* metacestode vesicles

We performed a preliminary experiment to get an idea what range of L-threonine would be suitable to be tested in a growth assay with *E. multilocularis* metacestode vesicles. For this we used DMEM without L-threonine and glucose, added 1 mM L-threonine and conditioned it by 10^6^ RH cells for six days at 37°C under a humid CO_2_ atmosphere. We sterile filtered the medium and added L-threonine (or water as control) in concentrations of 2, 4, 8 and 12 mM. Single metacestode vesicles were photographed and cultured in 1.5 mL of the different media in wells of a 24-well plate in triplicates for four days under a humid, microaerobic atmosphere (85% N_2_, 10% CO_2_, 5% O_2_). Supernatant samples were taken and stored at -20°C for measurement of the concentration of L-threonine via high-performance liquid chromatography (HPLC) at the Department of Chemistry, Biochemistry and Pharmaceutical Sciences, University of Bern (see S4 File). Metacestode vesicle size was measured via the semi-automated script (2.4.1.c, S3 File) and respective reduction of L-threonine in the culture medium was measured by HPLC. Shapiro-Wilk tests in R showed normal distribution of the data between all conditions and significance in the reduction of L-threonine in culture medium was assessed via multiple two sample two-tailed students t-tests and subsequent Bonferroni correction in R. Bonferroni-correct *p*-values of *p*<0.05 were considered to be significant. Shown are the metacestode vesicle diameters with mean values and SDs, as well as the reduction of L-threonine in the culture medium with mean values and SDs.

For the metacestode growth assay, metacestode vesicles cultured for three to four months with a mean size of 3.3 mm (± 0.5 mm) were changed to an axenic culture system without RH cells as described by others (58). A6 medium was prepared from low glucose DMEM (1 g/L glucose) supplemented with 10% FBS, penicillin (100 U/mL), streptomycin (100 µg/mL) and tetracycline (5 µg/mL) by conditioning with 10^6^ RH cells for six days at 37°C under a humid CO_2_ atmosphere and subsequent sterile filtration. The medium was stored at 4°C not longer than one week. n=24 single metacestode vesicles per condition were distributed individually in 24-well plates and incubated in 1.5 mL A6 medium. In a first experiment, L-threonine was added to final concentrations of 1, 2, or 4 mM. Alternatively, D-threonine was added to 4 mM final concentration. An equal amount of distilled water was added to the control. In an independent second experiment, we also wanted to assess the effect of the non-proteinogenic threonine analogue 3-hydroxynorvaline (3-HNV) (59,60). Thus, we performed an assay in which we supplemented 4 mM 3-HNV, a combination of 4 mM 3-HNV and 4 mM L-threonine, or the respective amount of distilled water to the media of individually placed metacestode vesicles. Both experiments were performed two times independently with 24 replica per condition and metacestode vesicles were incubated under microaerobic conditions and medium changes were performed once a week.

For assessment of parasite growth, metacestode vesicles were photographed at the start and the end of the experiment using a Nikon SMZ18 stereo microscope at 0.75X magnification. A lower magnification was chosen than in 2.4.1 due to the expected increase of metacestode vesicle diameters after six weeks of incubation. Metacestode vesicle diameters were assessed via the automated script (2.4.1.b, S2 File) and in case the macro did not work perfectly (due to metacestode vesicles being too close to the border of the well), images were processed with the modified, semi-automated version of the script (2.4.1.c, S3 File) in which the metacestode vesicle is manually encircled. The relative growth of each individual metacestode vesicle within six weeks was calculated in relation to the water control. Metacestode vesicles that had collapsed until week six were excluded from the analysis. The results were compared by statistical analysis using multiple two sample Welch tests with subsequent Bonferroni correction in R. Bonferroni-corrected *p*-values of *p*<0.05 were considered to be significant.

#### 2.4.3. Isolation of GL cells from *E. multilocularis* metacestode vesicles

GL cells were isolated as described by a recently updated protocol (30). In short, conditioned DMEM (cDMEM) was prepared by culturing high-glucose DMEM supplemented with 10% FBS, penicillin (100 U/mL), streptomycin (100 µg/mL) and tetracycline (5 µg/mL) with 10^6^ Rh cells in 50 mL medium for six days, and 10^7^ cells in 50 mL medium for four days at 37°C under a humid, 5% CO_2_ atmosphere and after sterile filtration, combining them 1:1. Six-months-old metacestode vesicles were incubated in distilled water for two minutes, washed with PBS and mechanically broken by a pipette. The vesicle tissue (VT) was washed in PBS and incubated in eight volumes trypsin-EDTA solution at 37°C for 30 minutes. GL cells were extracted by filtering through a 30 µm mesh (Sefar AG, Heiden, Switzerland) and separated from calcareous corpuscles by short centrifugation (50 x g, 30 seconds). The cells were centrifuged, re-suspended in cDMEM and a 1:100 dilution was used to measure the OD_600_. An OD_600_ value of 0.1 of this dilution was defined as one arbitrary unit (AU) per µL of the undiluted cell suspension. 1 AU corresponded to 0.93 ± 0.17 µg total protein for eight different GL cell isolations of this study, as determined by bicinchoninic acid (BCA) assay using the Pierce™ BCA Protein Assay Kit (Fisher Scientific AG, Reinach, Switzerland). 1,000 AU of GL cells were cultured in five mL cDMEM at 37°C overnight under a humid nitrogen atmosphere. The next day, 2,000 AU of GL cells were combined and further cultured for three hours at 37°C under a humid nitrogen atmosphere.

#### 2.4.4. Vesicle formation assay

Vesicle formation assays were carried out as described by Hemer and Brehm (2012) with a few modifications such as a microaerobic atmosphere and a less enriched medium. In short, 150 AU of *E. multilocularis* GL cells were cultured in a 96-well plate in high glucose DMEM containing 1% FBS and penicillin (100 U/mL), streptomycin (100 µg/mL) and tetracycline (5 µg/mL) under a humid, microaerobic atmosphere. In a first experiment, 4 mM L-threonine, 4 mM D-threonine, or the respective amount of distilled water was added to the GL cells. In an independent second experiment, 4 mM 3-HNV, a combination of 4 mM L-threonine and 4 mM 3-HNV or the respective amount of distilled water was added. Both experiments were setup in four biological replica per condition with two independent experiments. Three times a week, half of the medium in each well was changed. After two weeks, newly formed metacestode vesicles were counted in a blinded manner. Shapiro-Wilk tests showed normal distribution with *p*>0.05 for all groups of each experiment. Statistical analyses were performed using multiple two-tailed students t-tests with equal variance. The Bonferroni-corrected *p*-values of *p*<0.05 were considered to be significant.

### 2.5. Tracing [U-^13^C] L-threonine and metabolites in *E. multilocularis* metacestode vesicles

We studied how L-threonine is metabolized in *E. multilocularis in vitro* by tracing [U-^13^C]-L-threonine and metabolites in metacestode vesicles and culture medium. 3 mL of two- to three-months-old *E. multilocularis* metacestode vesicles were cultured in 3 mL DMEM without threonine or glucose, supplemented with 0.2% FBS, 110 mg/L sodium pyruvate, 4.5 g/L glucose, 4 mM L-glutamine and 5 mM unlabeled threonine or 5 mM [U-^13^C]-L-threonine, respectively, at 37°C for 24 hours under a humid, microaerobic atmosphere. Control medium (CM) without metacestode vesicles was incubated under the same conditions. Each condition was set up in four biological replicates. Medium samples of CM and assay medium in which metacestode vesicles were incubated (VM), metacestode VF and metacestode VT were extracted in an ice-cold buffer consisting of HPLC grade chloroform:methanol:water (1:3:1 ratio). Medium samples were centrifuged (13,000 x g, 5 minutes, 4°C), 10 µL of supernatant were mixed with 400 µL of extraction buffer by vortexing and centrifuged again (13,000 x g, 5 minutes, 4°C). The supernatant was stored at -80°C. Metacestode vesicles were washed three times in 50 mL ice-cold PBS and then mechanically disrupted with a pipette. The VF was centrifuged (13,000 x g, 5 minutes, 4°C) and 10 µL of supernatant were mixed with 400 µL of extraction buffer by vortexing. The sample was centrifuged again (13,000 x g, 5 minutes, 4°C) and the supernatant was stored at -80°C. The metacestode VT pellet was washed three times with 1 mL ice-cold PBS and a centrifugation step (500 x g, 5 minutes, 4°C). The pellet was homogenized in 10 mL extraction buffer by vortexing with a 5 mm glass bead (30 steps of vortexing (10 seconds) and resting on ice (50 seconds)). The sample was centrifuged (4700 x g, 5 minutes, 4°C), the supernatant was centrifuged again (13,000 x g, 5 minutes, 4°C) and the supernatant was stored at -80°C. Blanks consisted of extraction buffer using the same tubes and vortexing/centrifugation steps. For tracing of [U-^13^C]-L-threonine and metabolites, samples were analyzed via Hydrophilic interaction liquid chromatography (HILIC) on a Dionex UltiMate 3000 RSLC system (Thermo Fisher Scientific, Hemel Hempstead, UK) using a ZIC-pHILIC column (150 mm × 4.6 mm, 5 μm column, Merck Sequant). The column was maintained at 25°C and samples were eluted over 26 min at a flow rate of 0.3 mL/min with a linear gradient over 15 min from an initial ratio of 80% acetonitrile (B) and 20% 20 mM ammonium carbonate in water (A) to 20% B and 80% A, followed by 95% A and 5% B for 2 min followed by re-equilibration at 80% B and 20% A for 9 min. The injection volume was 10 μL and samples were maintained at 5°C prior to injection. For the MS analysis, a Thermo Orbitrap QExactive (Thermo Fisher Scientific) was operated in polarity switching mode and the MS settings were Resolution 70,000, AGC 1e6, *m*/z range 70– 1050, sheath gas 40, auxiliary gas 5, sweep gas 1, probe temperature 150°C and capillary temperature 320°C. The samples were analyzed in positive mode ionization (source voltage +3.8 kV, S-Lens RF Level 30.00, SLens Voltage 25.00 V, Skimmer Voltage 15.00 V, Inject Flatopole Offset 8.00 V, Bent Flatapole DC 6.00 V) and negative mode ionization (source voltage -3.8 kV). The calibration mass range was extended to cover small metabolites by inclusion of low-mass calibrants with the standard Thermo calmix masses (below *m/z* 138), butylamine (C_4_H_11_N_1_) for positive ion electrospray ionisation (PIESI) mode (*m/z* 74.096426) and COF3 for negative ion electrospray ionisation (NIESI) mode (*m/z* 84.9906726). For each sample subset (medium, VF and VT), LC-MS raw data was processed with IDEOM (62) which uses the XCMS (63) and mzMatch software (64) in the R environment. A list of putatively annotated metabolites was generated and the abundances of all [^13^C]-isotopologues for these were obtained using the software mzMatch-ISO (65). Metabolomics data have been deposited to the EMBL-EBI MetaboLights database (DOI: 10.1093/nar/gkad1045, PMID:37971328) with the identifier MTBLS10738.

### 2.6. Expression and activity of threonine metabolism genes in *E. multilocularis* metacestode vesicles *in vitro*

#### 2.6.1. Genes of threonine metabolism in *E. multilocularis*

The following protein sequences of threonine metabolism were blasted against the protein database of *E. multilocularis* PRJEB122 (39) via WormBase ParaSite (https://parasite.wormbase.org, assessed on 10/17/2023): TD, TDH, KBL and TA from reference organisms *Caenorhabditis elegans, Danio rerio, Drosophila melanogaster*, *Homo sapiens* and *Mus musculus* (if present as functional proteins, see S1 Table). Human *ta* and *tdh* were excluded since they are pseudogenes (56,66) and *td* is not present in *Da. rerio*. Top hits within the protein database of *E. multilocularis* were then blasted reciprocally against the NCBI non-redundant protein database of the reference organisms (https://blast.ncbi.nlm.nih.gov/Blast.cgi). We performed the same approach to identify TDH sequences in the closely related parasites *E. granulosus s.s.* with the protein database PRJEB121 and *E. canadensis* with the protein database PRJEB8992.

#### 2.6.2. Preparation of *E. multilocularis* metacestode vesicles and *in vivo* grown metacestodes for assessment of threonine metabolism gene expression

In order to study whether threonine metabolism of *in vitro* cultured metacestode vesicles of isolate H95 under axenic culture conditions in a microaerobic atmosphere was similar to previously published data from isolate G8065 (39), we analyzed gene expression via quantitative real-time PCR for the four genes, *emtd* (EmuJ_001093200), *emtdh* (EmuJ_000511900), *emkbl* (EmuJ_000107200) and the house-keeping gene *ezrin/radixin/moesin-like protein* (*emelp*) (EmuJ_000485800) (67,68). Five months old *E. multilocularis* metacestode vesicles were purified and changed to an axenic culture system without RH cells as described by others (58) and incubated under humid microaerobic atmosphere for two days. Metacestode vesicles were mixed with three volumes of A6 medium (see 2.4.2 but using DMEM with 4.5 g/L of glucose), and 4 mL were distributed to 6-well plates. Each condition was set up in four biological replica and two independent experiments were conducted. Metacestode vesicles were incubated under humid, microaerobic atmosphere for three days. Metacestode vesicles were washed three times in PBS, destroyed with a pipette and again washed three times in PBS with a centrifugation step (600 x g, 3 minutes, 4°C) after each washing step. The metacestode VT was taken up in 1.8 mL TRI Reagent®, shaken at 1,400 RPM in an Eppendorf® Thermomixer Compact (Vaudaux Eppendorf, Schönenbuch, Switzerland) for 15 minutes at RT and frozen to -20°C until further use.

Mice were intraperitoneally injected with *E. multilocularis* metacestode tissue for routine strain maintenance (see also 2.2). Metacestode tissue from four individual mice was washed in PBS, mechanically disrupted and homogenized in 1.8 mL TRI Reagent® in a 2 mL screw cap tube with a five mm glass bead in a FastPrep-24TM Classic homogenizer (MP Biomedicals, Illkirch-Graffenstaden, France) with five cycles of 4 m/s for 20 seconds. Then, samples were shaken at 1,400 RPM in an Eppendorf® Thermomixer Compact for 15 minutes at RT and centrifuged at (12,000 x g, 10 minutes, 4°C). The supernatant was transferred to a new tube and samples were frozen to -20°C until further use.

#### 2.6.3. Isolation of RNA from *E. multilocularis* metacestode vesicles and metacestode cysts

RNA from *E. multilocularis* metacestode vesicles and metacestode cyst tissue was isolated via a phenol/chloroform extraction as described by others (69) with a few modifications. In short, 0.2 volumes of chloroform were added per mL TRI Reagent® to each sample, they were incubated at RT for three minutes and subsequently centrifuged (12,000 x g, 15 minutes, 4°C). The aqueous phase was taken, and samples were pipetted on parafilm to remove residual chloroform, mixed with 1 mL isopropanol and incubated at RT for 10 minutes. After centrifugation (12,000 x g, 15 minutes, 4°C) the pellets were washed once in 75% ethanol and centrifuged (7,500 x g, 5 minutes, 4°C). The pellets were air-dried, and DNA digestion was performed with the Direct-zol RNA Miniprep Kit from Zymo Research (Lucerna-Chem AG, Lucerne, Switzerland). The RNA was resuspended in 87.5 µL RNAse-free water, mixed with 20 µL DNA Digestion Buffer and 5 µL DNAse I. Samples were incubated at RT for 10 minutes and then 0.1 volumes of 3 M Diethyl pyrocarbonate-treated sodium acetate, pH 5.2, and 2.5 volumes of 100% ethanol were added. Samples were incubated at -80°C for 1.5 hours. The samples were centrifuged (16,000 x g, 10 minutes, 4°C) and the pellet washed once in 75% ethanol and centrifuged (7,500 x g, 5 minutes, 4°C). Pellets were air-dried, resuspended in RNAse-free water and concentrations were measured via a NanoDrop™ One/OneC Microvolume UV-Vis Spectrophotometer (Thermo Fisher Scientific, Reinach, Switzerland). 1 µg of RNA was reverse transcribed via the GoScript™ Reverse Transcription System (Promega, Dübendorf, Switzerland) in a final volume of 20 µL.

#### 2.6.4. Quantitative real-time PCR of *E. multilocularis* RNA samples

Gene expression was analyzed via specific, intron-flanking primers, which are shown in in S2 Table. Quantitative real-time PCRs were performed on a CFX Opus 96 Real-Time PCR System (Biorad). Primer efficiency was calculated by performing RT-PCR reactions using 1 µL of cDNA, and 1 µL of four subsequent 1:4 dilutions as template, except for *emtd* for which 1:2 dilutions were made, due to the expected lower transcription (39). Gene expression was analyzed in technical duplicates for each biological quadruplicate and calculated relative to *emelp*. Data is shown as relative fold-change. Statistical analyses were performed using multiple two-tailed students t-tests with equal variance and subsequent Bonferroni-correction in R. Bonferroni-corrected *p*-values of *p*<0.05 were considered to be significant.

#### 2.6.5. Enzymatic activity of EmTDH in crude extracts of *E. multilocularis* metacestode vesicles

For the preparation of crude extracts of *E. multilocularis* metacestode vesicles, parasite vesicles grown for at least 6 months *in vitro* were ruptured by pipette and the pellet was washed three times in PBS. The pellet was taken up in a lysis buffer (80 mM Tris HCl pH 8.4 supplemented with 1% Triton X-100, 1% Halt™ Protease Inhibitor Cocktail (Thermo Fisher Scientific) and 1% EDTA) in a volume where one mL of buffer corresponded to 10 mL pure, intact metacestode vesicles. The protein amount was determined by the Pierce™ BCA Protein Assay Kit and resulted in 8 mg/mL.

We established an assay for the characterization of enzymatic activity of EmTDH within crude extract of *E. multilocularis* metacestode vesicles. The TDH assay was adapted for *E. multilocularis* metacestode vesicles using a protocol for *E. coli* TDH as a basis (70). The enzymatic assay buffer consisted of of 80 mM Tris HCl (pH 8.4) and 10 mM NAD. We incubated 80 µg protein crude extract of *E. multilocularis* metacestode vesicles per well with various concentrations of L-threonine and D-threonine (0, 1, 2, 4, 8, 16 mM) in triplicates. Enzyme blanks were included. The reaction was measured at 37°C via an increase in absorbance at 340 nm on a HIDEX Sense microplate reader (Hidex, Turku, Finland). Enzyme blanks were subtracted from the results and shown are mean values and SDs.

### 2.7. Inhibition of *E. multilocularis* threonine metabolism by TDH inhibitors

#### 2.7.1. TDH assay with recombinantly expressed EmTDH and MmTDH

The sequence of *emtdh* was obtained from WormBase ParaSite (https://parasite.wormbase.org) via the accession number EmuJ_000511900 and the sequence of *mmtdh* was obtained from the National Library of Medicine (https://www.ncbi.nlm.nih.gov) via the accession number ENSMUST00000022522.15. Both sequences, as well as the detailed cloning process with images is shown in S5 File.

Briefly, *emtdh* (EmuJ_000511900) was amplified from cDNA of *in vitro* grown *E. multilocularis* metacestode vesicles without the predicted mitochondrial target sequence as a 979 bp sequence. The amplification of *mmtdh* (ENSMUST00000022522.15) was difficult, due to low expression in tissue of adult mice (60) and was therefore ordered as a 1,007 bp fragment from LubioScience (Lucerne, Switzerland) without the predicted mitochondrial target sequence, and amplified with a resulting size of 991 bp. Both *emtdh* and *mmtdh* were cloned into the pET151/D-TOPO® vector using the Champion™ pET151 Directional TOPO™ Expression Kit (Fisher Scientific AG, Reinach, Switzerland). Clones were picked and tested via colony PCRs for the expected fragment size and correct orientation with the vector-specific forward primer and an insert-specific reverse primer. Plasmid DNA was isolated from positive clones using the ZymoPURE™ Plasmid Miniprep Kit (Zymo Research, Irvine, USA) and Sanger sequencing was conducted at Microsynth AG (Balgach, Switzerland). Sequences were compared in BioEdit (71) to their respective reference sequence and correct plasmids were used to transform *E. coli* BL21 for recombinant expression of His-tagged EmTDH as a 365 amino acids protein and His-tagged MmTDH as a 361 amino acid protein. RecEmTDH and recMmTDH were purified via the Macherey-Nagel™ Protino™ Ni-TED-IDA 1000 Kit (Fisher Scientific, Schwerte, Germany) according to the manufacturer’s protocol and eluates were checked on a 12% sodium dodecyl sulfate polyacrylamide gel. Correct protein size of 41.4 kDa for recEmTDH and 40.3 kDa for recMmTDH was confirmed by western blot using a mouse monoclonal anti-His tag antibody and an anti-mouse IgG (h+l) ap conjugate as secondary antibody (Promega, Dübendorf, Switzerland). Finally, protein concentration of the eluates was determined via BCA assay using the Pierce™ BCA Protein Assay Kit.

#### 2.7.2. Inhibition of recEmTDH and recMmTDH

We first tested the activity of recEmTDH and recMmTDH via the established TDH assay (see 2.6.5) using a concentration series of L-threonine and D-threonine (0, 1, 2, 4, 8, 16 mM), 10 mM NAD and 15 nM of either recEmTDH or recMmTDH per well, each condition in technical triplicates. Enzyme blanks (without recEmTDH or recMmTDH) and substrate blanks (without addition of L-threonine) were included. Enzymatic activity was measured at 37°C via an increase in absorbance at 340 nm on a HIDEX Sense microplate reader. Substrate blanks were subtracted from the enzymatic reaction wells.

For the subsequent inhibition experiments, the assay buffer was supplemented with 2 mM L-threonine, 3 mM NAD and 15 nM recEmTDH, or 15 nM recMmTDH, per well of a 96-well plate. Several published inhibitors of threonine metabolism were tested on recEmTDH and recMmTDH in the established TDH assay, namely disulfiram, myricetin, quercetin, seven quinazoline carboxamides (QC) and sanguinarine (Adjogatse, 2015; Alexander et al., 2011; Cross et al., 1975). The inhibitors were added at 20 µM and enzyme activity was normalized to respective DMSO controls. Significant inhibition was assessed via multiple two-sample one-tailed students t-tests assuming equal variance and subsequent Bonferroni-correction in R. Inhibitors were considered active when reduction of enzyme activity was significant according to Bonferroni-corrected *p*-values of *p*<0.05. Active inhibitors against recEmTDH were tested in concentration series from 111.1 µM to 0.02 µM in 1:3 serial dilutions in triplicates against both, recEmTDH and recMmTDH. IC_50_-values were calculated for each of the three independent experiments using an IC_50_-calculator in R (75). Fold changes between IC_50_-values of inhibitors against recEmTDH and recMmTDH were calculated and significant differences were calculated via two-sample two-tailed students t-test with equal variance and subsequent Bonferroni-correction in R. Bonferroni-correct *p*-values of *p*<0.05 were considered to be significant.

### 2.8. Assessment of EmTDH inhibitor activity against *E. multilocularis in vitro*

#### 2.8.1. Phosphoglucose isomerase (PGI) assay on *E. multilocularis* metacestode vesicles

Damage marker release assays, based on the marker PGI, were performed as described previously (32) with the modifications published recently (30). In short, two-to three-months-old metacestode vesicles were purified with 2% sucrose and several washing steps in PBS. Metacestode vesicles were mixed with two volumes of high glucose DMEM without phenol red containing penicillin (100 U/mL) and streptomycin (100 µg/mL). 1 mL of the metacestode vesicle-medium mix was distributed to a 48-well plate (Huberlab, Aesch, Switzerland) and disulfiram, myricetin and sanguinarine were added to final concentrations of 20 µM in triplicates. The respective amount of DMSO was used as a negative control and 0.1% Triton X-100 was used as a positive control. The plate was incubated at 37°C under a humid, microaerobic atmosphere and pictures and supernatant samples were taken after five days. Supernatant samples were measured on a HIDEX Sense microplate reader. The corresponding values of the DMSO controls were subtracted from the values of the compounds and then PGI activity was calculated relative to 0.1% Triton X-100. Given are mean values and SD. Significant differences of compound-treated metacestode vesicles compared to the DMSO control were calculated via two-sample two-tailed students t-test with equal variance and subsequent Bonferroni-correction in R. Bonferroni-corrected *p*-values of *p*<0.05 were considered to be significant.

IC_50_-calculations were carried out for sanguinarine, testing this compound on metacestode vesicles at concentrations from 40 to 1.25 µM in 1:2 serial dilutions in triplicates. IC_50_-values were calculated using an IC_50_-calculator in R for each of three independent experiments and then the mean value and SD were calculated.

#### 2.8.2. GL cell viability assay

Cell viability assays with *E. multilocularis* GL cells were carried out as described recently (30). In short, 15 AU of GL cells (see 2.4.3 for extraction of GL cells) were distributed into wells of a black 384-well plate in a volume of 12.5 µL. In another 12.5 µL, disulfiram, myricetin and sanguinarine were added to final concentrations of 20 µM, respective amounts of DMSO and Triton X-100 (0.1%) were included as controls. For overview screening, each drug was tested in quadruplicates. Cells were incubated at 37°C under a humid, microaerobic atmosphere for five days. Pictures were taken and to each well, 25 µL of CellTiter-Glo containing 1% Triton X-100 was added and cell aggregates were disrupted by pipetting. Luminescence was recorded on a HIDEX Sense microplate reader and mean values and SD were calculated. Significant differences of compound-treated GL cell cultures compared to the DMSO control were calculated via two-sample two-tailed students t-test with equal variance and subsequent Bonferroni-correction in R. Bonferroni-corrected *p*-values of *p*<0.05 were considered to be significant. Two independent experiments were performed for the overview screen. In order to calculate IC_50_-values for sanguinarine, this compound was added in a 1:2 dilution series from 40 to 0.08 µM to the GL cells in quadruplicates in three independent experiments. Activity was assessed as described for the overview screening and IC_50_-values were calculated using an IC_50_-calculator in R and mean and SD are given.

#### 2.8.3. Metacestode vesicle viability assay

Viability of metacestode vesicles was assessed for sanguinarine as described recently (30) with the difference that measurements were carried out in white 96-well plates. Metacestode vesicle viability was measured after 12 days using the same setups as for the PGI assay (see 2.8.1). In short, Triton X-100 was added to each well to a final concentration of 0.1% and then metacestode vesicles were mechanically broken with a pipette. 50 µL supernatant were transferred to a white 96-well plate and mixed with 50 µL of CellTiter-Glo (Promega, Dübendorf, Switzerland). Measurements were performed on a HIDEX Sense microplate reader and viability was set relative to the respective DMSO control. For each of the three independent experiments, IC_50_-values were calculated using an IC_50_-calculator in R and then the mean value and SD were calculated.

#### 2.8.4. Cytotoxicity assays with mammalian cells

For sanguinarine, cytotoxicity assays with *M. musculus* Hepa 1-6 cells, *Rattus norvegicus* RH cells and *H. sapiens* HFF cells were performed by alamar blue assay as described previously (76) with the modifications published recently (77). Three independent experiments were performed, and IC_50_-values were calculated for each of cell type using an IC_50_-calculator in R and mean values and SDs are given.

## 3. Results

### 3.1. Threonine metabolism promotes *E. multilocularis* metacestode vesicle growth and development *in vitro*

To study the effect of threonine on *E. multilocularis* growth *in vitro*, we developed a metacestode vesicle growth assay using an automated and a semi-automated script in ImageJ to precisely and objectively follow the growth of single metacestode vesicles over time. The scripts were validated with 150 photos of 50 individual metacestode vesicles being photographed each three times. The mean vesicle diameters and SDs are shown in S1 Fig. There was no significant difference between measurement of metacestode vesicle diameters via the automated script, the semi-automated script, or the manual measurement performed in ImageJ. The mean internal diameter variance between the three images of each metacestode vesicle was 1.3% for the manual measurements and 1.2% for both the automated and the semi-automated scripts. Due to the general low variance between the photos, the images of the metacestode vesicle growth assays were analyzed with the automated script and in case vesicles were not accurately detected, images were analyzed with the semi-automated script.

In a preliminary experiment the most stuitable L-threonine concentration for a growth assay was assessed. Under the described culture conditions, addition of 4 mM of L-threonine led to the highest reduction of L-threonine in the culture medium (S2 Fig). Thus, 4 mM L-threonine was used as a maximal concentration to be tested subsequently.

For experiment one, *E. multilocularis* metacestode vesicles were cultured *in vitro* in medium supplemented with L-threonine at 1, 2 and 4 mM, or 4 mM D-threonine, (Fig 2A). Of a total of 384 metacestode vesicles analyzed, seven (1.8%) collapsed within the six weeks of the experiment and thus these 14 images from week 0 and week 6 were excluded from the analysis. Of the remaining 754 images, 696 images (92.3%) were detected correctly by the automated script. The 55 images (7.3%) in which the metacestode vesicles were not accurately detected were processed with the semi-automated script. Over the time course of six weeks, metacestode vesicles supplemented with water as a control grew 1.9 mm ± 0.6 mm. The growth was significantly increased when metacestode vesicles were cultured with 1 mM L-threonine (2.5 mm ± 0.6 mm, *p*=0.025), 2 mM L-threonine (2.8 mm ± 0.8 mm, *p*=8e-4) and 4 mM L-threonine (3.0 mm ± 0.9 mm, *p*=1e-4). Supplementation with 4 mM D-threonine led to a growth of 2.0 mm ± 0.8 mm, which was not significantly different from the control. Thus, L-but not D-threonine, led to a significant stimulation of *E. multilocularis* metacestode vesicle growth *in vitro.* The experiment was repeated once independently, and we obtained similar results with significant increase of metacestode vesicle growth upon increasing concentrations of L-threonine but not D-threonine (S3 Fig).

**Fig 2:**
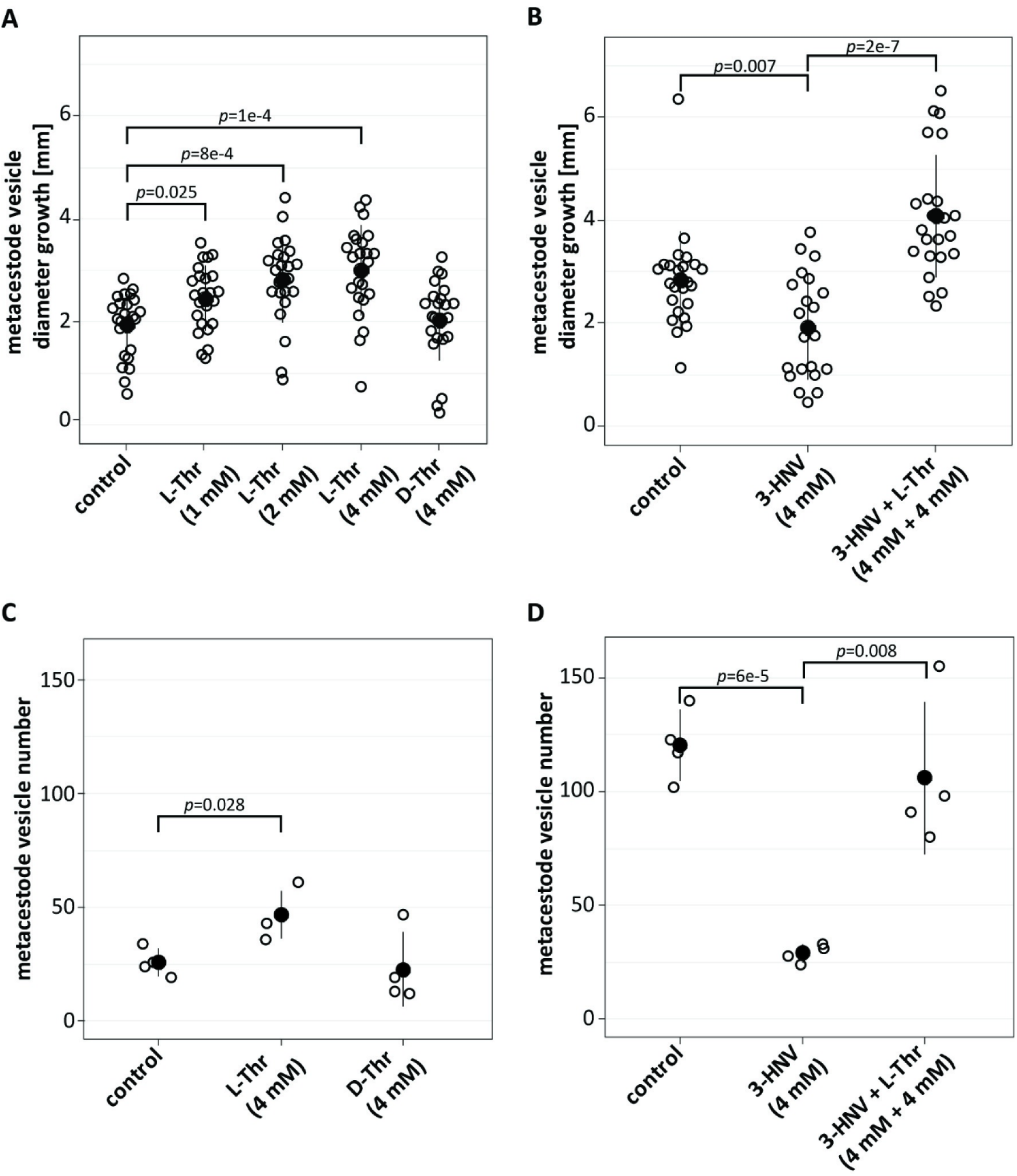
Threonine stimulates *E. multilocularis* metacestode vesicle growth and development *in vitro*. A, *E. multilocularis* metacestode vesicles were cultured with various amounts of L-threonine (1 mM, n=24; 2 mM, n=23; 4 mM, n=23), D-threonine (4 mM, n=23), or water as a control (n=24). B, *E. multilocularis* metacestode vesicles were cultured in the presence of 3-HNV (4 mM, n=21), a combination of 3-HNV and L-threonine (each at 4 mM, n=23), or water (n=24). Metacestode vesicle growth in A and B was calculated via an automated script and growth in mm and Bonferroni-corrected *p-*values as compared to the water controls are displayed. C, *E. multilocularis* GL cell cultures were supplemented with L-threonine (4 mM, n=4), D-threonine (4 mM, n=4), or water as a control (n=4). D, *E. multilocularis* GL cells were cultured with 3-HNV (4 mM, n=4), a combination of 3-HNV and L-threonine (both at 4 mM, n=4), or water as a control (n=4). Newly formed metacestode vesicles in C and D were counted manually in a blinded manner and the mean values, SD and Bonferroni-corrected *p-*values are shown. Abbreviations: Thr = threonine, 3-HNV = 3-hydroxynorvaline. Two independent experiments were performed for each setup, and the results of one representative experiment is shown, while the results of the second experiment are provided in S3 Fig.

In a second experiment (Fig 2B), *E. multilocularis* metacestode vesicles grew 2.8 mm ± 0.9 mm within the six weeks of this experiment and this growth was significantly reduced upon incubation with 4 mM 3-HNV (1.8 mm ± 1.1 mm, *p*=0.007). The combined incubation of 4 mM 3-HNV and 4 mM L-threonine compensated for this reduction significantly (4.1 mm ± 1.2 mm, *p*=2e-7). Upon independent repetition of the experiment, we obtained similar results with significant reduction in growth for metacestode vesicles treated with 4 mM 3-HNV and that was significantly counteracted upon treatment with a combination of 4 mM 3-HNV and 4 mM L-threonine (S3 Fig).

We then assessed the effects of threonine on the formation of new metacestode vesicles from GL cells. In the control culture supplemented with water 26 ± 5 metacestode vesicles were formed within two weeks (Fig 2C). This formation was significantly increased upon addition of 4 mM L-threonine (47 ± 9 metacestode vesicles, *p*=0.028), but not with the addition of D-threonine (23 ± 14 metacestode vesicles). We repeated this experiment once independently and obtained a similar trend, but not a significant difference between the control and L-threonine (see S3 Fig).

We then assessed the effect of 3-HNV with and without L-threonine accordingly on the metacestode vesicle formation rate (Fig 2D). While supplementation of water resulted in the formation 121 ± 14 metacestode vesicles from GL cell cultures, we observed a significant reduction in GL cell cultures incubated with 4 mM 3-HNV (29 ± 3 metacestode vesicles, *p*=6e-5). As observed in the metacestode vesicle growth assay, the combined supplementation of 4 mM 3-HNV and 4 mM L-threonine significantly counteracted this reduction (106 ± 26 metacestode vesicles, *p*=0.008). This experiment was repeated once independently, and we obtained similar and significant results (S3 Fig).

### 3.2. *E. multilocularis* metacestode vesicles metabolize threonine to glycine via EmTDH

In order to identify relevant pathways through which threonine is metabolized by *E. multilocularis*, we performed a flux experiment with [U-^13^C]-L-threonine and detected labeled metabolites by LC-MS in different fractions of metacestodes (VT, VF, CM and VM). Labeling patterns are visualized in Fig 3 and individual values of fractional enrichment are shown in S3 Table. We detected [U-^13^C]-L-threonine in VT, VF, CM and VM (93 ± 0.4%, 94.2 ± 0.1%, 96.1 ± 0% and 95.8 ± 0.1%, labeling, respectively) indicating an uptake of L-threonine by *E. multilocularis* metacestode vesicles and transportation to the VF. The direct metabolic product of TD-mediated threonine catabolism, α-ketobutyrate, was not detected in any of the samples. The direct product of TDH-mediated threonine catabolism, 2-amino-3-ketobutyrate, an unstable product, was not detected either. However, we detected [U-^13^C]-aminoacetone in VM (64.6 ± 5.6%), which is generated upon spontaneous decarboxylation of 2-amino-3-ketobutyrate (46). We further detected [U-^13^C]-glycine in VT, VF and VM (42.7 ± 2.4%, 30.1 ± 2.3% and 28 ± 3.3%, respectively), but only minor traces in CM (0.2 ± 0.2%) indicating that threonine metabolism fed into glycine production. Besides glycine, acetyl-coenzyme A is also generated via KBL-mediated metabolization of 2-amino-3-ketobutyrate. We detected various TCA cycle intermediates, namely citrate, its downstream metabolite α-ketoglutarate and its derivative 2-hydroxyglutarate, succinate, malate, and the transamination product of oxaloacetate, L-aspartate. None of these metabolites were found to be [^13^C]-labeled. Thus, our results do not suggest the use of L-threonine metabolites within the TCA cycle in *E. multilocularis* metacestode vesicles. Further, we detected [^13^C_2_]-glutathione in VT samples (6.6 ± 0.5%), which indicates that L-threonine-derived glycine fed into the biosynthesis of glutathione. Additionally, we found [^13^C_2_]-L-glutamate in VT with 2 ± 0.2% and some traces in VF and VM (0.3 ± 0.3% and 0.1 ± 0.1%, respectively), while CM was completely unlabeled, which indicates a L-threonine-derived formation of L-glutamate by *E. multilocularis in vitro*. We detected palmitate and propionate in our samples, but also in the blank samples and neither of these metabolites contained [^13^C]. Thus, the data was not exploited.

**Fig 3.**
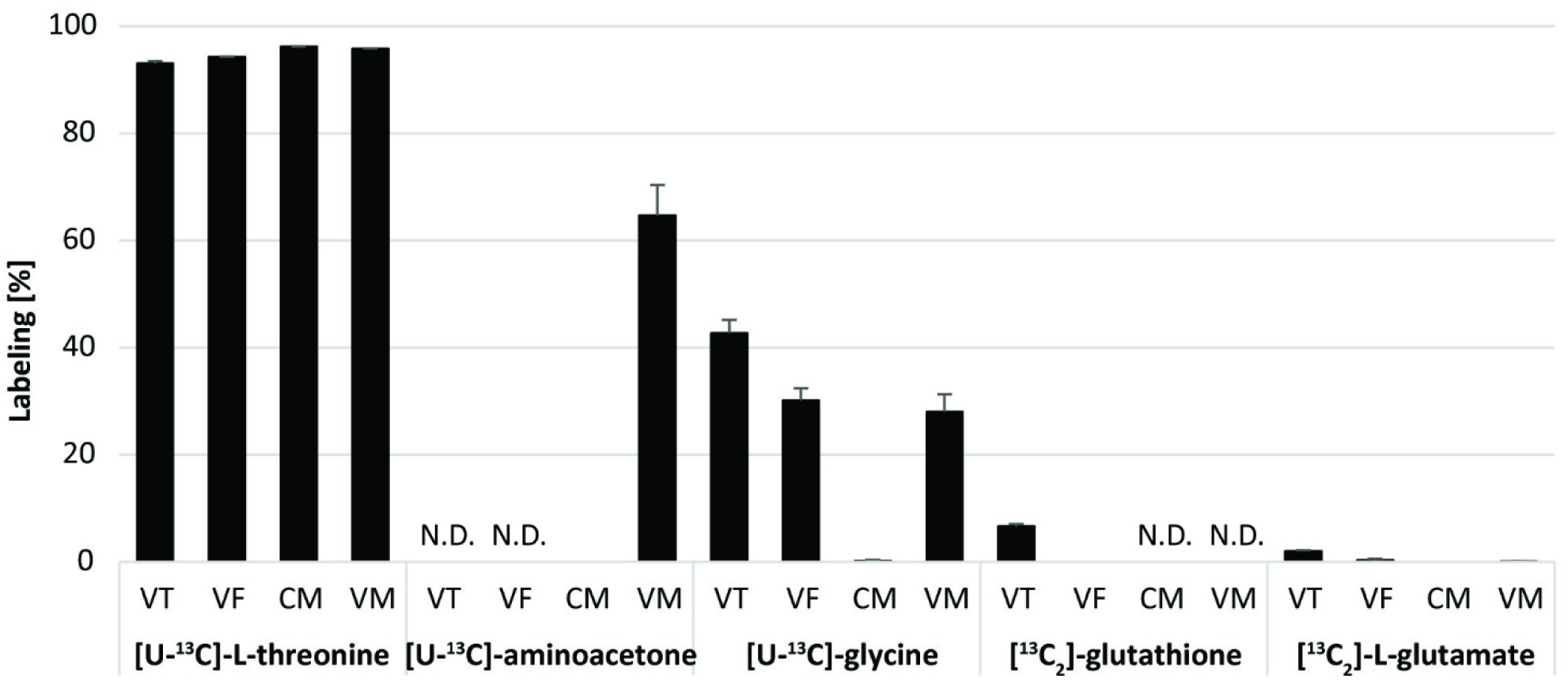
Results of the [U-^13^C]-L-threonine flux assay. *E. multilocularis* metacestode vesicles were cultured with 5 mM [U-^13^C]-L-threonine or unlabeled L-threonine for 24 hours at 37°C under a humid, microaerobic atmosphere (n=4). Labeled metabolites were detected in vesicle tissue (VT), vesicle fluid (VF), or medium without and with metacestode vesicles (CM and VM) via LC-MS. Shown are [^13^C]-labeled metabolites with average values and SD in %. Individual values of fractional enrichment of all metabolites are shown in S3 Table. N.D.: not detected.

### 3.3. Genes for threonine catabolism are present and expressed in the *E. multilocularis* metacestode stage

In order to search for threonine metabolism genes within *E. multilocularis,* we blasted protein sequences of TD, TDH, KBL and TA from various reference organisms (*C. elegans, Da. rerio, Dr. melanogaster*, *H. sapiens* and *M. musculus*) against the protein database of *E. multilocularis*. TD sequences from *H. sapiens* and *M. musculus* both resulted in one hit (EmuJ_001093200). TD sequences from *C. elegans* and *Dr. melanogaster* did not identify hits. TDH sequences from *C. elegans, Da. rerio*, *Dr. melanogaster* and *M. musculus* found only one hit (EmuJ_000511900). KBL sequences of *C. elegans*, *Da. rerio, Dr. melanogaster*, *H. sapiens* and *M. musculus* found several hits, but EmuJ_000107200 corresponded to the lowest E-values in all blasts. None of the tested TA sequences from *C. elegans, Da. rerio*, *Dr. melanogaster* and *M. musculus* resulted in any hits within the protein database of *E. multilocularis*. Reciprocal blasts of EmuJ_001093200, EmuJ_000511900 and EmuJ_000107200 were performed against protein databases of *C. elegans, Da. rerio*, *Dr. melanogaster, H. sapiens* and *M. musculus*. Reciprocal blasts confirmed EmuJ_001093200 as EmTD, EmuJ_000511900 as EmTDH and EmuJ_000107200 as EmKBL. Accession numbers and results of the BLASTP and reciprocal BLASTP with all sequences producing significant alignments and E-values can be found in S4 Table, S5 Table and S6 Table. We further performed BLASTP with TDH sequences from the same reference organisms against the protein databases of the closely related parasites *E. granulosus s.s.* and *E. canadensis* (S7 Table and S8 Table) and subsequent reciprocal blasts confirmed EgrG_000511900 as EgTDH and EcG7_08078 as EcTDH (S9 Table and S10 Table).

We then analyzed gene expression of *emtd*, *emtdh* and *emkbl* relative to the house keeping gene *emelp in vitro* under axenic culture conditions with a microaerobic atmosphere (Fig 4A) and found that both *emtdh* (relative expression of 0.55 ± 0.04) and *emkbl* (relative expression of 1.32 ± 0.26) were significantly higher expressed than *emtd* (relative expression of 0.003 ± 0.0001, *p*=5e-7 and *p*=3e-4, respectively). We repeated the experiment once and came to the same conclusion (S4 Fig). We also analyzed the expression of these genes in metacestode tissue obtained from experimentally infected mice and again noted significantly higher expression of *emtdh* (relative expression of 0.61 ± 0.09) and *emkbl* (relative expression of 0.33 ± 0.05) compared to *emtd* (relative expression of 0.0004 ± 0.0002, *p*=4e-5 and *p*=3e-5, respectively) (Fig 4B).

**Fig 4.**
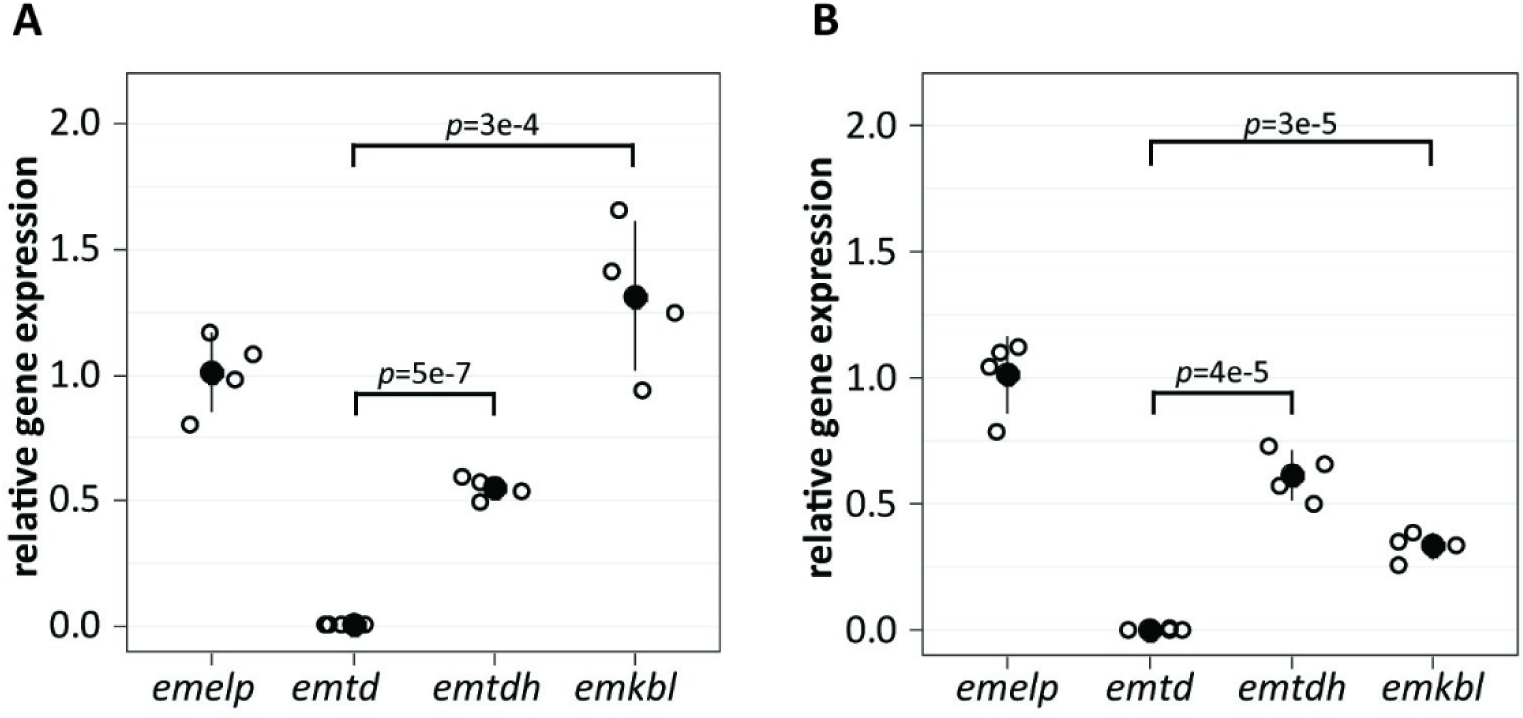
Relative expression of threonine metabolism genes in the *E. multilocularis* metacestode stage. Relative expression was analyzed in *E. multilocularis* metacestode vesicles cultured *in vitro* (A) or in *E. multilocularis* metacestode tissue obtained from experimentally infected mice (B), (n=4 for both). Q-RT-PCRs were performed in technical duplicates for each sample and gene expression was calculated relative to the housekeeping gene *emelp*. Shown are Bonferroni-corrected *p-*values. Two independent experiments were performed for A and the other experiment is shown in S4 Fig.

### 3.4. The protein EmTDH is expressed and enzymatically active in the *E. multilocularis* metacestode stage

In a recent study, EmTDH and EmKBL were found via non-targeted proteomics samples of VF and VT of *in vitro* cultured metacestode vesicles, as well as in VF of *in vivo* grown metacestodes obtained from experimentally infected mice (17). Interestingly, EmTD was not detected in these samples.

To assess whether *emtdh* is translated into an enzymatically active protein in *E. multilocularis in vitro*, we measured enzymatic activity of EmTDH in crude extracts of *in vitro* cultured metacestode vesicles. We observed a dose-dependent increase in absorbance, indicating enzyme activity upon addition of the substrate L-threonine, but not D-threonine (Fig 5A). The negative value calculated for the enzymatic reaction without addition of L-threonine or D-threonine was caused by a decrease in absorbance in these wells.

**Fig 5.**
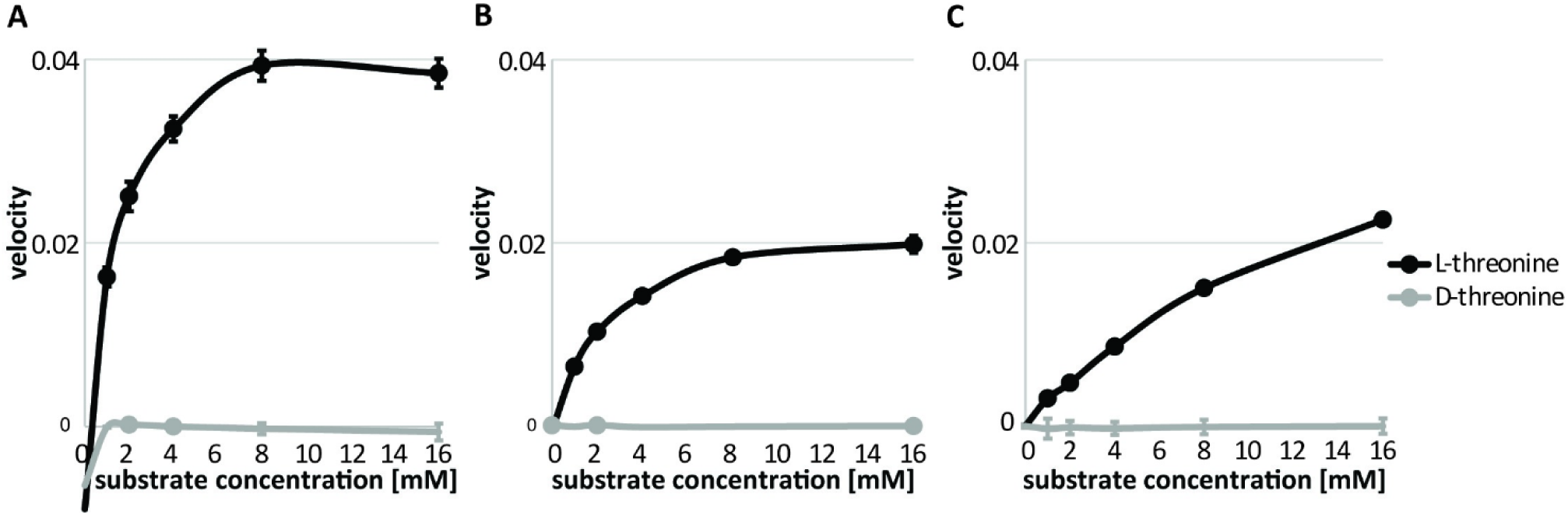
Enzymatic assays to assess TDH activity. Enzymatic assays were carried out via measuring the increase in absorbance at 340 nm at 37°C. Concentration series of L-threonine or D-threonine from 0 to 16 mM were tested with a NAD concentration of 10 mM (n=3). Shown are TDH assays with 80 µg protein of crude extracts of *E. multilocularis* metacestode vesicles per well (A), 15 nM recEmTDH per well (B) or 15 nM recMmTDH per well (C). Shown are mean values and SD.

We recombinantly expressed EmTDH (S5 File) and also here detected L-threonine-dependent activity (Fig 5B). As a control protein, we recombinantly expressed MmTDH (S5 File) and this protein was also enzymatically active (Fig 5C).

### 3.5. EmTDH can be inhibited by repurposed TDH inhibitors

We applied the enzymatic assays for recEmTDH and recMmTDH to test the activities of various previously published TDH inhibitors (disulfiram, myricetin, quercetin, seven QCs and sanguinarine) (72–74) (Fig 6).

**Fig 6.**
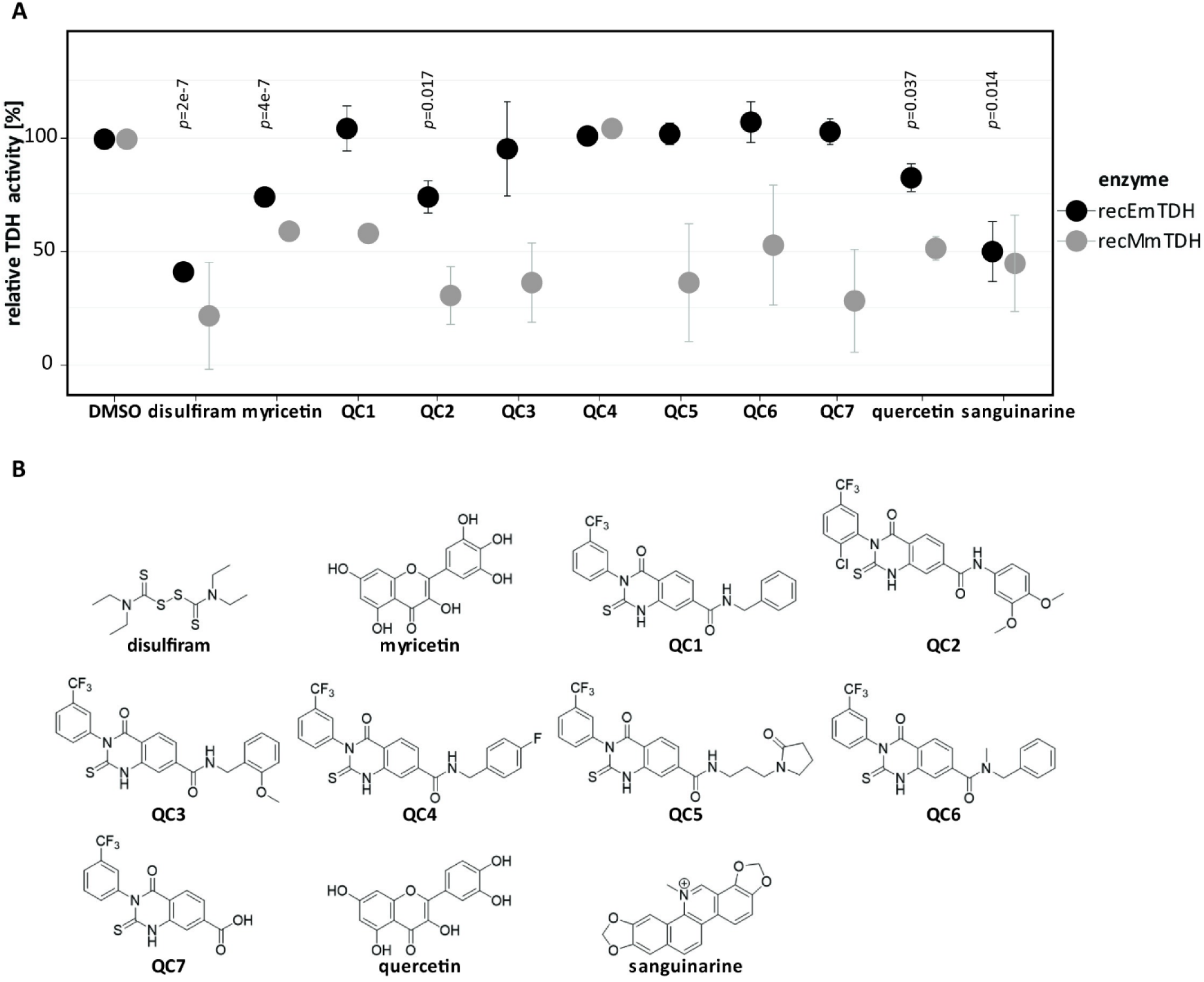
Effect of various potential TDH inhibitors on recEmTDH and recMmTDH. A, TDH activity was assessed as in Fig 5. Compounds were tested at 20 µM (n=3) and shown are mean values and SDs of three independent experiments. Bonferroni-corrected *p-*values are shown for relative activities of recEmTDH treated with the different compounds compared to their respective DMSO control. Bonferroni-corrected *p*-values for relative activities of recMmTDH are shown in S11 Table. B, structures of potential TDH inhibitors tested against recEmTDH and recMmTDH in this study. The synthesis for QC1 to QC7 is described in S1 File.

In relation to the DMSO control, several compounds, when applied at 20 µM, significantly reduced the activity of recEmTDH. Enzymatic activity was reduced to 41.5 ± 0.9% for disulfiram (*p*=2e-7), 74.7 ± 0.4% for myricetin (*p*=4e-7), 74.3 ± 5.7% for QC2 (*p*=0.017) 82.4 ± 4.8% for quercetin (*p*=0.037) and 50.1 ± 10.6% (*p*=0.014) for sanguinarine. However, many compounds also inhibited recMmTDH activity. Thus, we decided to determine the IC_50_-values of the five most active recEmTDH inhibitors disulfiram, myricetin, QC2, quercetin and sanguinarine on recEmTDH and recMmTDH (Table 1).

**Table 1.** Half-maximal inhibitory concentrations of TDH inhibitors against recEmTDH and recMmTDH. Activity of recEmTDH and recMmTDH was assessed upon incubation with drugs at concentrations from 111 to 0.02 µM in 1:3 serial dilutions (n=3). TDH activity was set relative to the DMSO control and dose-dependent response curves were used to calculate IC_50_-values. Shown are mean values and SDs of three independent experiments for both enzymes and the fold-change of IC_50_-values between recEmTDH and recMmTDH with significance given as ns (not significant) or as Bonferroni-corrected *p*-value.

| compound | recEmTDH IC<sub>50</sub> | recMmTDH IC<sub>50</sub> | fold change |
| --- | --- | --- | --- |
| disulfiram | $15.2 \pm 5.2 \mu\text{M}$ | $1.4 \pm 0.5 \mu\text{M}$ | 11.1 (ns) |
| myricetin | $9.1 \pm 1.8 \mu\text{M}$ | $16.6 \pm 14.4 \mu\text{M}$ | 0.5 (ns) |
| QC2 | $36.3 \pm 13.1 \mu\text{M}$ | $7.2 \pm 1.2 \mu\text{M}$ | 7.6 (ns) |
| quercetin | $30.9 \pm 8.8 \mu\text{M}$ | $21.6 \pm 10.1 \mu\text{M}$ | 1.4 (ns) |
| sanguinarine | $5.6 \pm 0.9 \mu\text{M}$ | $1.8 \pm 0.6 \mu\text{M}$ | 3.2 ($p=0.037$) |

Sanguinarine was the most active recEmTDH inhibitor with an IC_50_-value of 5.6 ± 0.9 µM, followed by myricetin with an IC_50_-value of 9.1 ± 1.8 µM and disulfiram with an IC_50_-value of 15.2 ± 5.2 µM. Quercetin and QC2 were less active with IC_50_-values of 30.9 ± 8.8 µM and 36.3 ± 13.1 µM, respectively. Most of these inhibitors were more active against recMmTDH with IC_50_-values of 1.8 ± 0.6 µM (fold-change of 3.2) for sanguinarine, 1.4 ± 0.5 µM (fold-change of 11.1) for disulfiram, 21.6 ± 10.1 µM (fold-change of 1.4) for quercetin and 7.2 ± 1.2 µM (fold-change of 7.6) for QC2. Only myricetin showed a lower activity against recMmTDH with an IC_50_-value of 16.6 ± 14.4 µM and thus a fold-change of 0.5.

### 3.6 Sanguinarine is active against *E. multilocularis in vitro*

The three most active recEmTDH inhibitors disulfiram, myricetin and sanguinarine were not specifically active against recEmTDH over recMmTDH. However, since human *tdh* is a pseudogene (56) specificity of potential inhibitors on EmTDH over other TDH proteins could be less relevant. Thus, we tested these three inhibitors on *E. multilocularis* metacestode vesicles via damage marker release assay (PGI assay) and GL cell viability assay (Fig 7). The negative control DMSO did not affect the physical appearance of metacestode vesicles after five days (Fig 7A), while Triton X-100 led to maximum physical damage and PGI release (100 ± 8.8 %, *p*=1.7e-4). Disulfiram and myricetin did not affect metacestode vesicle integrity and showed no activity in the PGI assay (-0.2 ± 0.3 % and 0.1 ± 0.1 %, respectively). On the other hand, metacestode vesicles treated with sanguinarine all collapsed after five days of *in vitro* treatment, and strong activity was shown in the PGI assay (59.5 ± 2.2%, *p*=6e-6). We repeated this experiment once independently and obtained similar results with significant activity of sanguinarine against metacestode vesicles (S5 Fig). In the GL cell viability assay, DMSO resulted in the formation of round aggregates with a GL cell viability of 100 ± 7.4%. No aggregates were observed at all upon treatment with the positive control Triton X-100 and GL cell viability was highly reduced (-0.4 ± 0%, *p*=8e-7) (Fig 7B). Treatment of GL cell cultures with disulfiram resulted in irregularly shaped aggregates and strongly reduced GL cell viability (8 ± 3.5%, *p*=2e-6) while myricetin had no effect on aggregate formation nor on GL cell viability (107.8 ± 8.3%). No aggregates were formed upon treatment with sanguinarine, and GL cell viability was strongly reduced (0.5 ± 0.1%, *p*=8e-7). We repeated this experiment once independently and obtained similar results with significant activity of disulfiram and sanguinarine against GL cells (S5 Fig).

**Fig 7.**
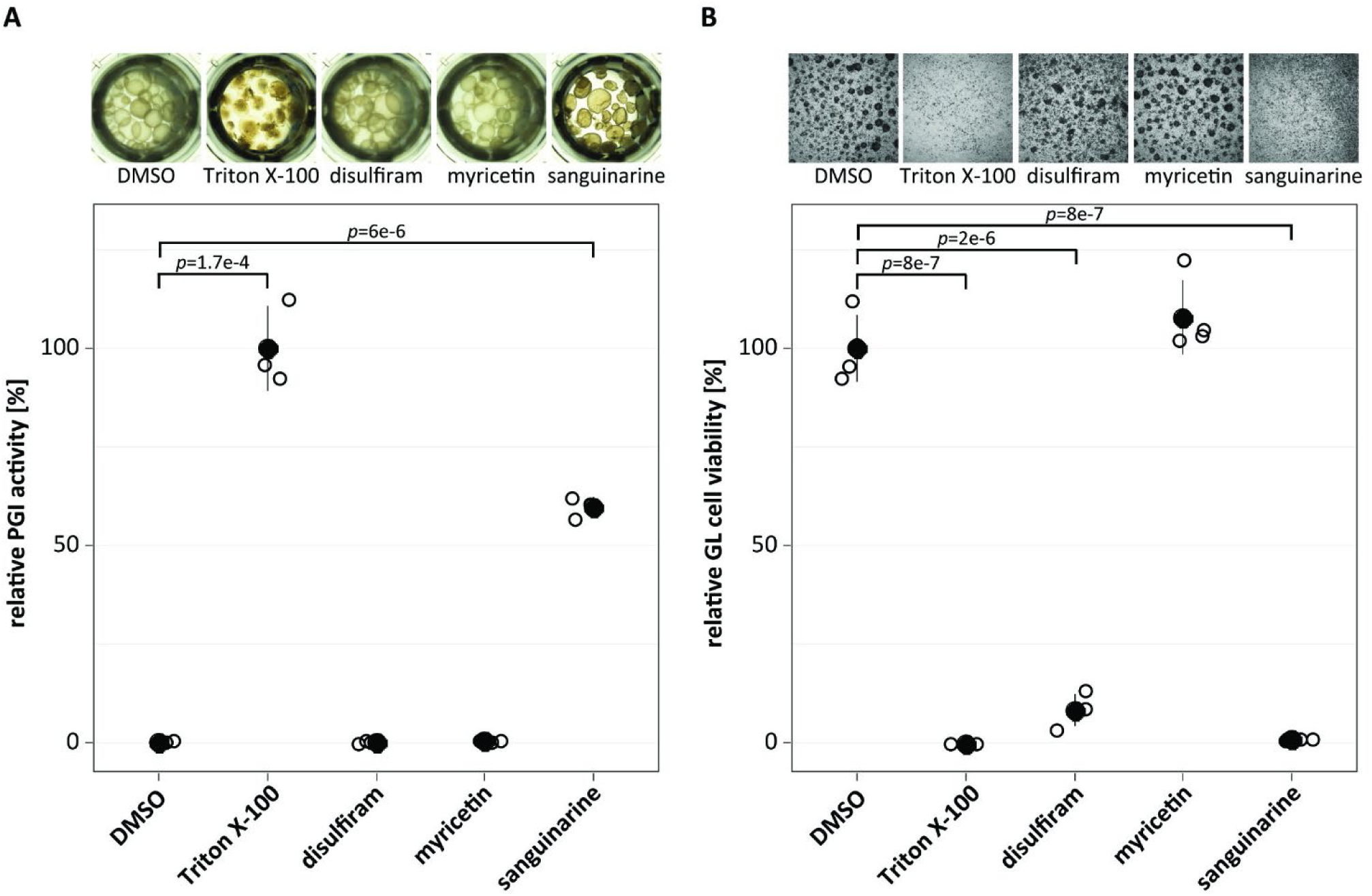
Effect of recEmTDH inhibitors on *E. multilocularis* metacestode vesicles and GL cell cultures. **A,** *E. multilocularis* metacestode vesicles were incubated with the negative control 0.1% DMSO, the positive control 0.1% Triton X-100, or the three recEmTDH inhibitors disulfiram, myricetin and sanguinarine at 20 µM (n=3 for each condition), the negative control 0.1% DMSO, or the positive control 0.1% Triton X-100. Metacestode vesicles were incubated under a humid, microaerobic atmosphere and after five days, pictures were taken and damage marker release was assessed relative to Triton X-100 treatment. Shown are photos of metacestode vesicles upon treatment, as well as PGI assay results with individual values (empty circles), mean values (filled circles), SDs and Bonferroni-corrected *p*-values of one representative experiment. The second independent experiment is shown in S5 Fig. B, *E. multilocularis* GL cell cultures were incubated with recEmTDH inhibitors disulfiram, myricetin and sanguinarine at 20 µM, the negative control 0.2% DMSO and the positive control 0.1% Triton X-100, (n=4 for each condition). GL cell cultures were incubated under a humid, microaerobic atmosphere. After five days, pictures were taken, and cell viability was measured and calculated relative to DMSO. Shown are individual values (empty circles) mean values (filled circles), SDs and Bonferroni-corrected *p*-values of one representative experiment. The other experiment is shown in S5 Fig.

Sanguinarine, as the most promising recEmTDH inhibitor with anti-echinococcal activity *in vitro*, was further characterized by calculating IC_50_-values for *E. multilocularis* metacestode vesicle damage, metacestode vesicle viability and GL cell viability. In addition, IC_50_-values for cytotoxicity on pre-confluent and confluent Hepa 1-6 cells were calculated (Table 2).

**Table 2:**
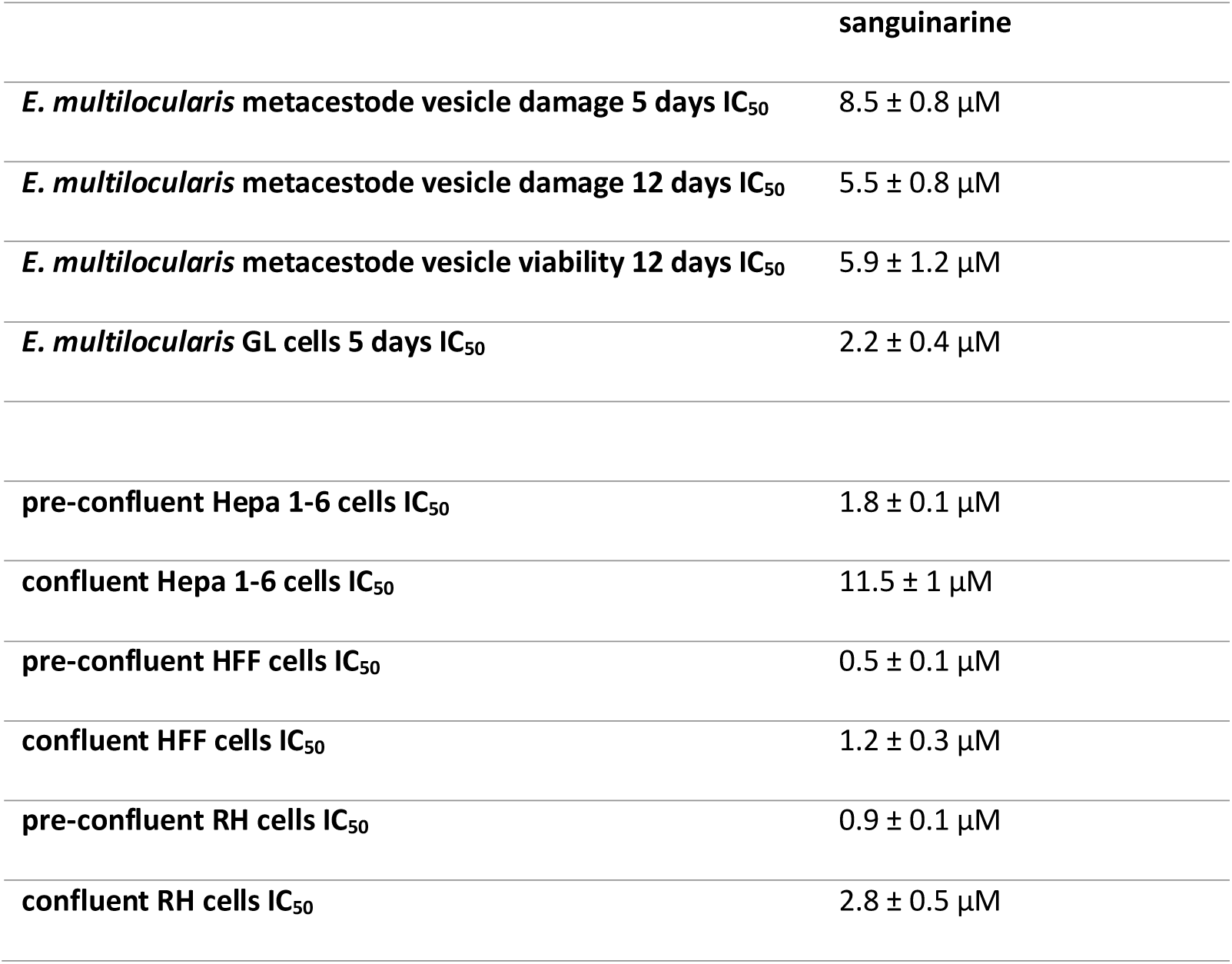
Half-maximal inhibitory concentrations of sanguinarine against *E. multilocularis* metacestode vesicles, GL cells and mammalian cells. Effect of sanguinarine was assessed against *E. multilocularis* on metacestode vesicles damage after five and twelve days via PGI assay and viability was assessed after twelve days via metacestode vesicle viability assay with concentrations of 40 to 1.25 µM in 1:2 serial dilutions (n=3) in relation to the respective DMSO controls. Effects of sanguinarine on *E. multilocularis* GL cells were assessed after five days via GL cell viability assay with concentrations of 40 to 0.08 µM in 1:2 serial dilutions (n=4) in relation to the respective DMSO controls. Mammalian cell toxicity of sanguinarine was assessed by cytotoxicity assays after five days with pre-confluent and confluent Hepa 1-6, HFF cells and RH cells with concentrations from 40 to 0.04 µM 1:2 serial dilutions (n=3). IC_50_-values were calculated for each of the three independent experiments per cell type and shown are mean values and SDs.

Sanguinarine resulted in activity against *E. multilocularis* metacestode vesicles with IC_50_-values of 8.5 ± 0.8 µM and 5.5 ± 0.8 µM after five and twelve days, respectively. Metacestode vesicle viability was in a similar range with an IC_50_-value of 5.9 ± 1.2 µM. Stronger effects were seen on GL cell cultures with an IC_50_-value of 2.2 ± 0.4 µM. However, sanguinarine also showed cytotoxicity against mammalian cells. We determined IC_50_-values of 1.8 ± 0.1 µM and 11.5 ± 1 µM for pre-confluent and confluent Hepa 1-6 cells, respectively. IC_50_-values for HFF cells were 0.5 ± 0.1 µM for pre-confluent cells and 1.2 ± 0.3 µM for confluent cells and IC_50_-values for RH cells were 0.9 ± 0.1 µM for pre-confluent cells, and 2.8 ± 0.5 µM for confluent cells, respectively.

## 4. Discussion

AE is a severe zoonotic disease with limited curative treatment options (8). New drugs are urgently needed and a better understanding of the metabolism may lead to the discovery of novel targets for interventions (20,37). Previously, our group has reported on the high uptake of the amino acid L-threonine by *E. multilocularis* metacestode vesicles *in vitro* (Ritler et al., 2019) and uptake of L-threonine was also reported from *E. granulosus s.l.* metacestodes *ex vivo* obtained from experimentally infected mice (78). Threonine as a source for energy generation has been suggested to play a role in various parasites such as *Entamoeba* spp. (79)*, T. brucei* (54,72) and *Trichomonas vaginalis* (80). In this project, we analyzed which effects L-threonine has on *E. multilocularis in vitro*, how L-threonine is metabolized and if the threonine-catabolic pathways would suggest potential targets for new interventions.

Growth assays provide simple and efficient tools to study the effects of nutrients on larval parasites. Different forms of measuring growth of *E. multilocularis* metacestode vesicles have been applied in the past, such as measuring their diameter via a microscope (81) or via volume measurements of single or multiple metacestode vesicles in falcon tubes (58,82,83). These assays provided valuable information regarding the improvement of *in vitro* culture conditions and the effects of growth factors on metacestode vesicles. However, when evaluating the effect of nutrients on single metacestode vesicles combined with an upscaling of replica, these measurement methods are highly time-consuming. Here, we established a growth assay using robust numbers of metacestode vesicles per condition (24 replica) in which the metacestode vesicle diameter is analyzed via automated and semi-automated scripts in ImageJ, enabling a fast and objective analysis. Both scripts reliably measured the diameter of metacestode vesicles faster and with a smaller error than the manual measurements. We thus employed these scripts to measure metacestode vesicle growth, which allowed us to use large numbers of replica. Our results showed that growth of *E. multilocularis* metacestode vesicles and development of metacestode vesicles from GL cell cultures was dependent on an active threonine metabolism, where L-threonine addition showed positive effects and its non-proteinogenic analogue 3-HNV negative effects. This could be explained by a possible competition of L-threonine and 3-HNV as TDH substrates, as suggested for TDH from mouse embryonic stem cells (60). This process also seemed to occur within *E. multilocularis* metacestode vesicles, as a combination of L-threonine and 3-HNV counteracted the negative effect of 3-HNV alone. We thus could show, that in a standardized *in vitro* setting (17), an active threonine metabolism is important for *E. multilocularis*. It is of note that we encountered variation in the number of metacestode vesicles formed from GL cell cultures, even in the control groups between different assays. These batch-effects were probably caused by the isolation and cultivation of varying numbers of GL cells.

To investigate which threonine catabolism pathways are active in *E. multilocularis in vitro*, we performed a [U-^13^C]-L-threonine flux assay that showed uptake of L-threonine by metacestode vesicles, metabolization to glycine and subsequent secretion of this metabolite to the culture medium, thus confirming previous reports (Ritler et al., 2019). The presence of [U-^13^C]-aminoacetone and [U-^13^C]-glycine, as well as lack of detection of α-ketobutyrate in our metabolomic samples, suggests an EmTDH-mediated threonine catabolism. In *C. elegans*, the acetyl-coenzyme A that is generated together with glycine via KBL feeds into the TCA cycle (43). Acetyl-coenzyme A was not detected in our samples, as the here applied detection method was not optimized for this metabolite. We detected a variety of TCA cycle intermediates, but none of them contained L-threonine-derived [^13^C]. This indicates that at least within the 24 hours of our *in vitro* setup under microaerobic conditions, L-threonine uptake did not feed into the TCA cycle. The protozoan parasite *T. brucei* also was reported to metabolize L-threonine via TDH and KBL and the generated acetyl-coenzyme A did not majorly feed into the TCA cycle, but rather was used for the synthesis of acetate and lipids (53,72,84–86). We did not detect acetate and although we detected propionate and palmitate in our samples, they did not incorporate [^13^C]. However, given that these metabolites were also present in the blank samples, we cannot exclude that a small proportion contained [^13^C] that was masked by the blank samples.

Very interestingly, we detected [^13^C_2_]-glutathione in VT, suggesting that *E. multilocularis* metacestode vesicles synthesize glutathione *de novo* using threonine-derived glycine. Besides its role for DNA synthesis (87), glutathione can also act protective against oxidative stress by neutralizing free radicals (88). Further studies are needed to investigate whether the same applies for *E. multilocularis*.

Threonine has been shown to be essential for cell proliferation and DNA synthesis of mouse embryonic stem cells (60). Threonine is hereby metabolized via TDH to glycine and acetyl-coenzyme A, which is used for the synthesis of *S*-adenosylmethionine (SAM) (89). Restriction of threonine from *in vitro* cultured mouse embryonic stem cells decreased trimethylation of histone H3 lysine 4 (89). SAM was not detected in our set up, but its precursor L-methionine did not incorporate [^13^C]. However, potential effects of L-threonine on the histone modification in *E. multilocularis* metacestode vesicles were not further focus of our study.

Besides the results of the [U-^13^C]-L-threonine flux assay, we also measured significantly higher gene expression of *emtdh* and *emkbl* compared to *emtd* in metacestode vesicles, confirming previously reported results in metacestode vesicles *in vitro* and metacestodes *in vivo* (39,90). Furthermore, EmTDH and EmKBL, but not EmTD, were detected via non-targeted proteomics in VF and VT of *in vitro* cultured metacestode vesicles and also in VF of *in vivo* grown metacestodes obtained from experimentally infected mice (17). Interestingly, EmTD was not detected in these samples. Finally, by testing crude extracts of *in vitro* cultured *E. multilocularis* metacestode vesicles in an enzymatic assay, we confirmed that EmTDH was translated into an enzymatically active protein. Taken together, our experiments strongly suggest that EmTDH is the major threonine catabolic enzyme in *in vitro* cultured metacestode vesicles. Given that L-threonine did not feed into the TCA cycle in our [U-^13^C]-L-threonine flux assay, energy generation via the TCA cycle cannot explain the positive effects of L-threonine on metacestode vesicle growth and development. Since L-threonine feeds into the synthesis of glutathione, positive effects might have been caused by reduced oxidative stress due to higher amounts of reduced glutathione within the parasite. Future studies need to investigate this.

Since our experiments confirmed the importance of a TDH-mediated threonine metabolism in *E. multilocularis in vitro*, we wanted to investigate whether this enzyme could serve as a potential drug target candidate. We recombinantly expressed MmTDH alongside since human TDH is a non-functional pseudogene (56) and potential selective recEmTDH inhibitors would be first tested in the mouse models of AE (91,92). MmTDH is highly expressed in embryonic stem cells but shows low expression in tissue of different organs from adult mice (60).

We tested a series of QCs, which have been found active against recMmTDH upon a high-throughput screen of 20,000 compounds (74) and additionally disulfiram, myricetin, quercetin and sanguinarine, which have been reported to show activity against recTbTDH from *T. brucei* (73). Disulfiram, myricetin and sanguinarine strongly inhibited EmTDH activity, but when tested on *E. multilocularis* metacestode vesicles, only sanguinarine displayed anti-parasitic effects. Disulfiram exhibited, in addition to sanguinarine, activity against GL cells of *E. multilocularis,* but was not further followed up due to its inefficacy against metacestode vesicles in the PGI assay confirming previous reports (77). Sanguinarine was previously reported to perturb anterior regeneration of the planarian *Dugesia japonica* (93). Furthermore, sanguinarine was shown be active against the ciliate *Ichthyophthirius multifiliis in vitro* and in an *in vivo* grass carp (*Ctenopharyngodon idella*) model (94), as well as against the nematodes *Toxocara canis in vitro* (95) and *Trichinella spiralis in vitro* and in an *in vivo* mouse model (96). Regarding platyhelminths, effects of sanguinarine were reported against the monogenean parasite *Dactylogyrus intermedius* in *in vivo* models with goldfish (*Carassius auratus*) (97,98), the trematode *Schistosoma mansoni in vitro* (23,99) and *in vitro* cultured protoscoleces of the cestode *E. granulosus sensu lato* (100). In the present study, we calculated IC_50_-values against *E. multilocularis* metacestode vesicles and GL cells *in vitro* and the respective concentrations of sanguinarine were substantially lower than reported for *T. spiralis* (96). Since in an *in vivo* mouse model sanguinarine caused significantly reduced worm burdens of *T. spiralis* (96), there might also be a potential therapeutic window for mice infected with *E. multilocularis*. Future studies should investigate if this compound also shows activity in AE mouse models.

It is crucial in the future to evaluate the relevance of EmTDH-mediated threonine metabolism for *E. multilocularis in vivo* and further confirm EmTDH as a potential drug target. Upon its validation, the here established enzymatic assay for EmTDH could serve as a discovery platform that allows for targeted medium-throughput screening of inhibitors. It should be further investigated whether this tool could also be used to screen potential inhibitors against TDH of the closely related *E. granulosus s.s.* (EgTDH: EgrG_000511900, PRJEB121) and *E. canadensis* (EcTDH: EcG7_08078, PRJEB8992), which are the main causative agents for human cystic echinococcosis. Active drugs could then be confirmed via the here mentioned whole-organism-based *in vitro* screening assays, as they have been recently validated for *E. granulosus s.s.* (30).

## Acknowledgements

The authors thank Arunasalam Naguleswaran (Institute of Animal Pathology, University of Bern) and Joachim Müller (Institute of Parasitology, University of Bern) for the helpful discussion regarding the recombinant expression of recEmTDH and recMmTDH and the analysis of TDH activity. The authors also thank Magali Roques (Institute of Cell Biology, University of Bern) for kindly sharing the Hepa 1-6 cells. We thank the Analytical Services from the Department of Chemistry, Biochemistry and Pharmaceutical Sciences, University of Bern, Switzerland, for measuring NMR and MS spectra of synthetic intermediates and final QC compounds. We thank Andrew Hemphill (Institute of Parasitology, University of Bern) for critical reading of the manuscript.

## Author contribution

MK and BLS designed the study. MK, PZ, MP and AB performed most of the experiments and the long-term culture of the parasites. PG and ML synthesized the quinazoline carboxamides. SS performed the HPLC experiment to determine the concentration of L-threonine in the culture medium. CR measured and analyzed the metabolic samples of the [U-^13^C]-L-threonine flux assay. MK and DVR developed and validated the automated and semi-automated scripts for the metacestode vesicle growth assay. MK, PZ and BLS performed the data analysis. MK and BLS drafted the original version of the manuscript, finalized it and prepared the figures. All authors read and approved the manuscript.

## Supporting information

**S1 File: Synthesis of Quinazoline carboxamides. S2 File. Automated script in ImageJ.**

**S3 File. Semi-automated script in ImageJ.**

**S4 File. Analysis of amino acids in supernatant samples.**

**S5 File. Cloning and recombinant expression of EmTDH and MmTDH.**

**S1 Table. Blast of protein sequences of TD, TDH, KBL and TA of reference organisms against *E. multilocularis* protein database PRJEB122.** Shown are reference organisms (*C. elegans*, *Da. rerio*, *Dr. melanogaster*, *H. sapiens* and *M. musculus*) with accession numbers of TA, TD, TDH and KBL. Shown are sequences that produced a significant alignment with the corresponding E-values.

**S2 Table. Primer sequences, gene description and accession numbers for *emtd*, *emtdh*, *emkbl* and the house-keeping gene *emelp*.**

**S3 Table. Labeling patterns of potential metabolites of L-threonine.** Abbreviations: VT = vesicle tissue, VF = vesicle fluid, CM = culture medium, VM = vesicle medium.

**S4 Table. Reciprocal blast of EmuJ_001093200 against coding sequences of reference organisms.** Shown are significant hits against coding sequences of reference organisms (*C. elegans*, *Da. rerio*, *Dr. melanogaster*, *H. sapiens* and *M. musculus*) with protein description, organism, max score, total score, query cover, E-values, identity and the accession numbers.

**S5 Table. Reciprocal blast of EmuJ_000511900 against coding sequences of reference organisms.** Shown are significant hits against coding sequences of reference organisms (*C. elegans*, *Da. rerio*, *Dr. melanogaster*, *H. sapiens* and *M. musculus*) with protein description, organism, max score, total score, query cover, E-values, identity and the accession numbers.

**S6 Table. Reciprocal blast of EmuJ_000107200 against coding sequences of reference organisms.** Shown are significant hits against coding sequences of reference organisms (*C. elegans*, *Da. rerio*, *Dr. melanogaster*, *H. sapiens* and *M. musculus*) with protein description, organism, max score, total score, query cover, E-values, identity and the accession numbers.

**S7 Table. Blast of TDH protein sequences of reference organisms against *E. granulosus s.s.* protein database PRJEB121.** Shown are the accession numbers of TDH sequences of reference organisms (*C. elegans*, *Da. rerio*, *Dr. melanogaster*, *H. sapiens* and *M. musculus*), as well as sequences that produced significant alignments with the corresponding E-values.

**S8 Table. Blast of TDH protein sequences of reference organisms against *E. canadensis* protein database PRJEB8992.** Shown are the accession numbers of TDH sequences of reference organisms (*C. elegans*, *Da. rerio*, *Dr. melanogaster*, *H. sapiens* and *M. musculus*), as well as sequences that produced a significant alignment with the corresponding E-values.

**S9 Table. Reciprocal blast of EgrG_000511900 against coding sequences of reference organisms.** Shown are significant hits against coding sequences of reference organisms (*C. elegans*, *Da. rerio*, *Dr. melanogaster*, *H. sapiens* and *M. musculus*) with protein description, organism, max score, total score, query cover, E-values, identity and the accession numbers.

**S10 Table. Reciprocal blast of EcG7_08078 against coding sequences of reference organisms.** Shown are significant hits against coding sequences of reference organisms (*C. elegans*, *Da. rerio*, *Dr. melanogaster*, *H. sapiens* and *M. musculus*) with protein description, organism, max score, total score, query cover, E-values, identity and the accession numbers.

**S11 Table. Bonferroni-corrected *p*-values for the relative enzymatic activity of recMmTDH treated with different compounds at 20 μM.**

**S1 Fig. Comparison of measurement methods for *E. multilocularis* metacestode vesicles.** 50 *E. multilocularis* metacestode vesicles were photographed three times each. The diameter of each metacestode vesicle was measured via an automated script with a mean of 360 diameters (black circles), via a semi-automated script with a mean of 360 diameters (dark gray squares), or manually with a mean of two diameters (light gray triangles). Shown are mean values and SD for the three images per metacestode vesicle.

**S2 Fig. Reduction of L-threonine in culture medium with *E. multilocularis* metacestode vesicles.**

*E. multilocularis* metacestode vesicles were cultured in conditioned medium to which L-threonine was added in various concentrations (0 mM, 2 mM, 4 mM, 8 mM or 12 mM) under microaerobic conditions in triplicates. The reduction of L-threonine after four days of incubation was measured in supernatant samples via HPLC. A, metacestode vesicle diameter measured via the semi-automated script. B, reduction of L-threonine in the culture medium. Shown are mean values, SD and Bonferroni-corrected *p-*values.

**S3 Fig. Effect of threonine on *E. multilocularis* metacestode vesicle growth and development *in vitro*. A,** *E. multilocularis* metacestode vesicles were cultured with various amounts of L-threonine (1mM, 2 mM, 4 mM), D-threonine (4 mM), or water as a control (n=24 for each condition). B, *E. multilocularis* metacestode vesicles were cultured with 3-HNV (4 mM, n=23), a combination of 3-HNV and L-threonine (each at 4 mM, n=24), or water (n=24). Metacestode vesicle growth in A and B was calculated via an automated script or a semi-automated script and shown is growth in mm and Bonferroni-corrected *p*-values as compared to the water controls. C, E. multilocularis GL cells were cultured with L-threonine (4 mM), D-threonine (4 mM), or water as a control (n=4 for each condition). D, *E. multilocularis* GL cells were cultured with 3-HNV (4 mM), a combination of 3-HNV and L-threonine (both at 4 mM), or the water control (n=4 for each condition). Newly formed metacestode vesicles in C and D were counted manually in a blinded manner and shown are mean values, SD and Bonferroni-corrected *p*-values. Abbreviations: Thr = threonine, 3-HNV = 3-hydroxynorvaline.

**S4 Fig. Relative expression of threonine metabolism genes in *E. multilocularis* metacestode vesicles under microaerobic conditions.** Relative expression was analyzed in *E. multilocularis* metacestode vesicles cultured *in vitro* (n=4). Q-RT-PCRs were performed in technical duplicates for each sample and gene expression was calculated relative to the housekeeping gene ELP. Shown are Bonferroni-corrected *p-*values.

**S5 Fig. Effect of recEmTDH inhibitors against *E. multilocularis* metacestode vesicles and GL cell cultures. A,** *E. multilocularis* metacestode vesicles were incubated with the three recEmTDH inhibitors disulfiram, myricetin and sanguinarine at 20 μM (n=3 for each condition), the negative control 0.1% DMSO, or the positive control 0.1% Triton X-100. Metacestode vesicles were incubated under a humid, microaerobic atmosphere and after five days, pictures were taken and damage marker release was assessed relative to Triton X-100 treatment. Shown are photos of metacestode vesicles upon treatment, as well as PGI assay results with individual values (empty circles), mean values (filled circles), SD and Bonferroni-corrected *p*-values of one representative experiment. B, *E. multilocularis* GL cell cultures were incubated with recEmTDH inhibitors disulfiram, myricetin and sanguinarine at 20 μM, the negative control 0.2% DMSO and the positive control 0.1% Triton X-100, (n=4 for each condition). GL cell cultures were incubated under a humid, microaerobic atmosphere. After five days, pictures were taken, and GL cell viability was measured and calculated relative to DMSO. Shown are individual values (empty circles) mean values (filled circles), SD and Bonferroni-corrected *p-*values.

## Notes

### Competing Interest Statement

The authors have declared no competing interest.

