## Supplementaries for "Investigation of the threonine metabolism of *Echinococcus multilocularis*: the threonine dehydrogenase as a potential drug target in alveolar echinococcosis": S1 File.docx

**S1 File: Synthesis of Quinazoline carboxamides.**

### General remarks

All reactions requiring anhydrous conditions were performed in heat-gun, oven or flame dried glassware under inert atmosphere (Ar). Silica gel 60 Å (40–63 mm) from Sigma-Aldrich was used for dry loads. Flash column chromatography was performed on a Teledyne Isco CombiFlash® Rf+ with the corresponding RediSep® prepacked silica cartouches unless otherwise stated. Thin layer chromatography (TLC) was performed on Macherey & Nagel Alugram® xtra SIL G/UV 254 visualization under UV light (254 nm) and/ or (366 nm) and/ or by dipping in anisaldehyde stain and subsequent heating.

Commercial reagents and solvents (Acrôs Organics, Apollo scientific, Combi-Blocks, Fluorochem, Grogg Chemie, Hänseler, Sigma-Aldrich) were used without further purification unless otherwise stated. Dry solvents for reactions were distilled and filtered over columns of dry neutral aluminium oxide under positive argon pressure. Solvents for extraction and flash chromatography were used without further purification.

^1^H , ^13^C and ^19^F NMR spectra were recorded on a Bruker AVANCE-300 or 400 spectrometers operating at 300 or 400 MHz for ^1^H, 75 or 101 MHz for ^13^C and 376 MHz for ^19^F, at room temperature unless otherwise stated. Chemical shifts (*δ*) are reported in parts per million (ppm) relative to tetramethylsilane (TMS), calibrated using residual signals of the solvent or TMS. Coupling constants (*J*) are reported in Hz. HRMS analyses and accurate mass determinations were performed on a Thermo Scientific LTQ Orbitrap XL mass spectrometer using ESI ionisation and positive or negative mode by the analytical services (mass spectrometry lab of Prof. Dr. Stefan Schürch) from the Department of Chemistry, Biochemistry and Pharmaceutical Sciences (DCBP) of the University of Bern, Switzerland.

### Synthesis of quinazoline carboxamides QC1-6

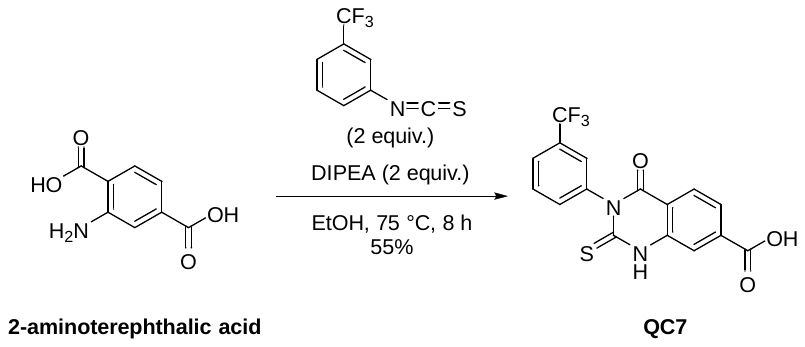

**4-oxo-2-thioxo-3-(3-(trifluoromethyl)phenyl)-1,2,3,4-tetrahydroquinazoline-7-carboxylic acid (QC7).** To a stirred suspension of 2-aminoterephthalic acid (0.5079 g, 2.8038 mmol) in absolute ethanol (6 mL) at 75 °C DIPEA (1 mL, 0.76 g, 5.88 mmol) was added, followed by 3-(trifluoromethyl)phenyl thioisocyanate (0.8 mL, 1.068 g, 5.25 mmol). The mixture was placed under an argon atmosphere and stirred for 8 h. The mixture was concentrated under reduced pressure. The residue was taken up in acetone and concentrated onto silica gel. The crude was purified by flash column chromatography (cyclohexane/ (ethyl acetate/ acetic acid 9:1) gradient from 1:0 to 0:1). The fractions containing the desired product were concentrated and co-evaporated with toluene to remove the acetic acid, which gave the desired product in 55% yield (0.56025 g, 1.52942 mmol). The product was still contaminated with ethyl acetate and acetic acid (purity ~83% by NMR) and used in the next step without further purification. An aliquot was dried to constant weight for biological testing. White solid: **^1^H NMR** (300 MHz, DMSO-*d*_6_) δ 13.28 (s, 1H), 12.84 (s, 1H), 8.06 (d, *J* = 8.2 Hz, 1H), 8.02 (d, *J* = 1.4 Hz, 1H), 7.87 – 7.77 (m, 3H), 7.77 – 7.70 (m, 1H), 7.66 (dt, *J* = 7.7, 1.8 Hz, 1H). **^13^C NMR** (75 MHz, DMSO) δ 176.24, 166.08, 159.51, 139.98, 139.62, 136.94, 133.55, 130.18, 130.02, 129.60, 127.99, 126.33, 126.27, 125.77, 125.13, 124.27, 122.16, 119.25, 116.86. **^19^F NMR** (282 MHz, DMSO) δ -60.97. **HRMS** (ESI) calculated for [M+H]^+^ C_16_H_10_F_3_N_2_O_3_S 367.0359, found 367.0342.

**^1^H NMR**

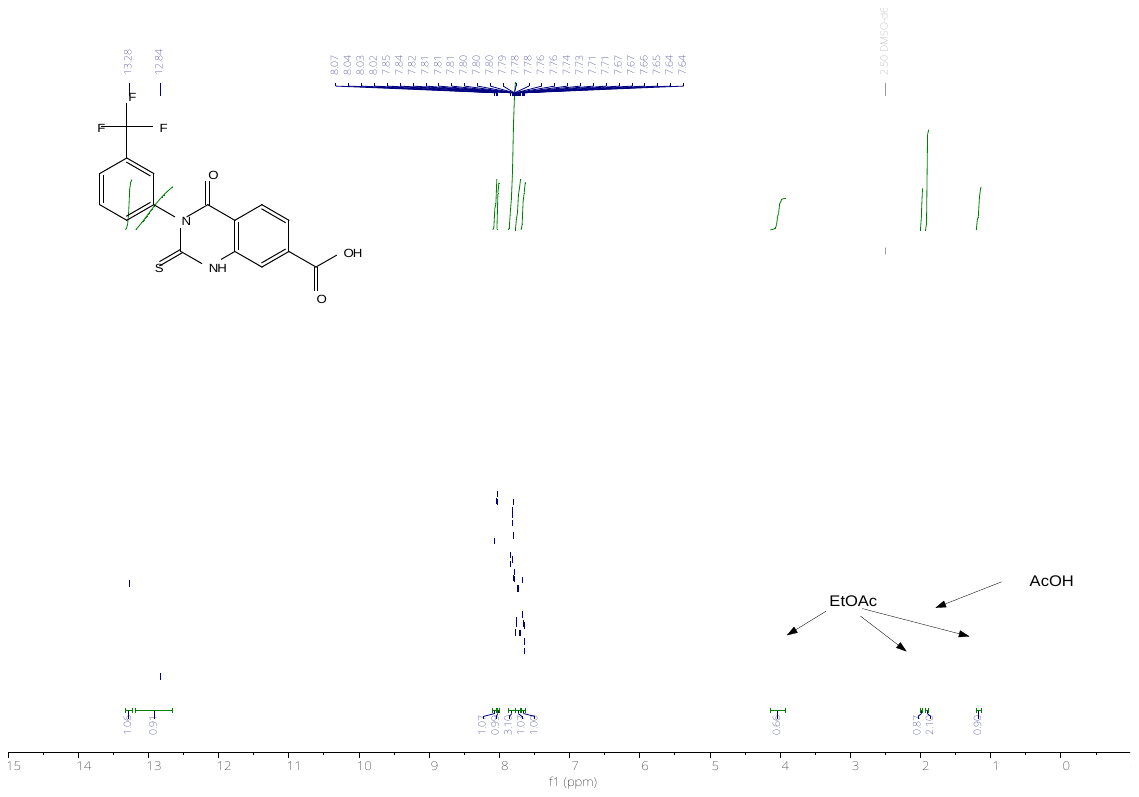

**^13^C NMR**

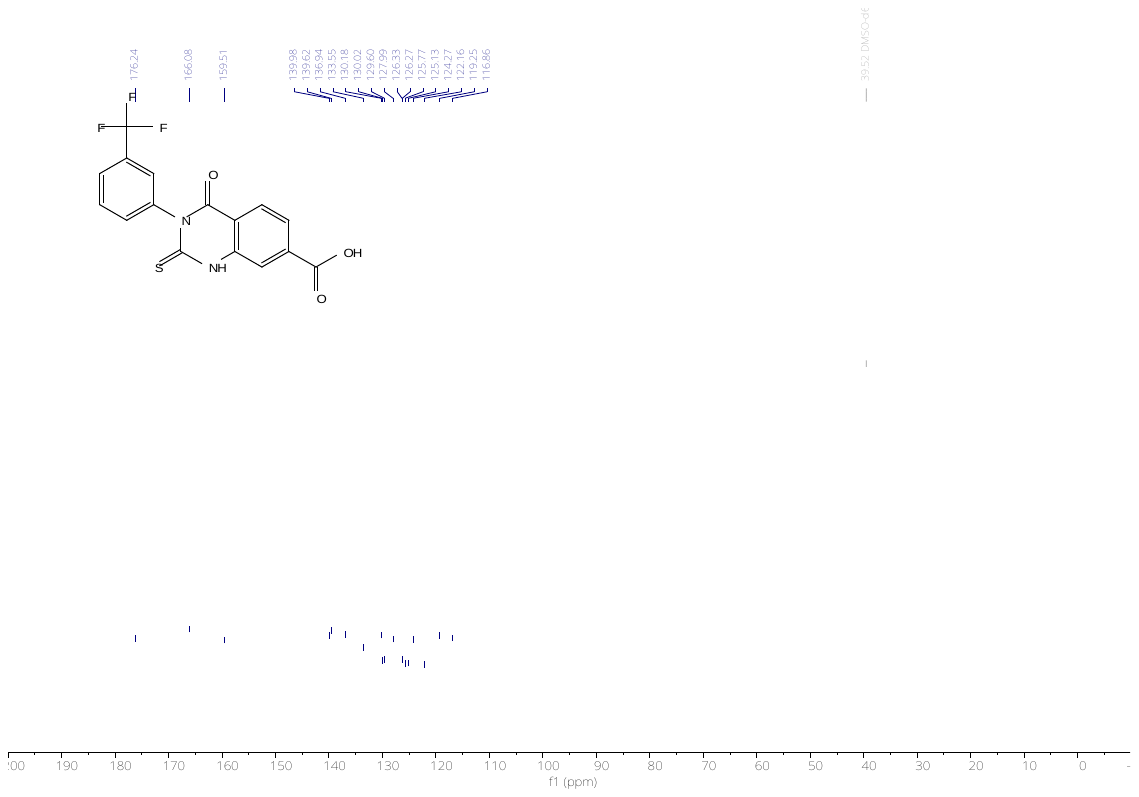

**^19^F NMR**

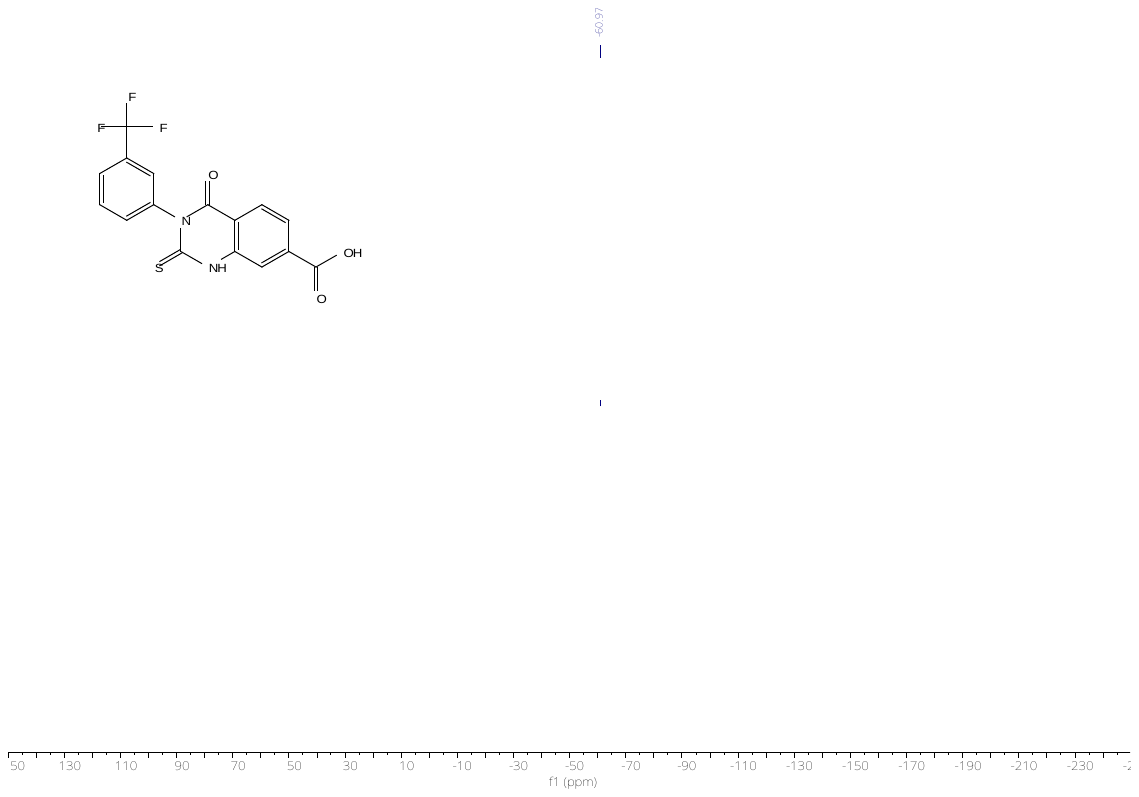

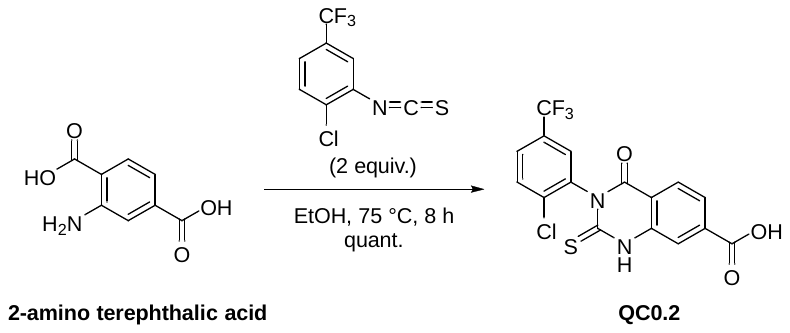

**3-(2-chloro-5-(trifluoromethyl)phenyl)-4-oxo-2-thioxo-1,2,3,4-tetrahydroquinazoline-7-carboxylic acid (QC0.2).** To a stirred suspension of 2-aminoterephthalic acid (0.11331 g, 0.6255 mmol) in absolute ethanol (2 mL), DIPEA (0.180 µL, 0.1368 g, 1.058 mmol) and 1-chloro-2-isothiocyanato-4-(trifluoromethyl)benzene (0.180 µL, 0.261 g, 1.098 mmol) was added. The mixture was heated to 75 °C and stirred for 8 h. The mixture was concentrated under reduced pressure. The residue was taken up in acetone and concentrated onto silica gel. The crude was purified by flash column chromatography (cyclohexane/ (ethyl acetate/ acetic acid 9:1) gradient from 1:0 to 0:1). The fractions containing the desired product were concentrated and co-evaporated with toluene to remove residual acetic acid, to give the desired product in quantitative yield (0.32843 g, 0.81953 mmol). The product was contaminated with toluene (purity ~80% by NMR) and used in the next step without further purification. Yellow solid: **^1^H NMR** (300 MHz, DMSO-*d*_6_) δ 13.49 (s, 2H), 8.16 – 7.99 (m, 3H), 7.94 – 7.83 (m, 3H). **^13^C NMR** (75 MHz, DMSO) δ 175.16, 165.97, 158.71, 139.71, 137.63, 137.33, 136.40, 130.99, 129.04, 128.61, 128.10, 127.16, 124.63, 121.67, 118.43, 117.03. **^19^F NMR** (282 MHz, DMSO) δ -61.04. **HRMS** (ESI) calculated for [M+H]^+^ C_16_H_9_ClF_3_N_2_O_3_S 400.9969, found 400.9957.

**^1^H NMR**

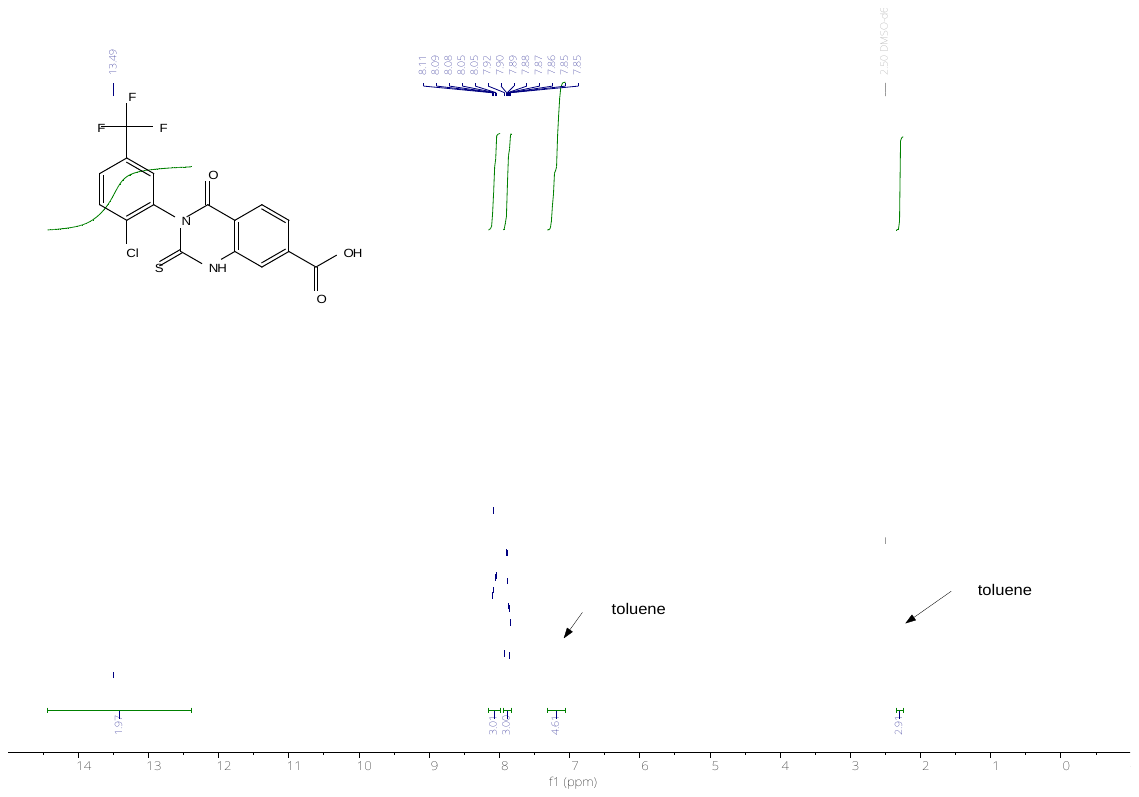

**^13^C NMR**

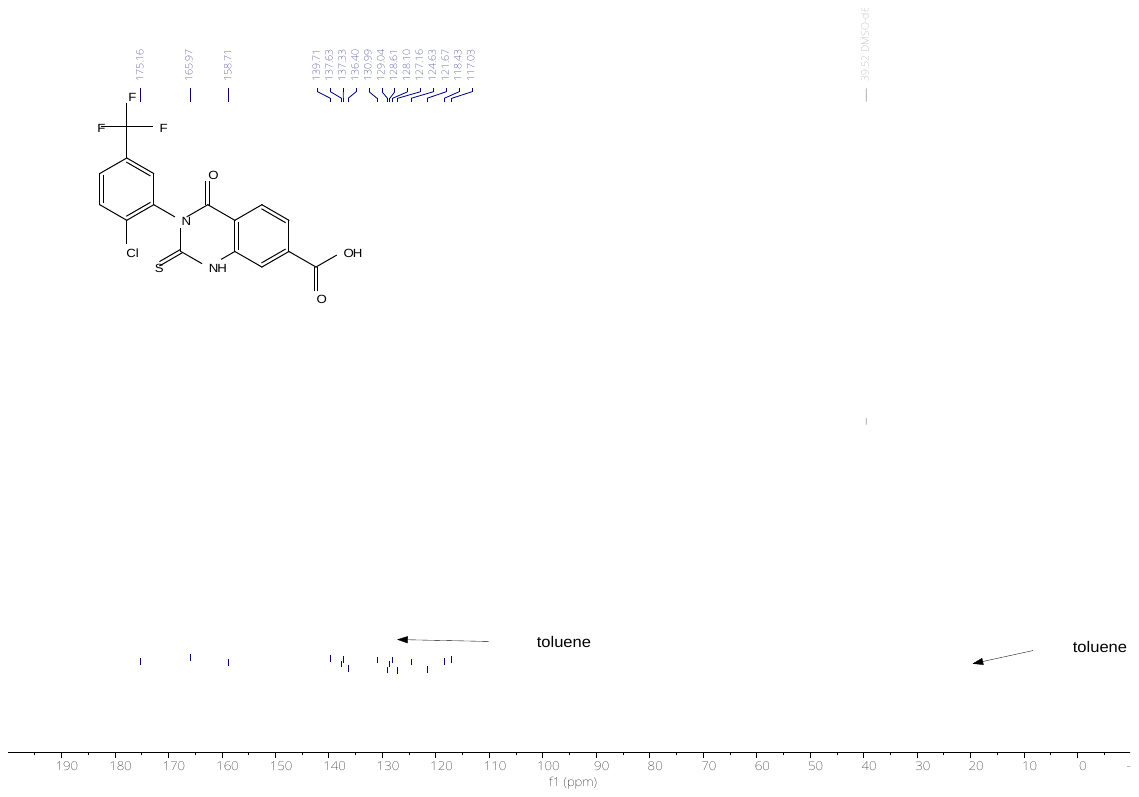

**^19^F NMR**

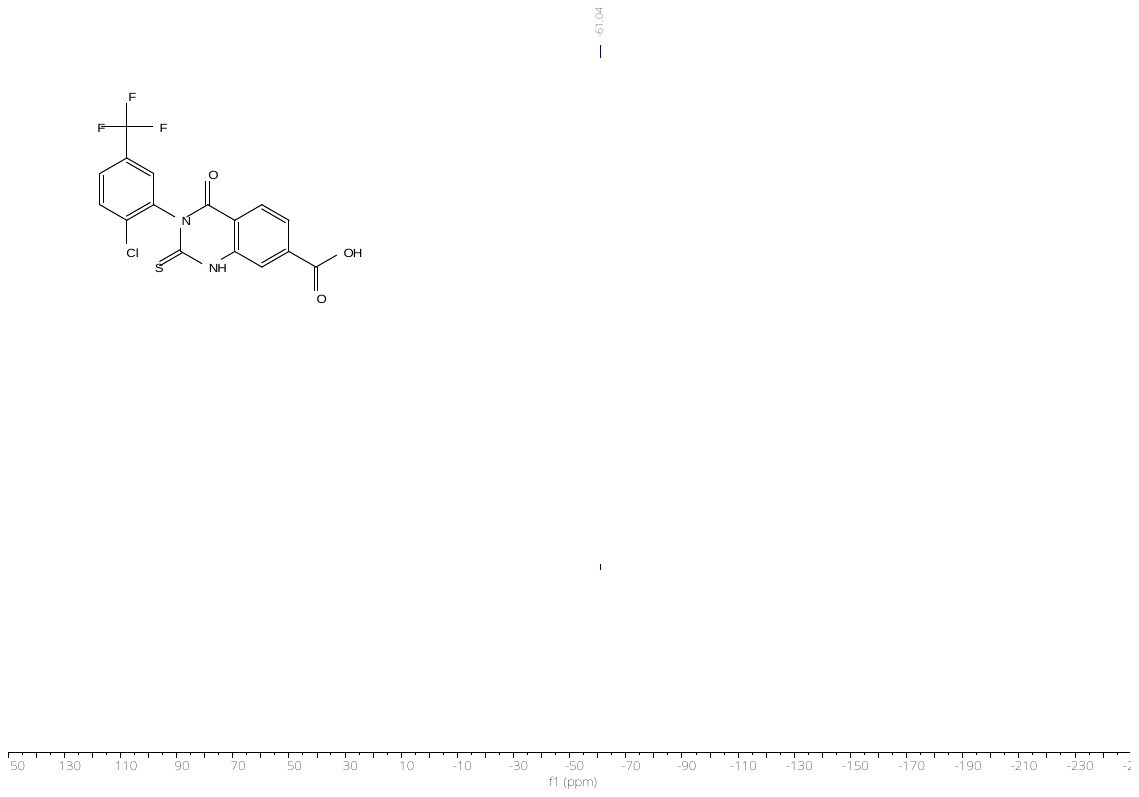

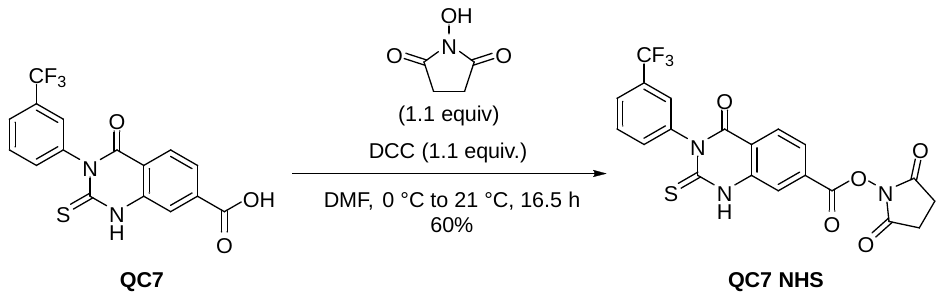

**2,5-dioxopyrrolidin-1-yl 4-oxo-2-thioxo-3-(3-(trifluoromethyl)phenyl)-1,2,3,4-tetrahydroquinazol-ine-7-carboxylate (QC7 NHS).** To a stirred solution of 4-oxo-2-thioxo-3-(3-(trifluoromethyl)phenyl)-1,2,3,4-tetrahydroquinazoline-7-carboxylic acid (0.20167 g, 0.5504 mmol) in dry DMF at 0 °C under an argon atmosphere was added DCC (0.12390 mg, 0.6006 mmol). The mixture was stirred for 30 minutes. Then *N*-hydroxysuccinimide (00931 g, 0.80893 mmol) was added. The mixture was stirred for 16 h. The volatiles were removed under reduced pressure. The residue was taken up in acetone and concentrated onto silica gel. The crude was purified by flash column chromatography (cyclohexane/ ethyl acetate gradient from 1:0 to 0:1) to give the desired product in 94% (0.23975 g, 0.51738 mmol). Product was contaminated with cyclohexane and ethyl acetate (purity ~64% by NMR) and used in the next step without further purification. White solid: **^1^H NMR** (300 MHz, DMSO-*d*_6_) δ 13.36 (s, 1H), 8.19 (d, *J* = 8.3 Hz, 1H), 8.16 (d, *J* = 1.6 Hz, 1H), 7.96 (dd, *J* = 8.3, 1.6 Hz, 1H), 7.89 – 7.59 (m, 4H), 2.93 (s, 4H). **^13^C NMR** (75 MHz, DMSO) δ 176.48, 170.14, 160.95, 159.22, 139.83, 133.47, 130.26, 130.06, 129.63, 129.01, 126.21 (q, *J* = 3.9 Hz), 125.74, 125.25 (m), 124.32, 122.13, 121.15, 117.45, 25.62. **^19^F NMR** (282 MHz, DMSO) δ -60.98. **HRMS** (ESI) calculated for [M+H]^+^ C_20_H_13_F_3_N_3_O_5_S 464.0523, found 464.0511.

**^1^H NMR**

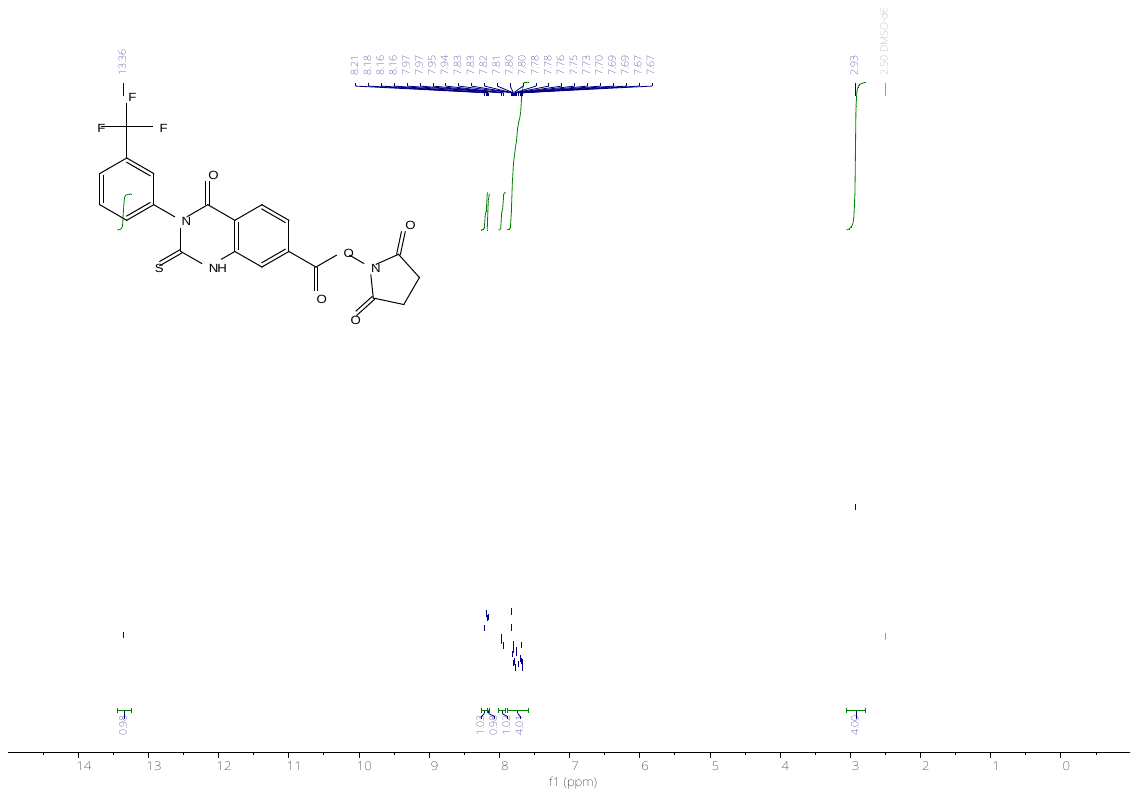

**^13^C NMR**

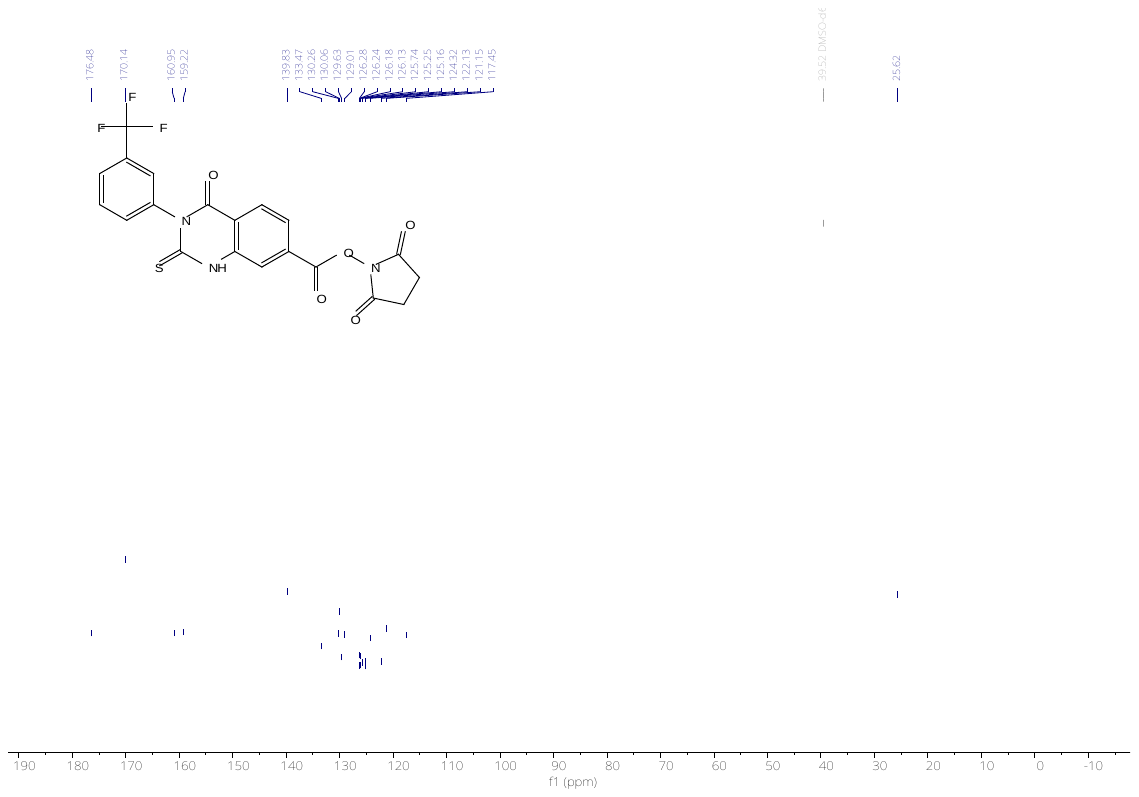

**^19^F NMR**

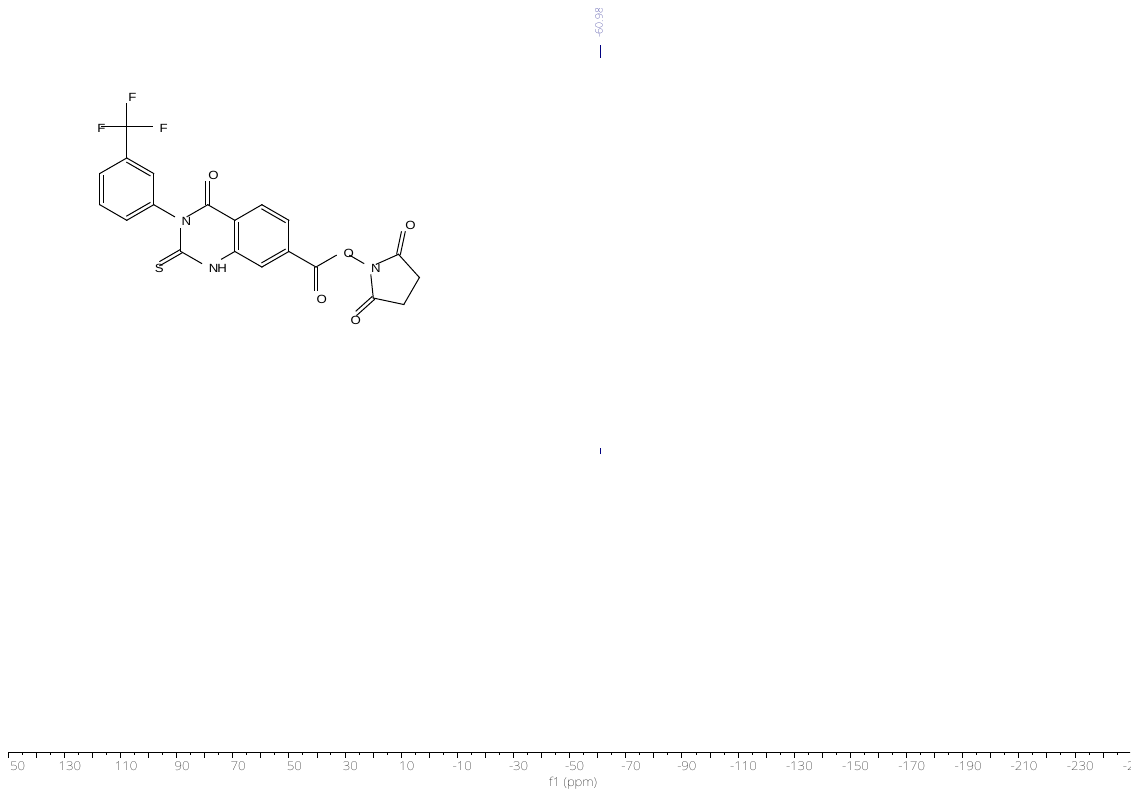

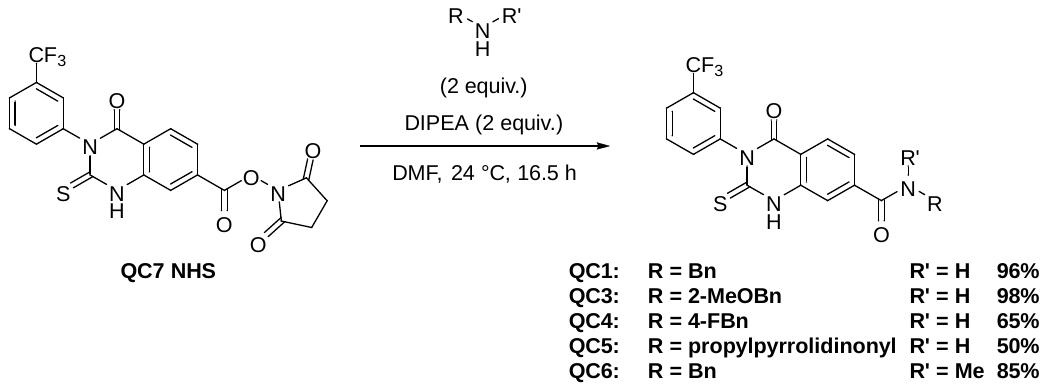

**General procedure I: Synthesis of QC amides from QC7 NHS**

To a stirred solution of 2,5-dioxopyrrolidin-1-yl 4-oxo-2-thioxo-3-(3-(trifluoromethyl)phenyl)-1,2,3,4-tetrahydroquinazoline-7-carboxylate (**QC7 NHS**) in dry dimethylformamide (0.1 M) at 24 °C under an argon atmosphere *N,N*-Diisopropylethylamine (2 equiv.) was added. Then, the corresponding amine (2 equiv.) was added. The mixture was stirred for 24 h. The volatiles were removed under reduced pressure. The residue was taken up in a mixture of acetonitrile and methanol and concentrated onto silica gel. The crude was purified by reverse phase flash column chromatography (water + 0.1% TFA/ acetonitrile + 0.1% TFA gradient from 1:0 to 0:1), to give the desired product.

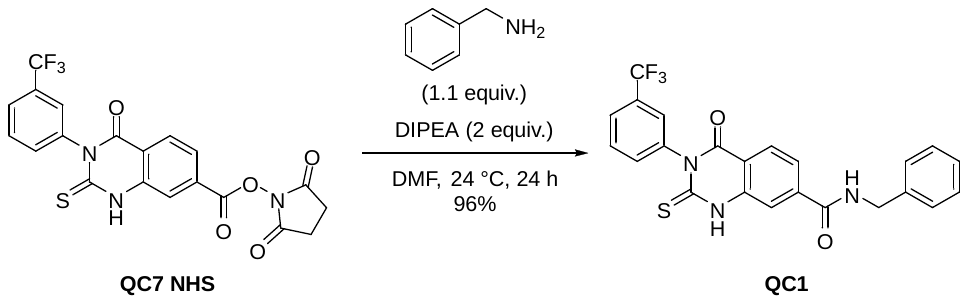

***N*-benzyl-4-oxo-2-thioxo-3-(3-(trifluoromethyl)phenyl)-1,2,3,4-tetrahydroquinazoline-7-carboxamide (QC1).** Prepared according to general procedure I: 96% yield (0.09529 g, 0.2094 mmol). White solid: **^1^H NMR** (400 MHz, DMSO-*d*_6_) δ 13.26 (s, 1H), 9.35 (t, *J* = 6.0 Hz, 1H), 8.04 (d, *J* = 8.2 Hz, 1H), 7.91 (d, *J* = 1.5 Hz, 1H), 7.83 – 7.76 (m, 3H), 7.73 (t, *J* = 7.7 Hz, 1H), 7.66 (d, *J* = 8.1 Hz, 1H), 7.36 (d, *J* = 4.3 Hz, 4H), 7.30 – 7.24 (m, 1H), 4.51 (d, *J* = 5.9 Hz, 2H). **^13^C NMR** (101 MHz, DMSO) δ 176.18, 165.07, 159.51, 140.92, 140.01, 139.59, 139.21, 133.56, 130.12, 129.74 (q, *J* = 32.1 Hz), 128.35, 127.72, 127.34, 126.90, 126.43 – 126.17 (m), 125.30, 125.06 (d, *J* = 3.4 Hz), 122.59, 122.26, 115.38, 42.86, 30.67. **^19^F NMR** (282 MHz, DMSO) δ -60.93. **HRMS** (ESI) calculated for [M+H]^+^ C_23_H_17_F_3_N_3_O_2_S 456.0988, found 456.0979.

**^1^H NMR**

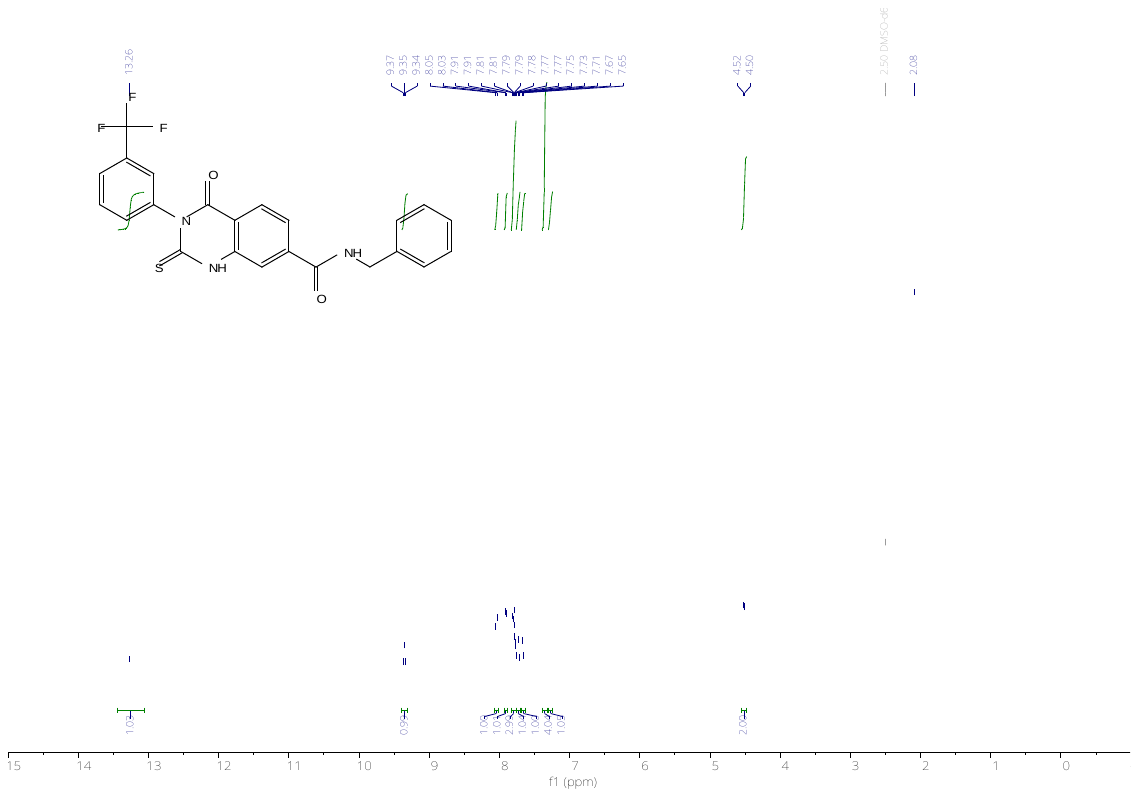

**^13^C NMR**

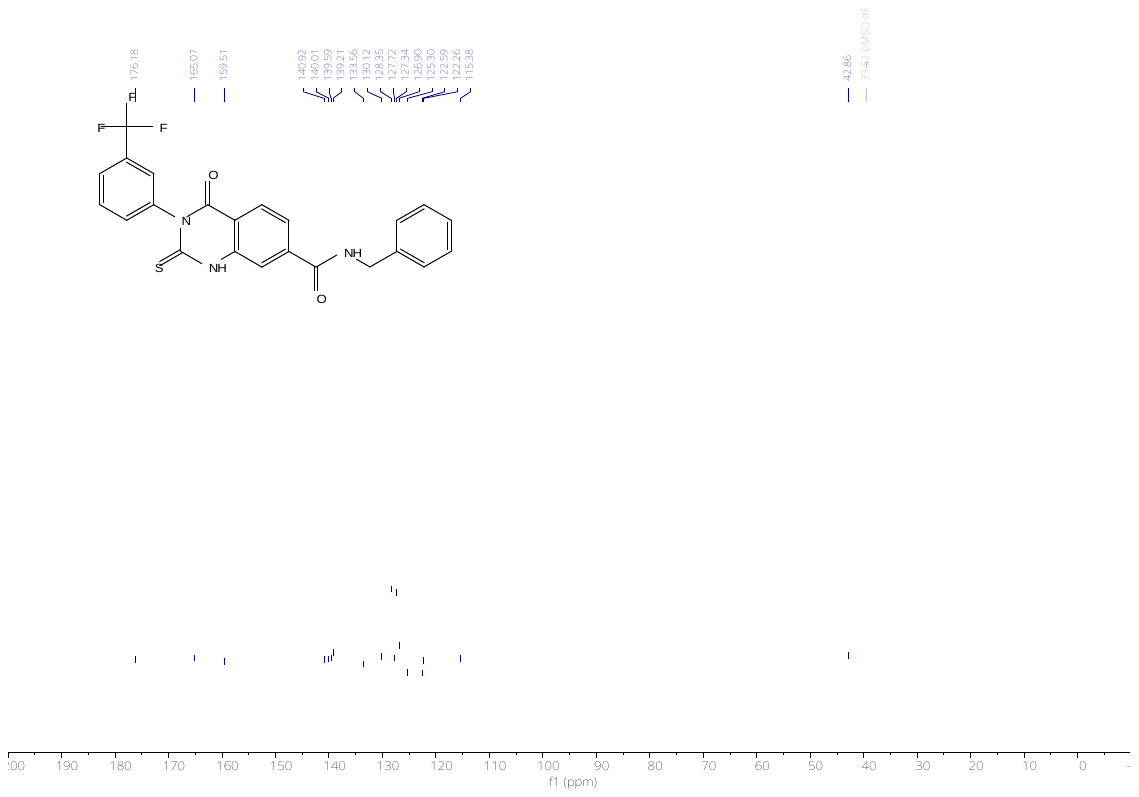

**^19^F NMR**

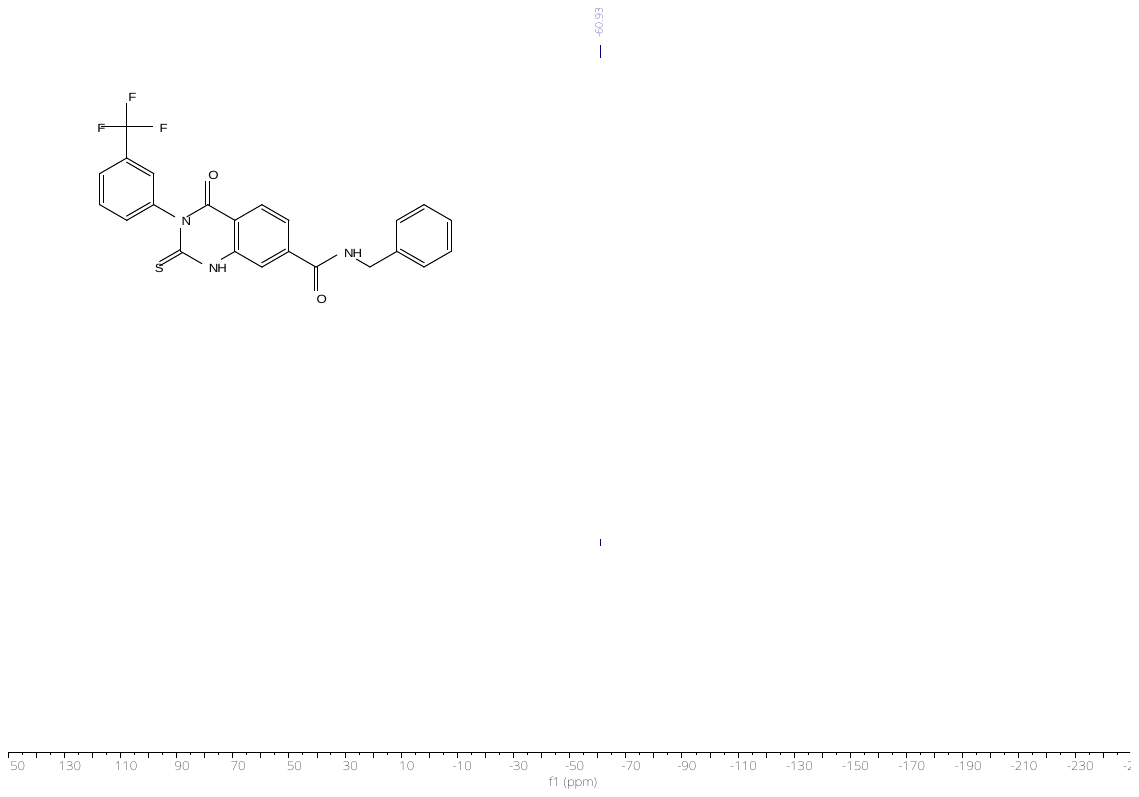

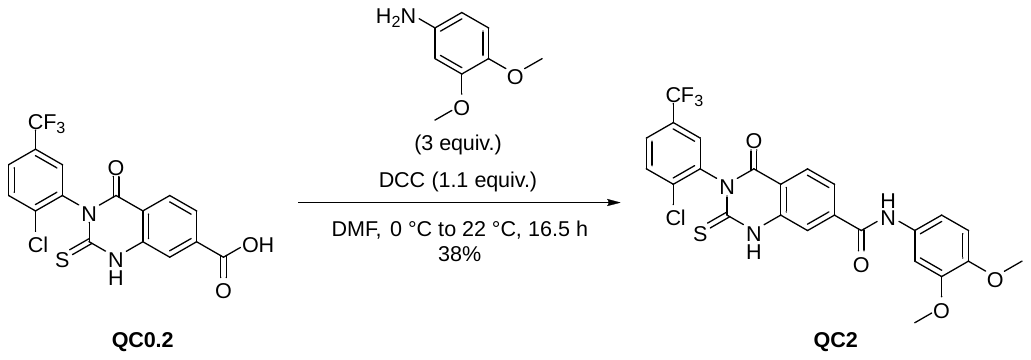

**3-(2-chloro-5-(trifluoromethyl)phenyl)-*N*-(3,4-dimethoxyphenyl)-4-oxo-2-thioxo-1,2,3,4-tetrahydroquinazoline-7-carboxamide (QC2).** To a stirred solution of 3-(2-chloro-5-(trifluoromethyl)phenyl)-4-oxo-2-thioxo-1,2,3,4-tetrahydroquinazoline-7-carboxylic acid (0.1169 g, 0.2917 mmol) in dry DMF (2 mL) at 0 °C under an argon atmosphere DCC (0.077 g, 0.3732 mmol) was added and the mixture was stirred for 30 minutes. Then, 3,4-Dimethoxyaniline (0.133 g, 0.8683 mmol) was added, and the mixture was stirred for 16 h whilst warming to 22 °C. The volatiles were removed, and the residue taken up in a mixture of acetonitrile/ water with little TFA. The crude was purified by reverse phase flash column chromatography (water + 0.1% TFA/ MeCN + 0.1% TFA gradient from 1:0 to 0:1), to give the desired compound in 38% yield (0.05885 g, 0.1098 mmol). Yellow solid: **^1^H NMR** (300 MHz, DMSO-*d*_6_) δ 13.50 (s, 1H), 10.45 (s, 1H), 8.12 (d, *J* = 8.1 Hz, 2H), 8.03 – 7.81 (m, 4H), 7.48 (d, *J* = 2.0 Hz, 1H), 7.35 (dd, *J* = 8.7, 2.0 Hz, 1H), 6.97 (d, *J* = 8.8 Hz, 1H), 3.77 (s, 3H), 3.76 (s, 3H). **^13^C NMR** (101 MHz, DMSO) δ 175.65, 164.16, 159.19, 148.96, 146.03, 142.53, 140.15, 137.84, 136.88, 136.87, 132.64, 131.46, 129.29 (q, *J* = 32.9 Hz), 129.18 (q, *J* = 7.6, 3.8 Hz), 127.76 – 127.45 (m), 128.30, 123.96 (q, *J* = 272.3 Hz), 123.46, 117.86, 116.17, 112.98, 112.35, 105.99, 56.18, 55.90. **^19^F NMR** (282 MHz, DMSO) δ -61.01. **HRMS** (ESI) calculated for [M+H]^+^ C_24_H_17_ClF_3_N_3_O_4_S 536.0653, found 536.0620.

**^1^H NMR**

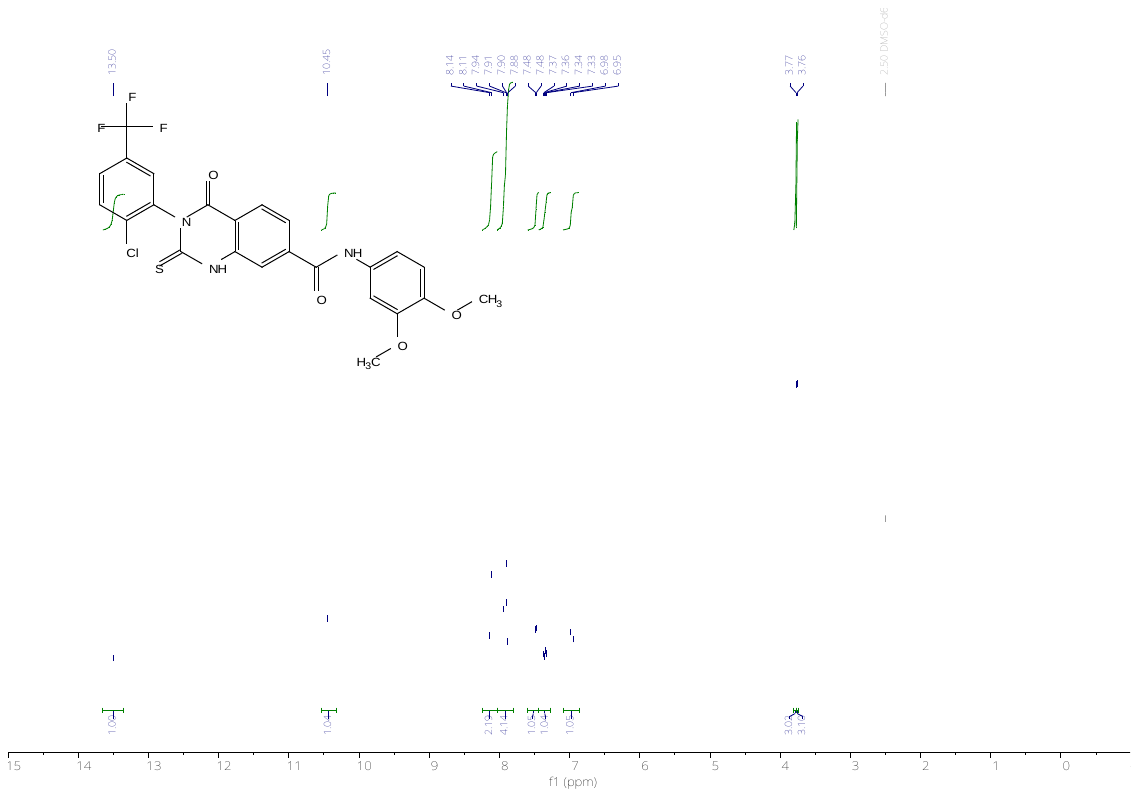

**^13^C NMR**

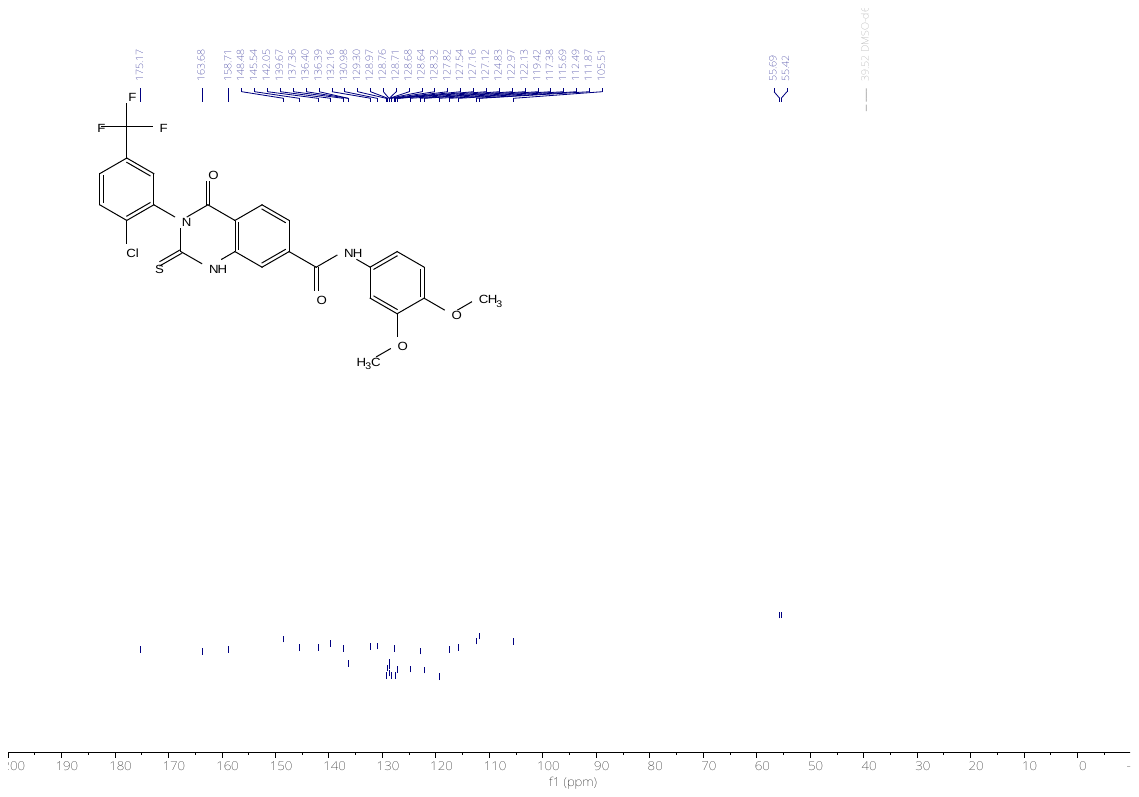

**^19^F NMR**

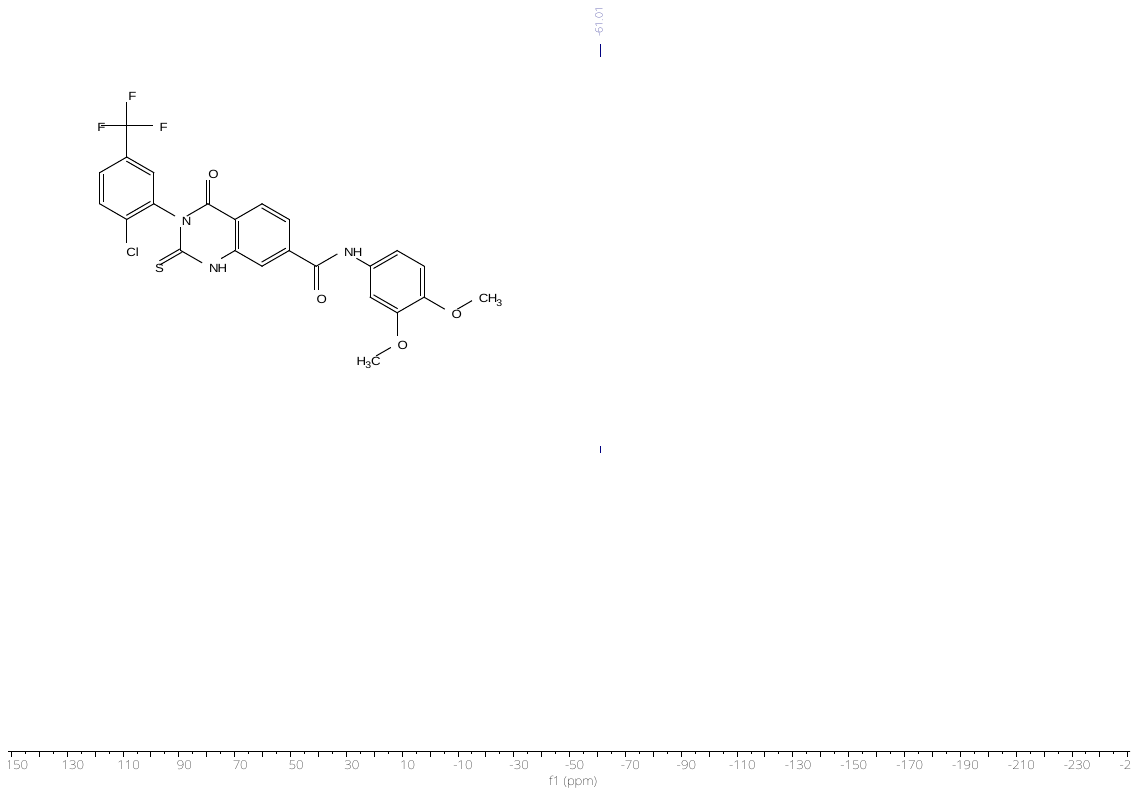

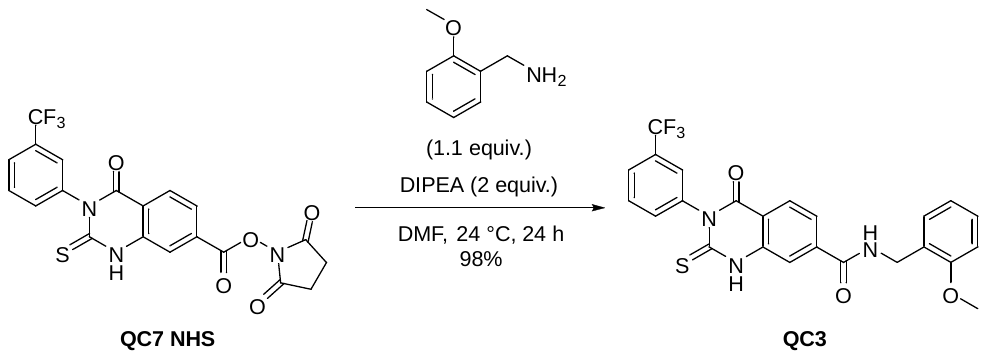

***N*-(2-methoxybenzyl)-4-oxo-2-thioxo-3-(3-(trifluoromethyl)phenyl)-1,2,3,4-tetrahydroquinazoline-7-carboxamide (QC3).** Prepared according to general procedure I: 98% yield (0.10493 g, 0.21614 mmol). White solid: **^1^H NMR** (400 MHz, DMSO-*d*_6_) δ 13.26 (s, 1H), 9.16 (t, *J* = 5.8 Hz, 1H), 8.04 (d, *J* = 8.3 Hz, 1H), 7.91 (d, *J* = 1.6 Hz, 1H), 7.80 (dd, *J* = 9.9, 1.7 Hz, 3H), 7.74 (t, *J* = 7.8 Hz, 1H), 7.67 (d, *J* = 7.8 Hz, 1H), 7.30 – 7.19 (m, 2H), 7.02 (dd, *J* = 8.2, 1.1 Hz, 1H), 6.93 (td, *J* = 7.4, 1.1 Hz, 1H), 4.48 (d, *J* = 5.7 Hz, 2H), 3.84 (s, 3H). **^13^C NMR** (101 MHz, DMSO) δ 176.18, 165.22, 159.52, 156.66, 141.02, 140.02, 139.58, 133.57, 130.12, 129.75 (q, *J* = 32.1 Hz), 128.13, 127.68, 127.51, 126.51 – 126.20 (m), 126.40, 125.30, 125.22 – 124.87 (m), 122.60, 122.36, 120.14, 115.38, 110.57, 55.38, 37.93. **^19^F NMR** (282 MHz, DMSO) δ -60.93. **HRMS** (ESI) calculated for [M+H]^+^ C_24_H_19_F_3_N_3_O_3_S 486.1094, found 486.1081.

**^1^H NMR**

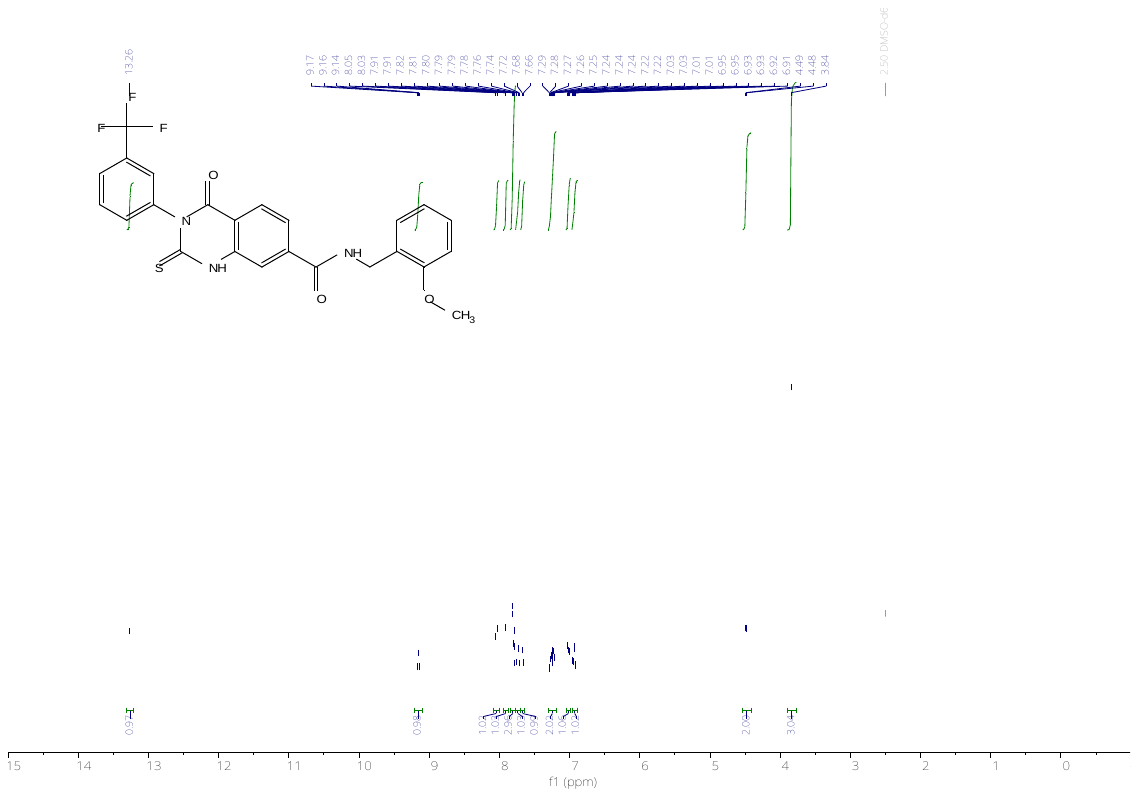

**^13^C NMR**

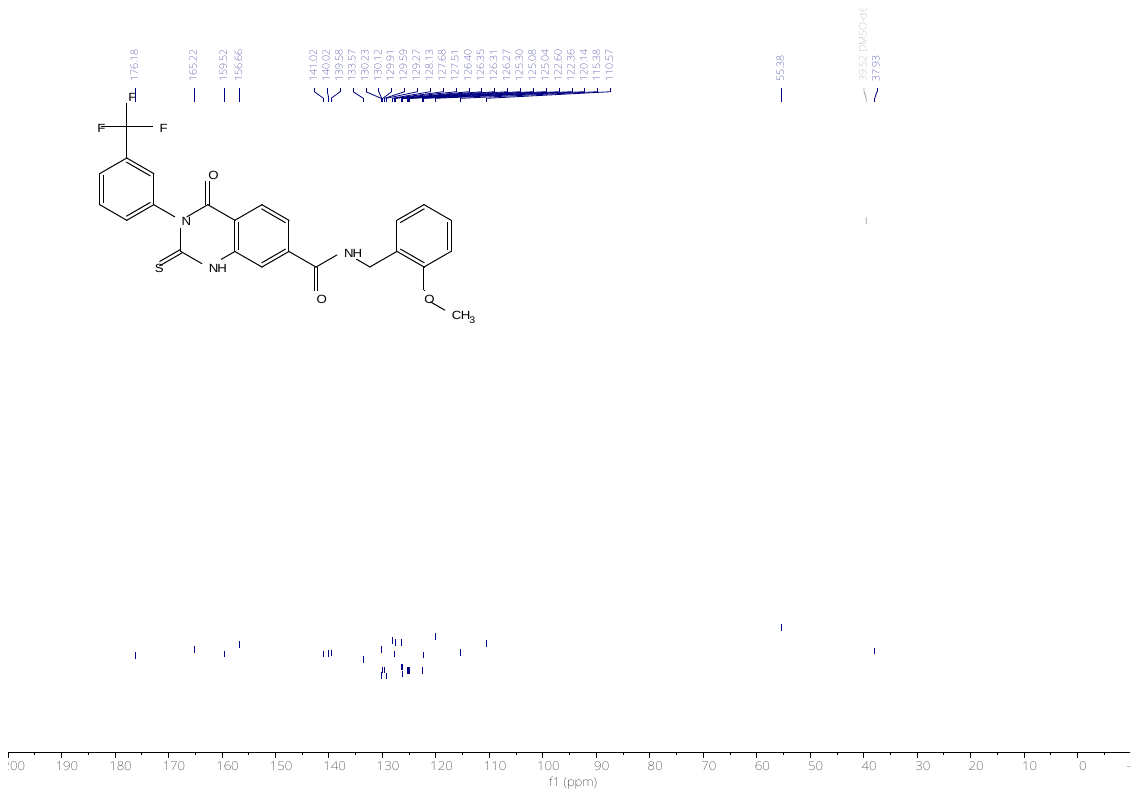

**^19^F NMR**

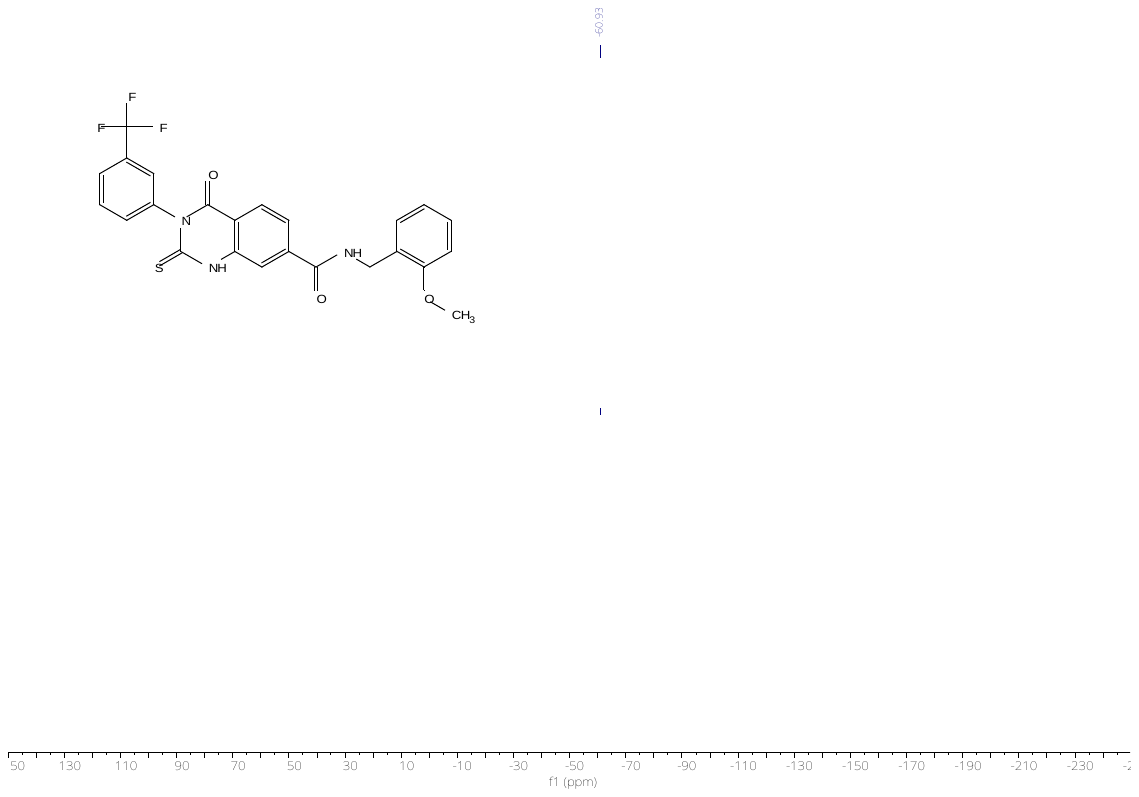

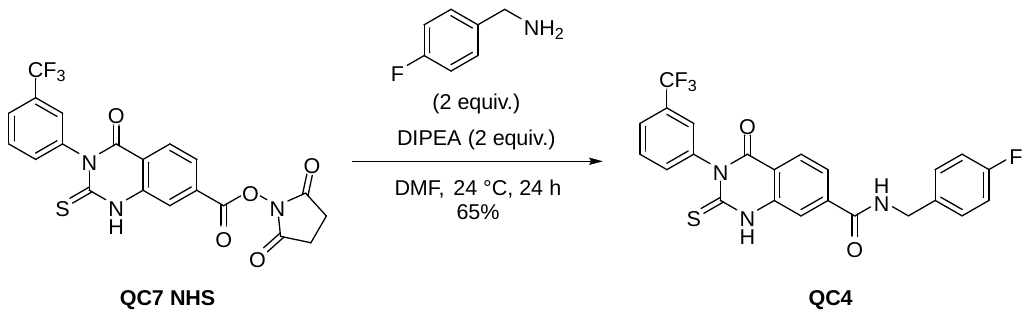

***N*-(4-fluorobenzyl)-4-oxo-2-thioxo-3-(3-(trifluoromethyl)phenyl)-1,2,3,4-tetrahydroquinazoline-7-carboxamide (QC4).** Prepared according to general procedure I: 65% yield (0.06704 g, 0.14160 mmol). Yellowish solid: **^1^H NMR** (300 MHz, DMSO-*d*_6_) δ 13.26 (s, 1H), 9.36 (t, *J* = 6.0 Hz, 1H), 8.04 (d, *J* = 8.2 Hz, 1H), 7.90 (d, *J* = 1.5 Hz, 1H), 7.84 – 7.61 (m, 5H), 7.39 (dd, *J* = 8.5, 5.7 Hz, 2H), 7.24 – 7.12 (m, 2H), 4.48 (d, *J* = 5.8 Hz, 2H). **^13^C NMR** (101 MHz, DMSO) δ 176.18, 165.05, 162.44, 160.03, 159.50, 140.81, 140.00, 139.59, 135.56 – 135.30 (m), 133.82 – 133.37 (m), 130.13, 129.90, 129.58, 129.43, 129.35, 127.73, 126.32 (q, *J* = 3.6 Hz), 125.30, 125.21 – 124.88 (m), 122.59, 122.24, 115.37, 115.18, 114.97, 42.19. **^19^F NMR** (282 MHz, DMSO) δ -60.93. **HRMS** (ESI) calculated for [M+H]^+^ C_23_H_16_F_4_N_3_O_2_S 474.0894, found 474.0881.

**^1^H NMR**

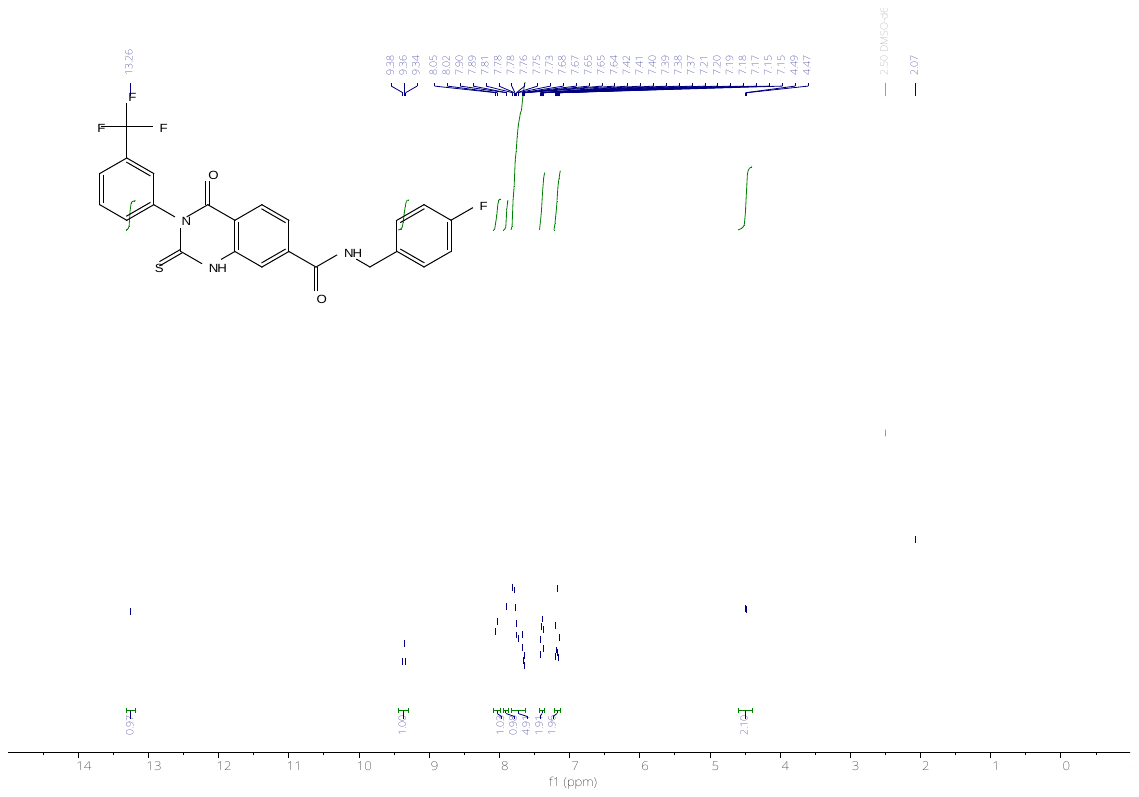

**^13^C NMR**

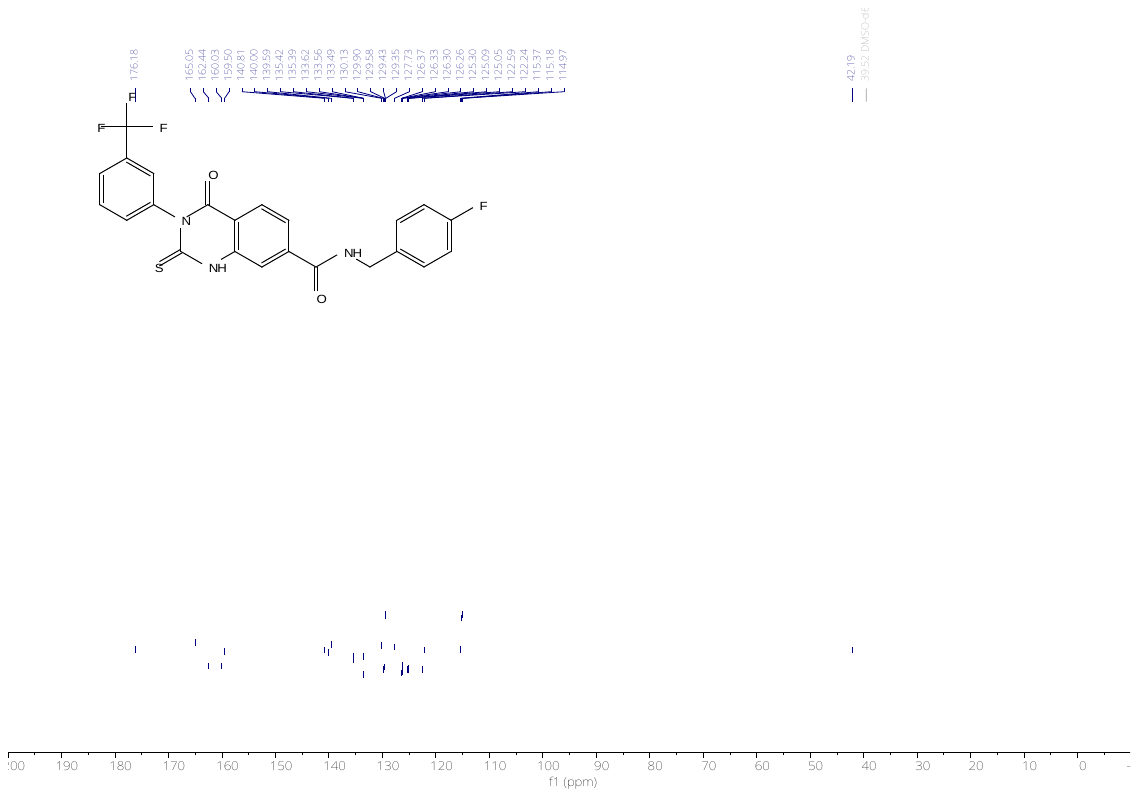

**^19^F NMR**

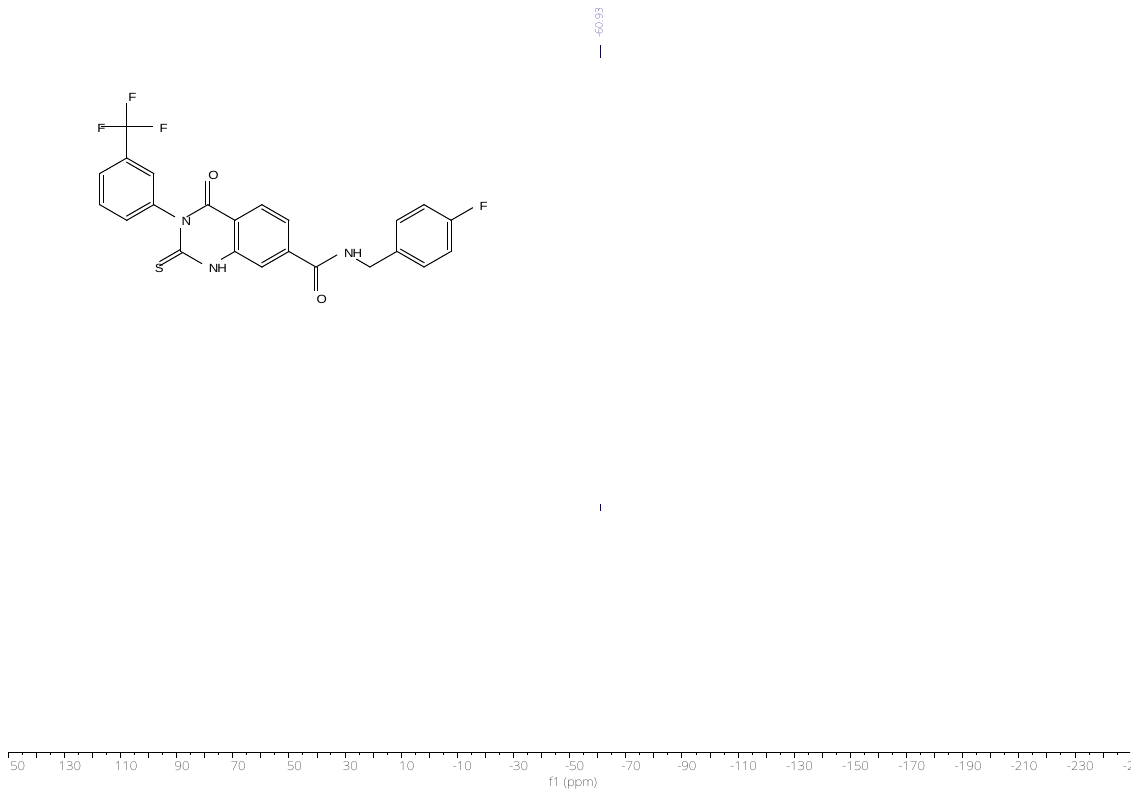

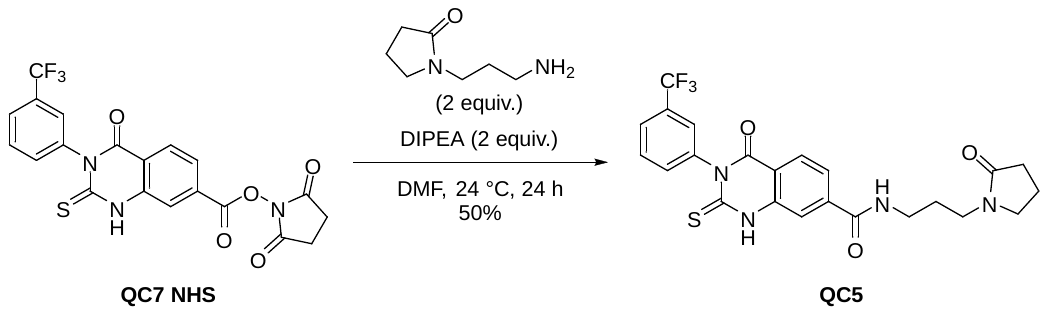

**4-oxo-*N*-(3-(2-oxopyrrolidin-1-yl)propyl)-2-thioxo-3-(3-(trifluoromethyl)phenyl)-1,2,3,4-tetrahydro-quinazoline-7-carboxamide (QC5).** Prepared according to general procedure I: 50% yield (0.05538 g, 0.11299 mmol). White solid: **^1^H NMR** (400 MHz, DMSO-*d*_6_) δ 13.26 (s, 1H), 8.75 (t, *J* = 5.6 Hz, 1H), 8.03 (d, *J* = 8.2 Hz, 1H), 7.87 (d, *J* = 1.5 Hz, 1H), 7.80 (d, *J* = 8.8 Hz, 2H), 7.74 (d, *J* = 8.1 Hz, 1H), 7.73 – 7.70 (m, 1H), 7.66 (d, *J* = 7.9 Hz, 1H), 3.36 (t, *J* = 7.0 Hz, 2H), 3.29 – 3.22 (m, 4H), 2.23 (t, *J* = 8.0 Hz, 2H), 1.92 (q, *J* = 7.7 Hz, 2H), 1.74 (p, *J* = 7.1 Hz, 2H). **^13^C NMR** (101 MHz, DMSO) δ 176.17, 173.99, 164.90, 159.51, 141.09, 140.01, 139.61, 133.56, 130.12, 129.74 (q, *J* = 32.1 Hz), 127.67, 126.49 – 126.08 (m), 125.30, 125.18 – 124.94 (m), 122.59, 122.12, 117.98, 115.22, 46.34, 37.10, 30.45, 26.68, 17.52. **^19^F NMR** (282 MHz, DMSO) δ -60.93. **HRMS** (ESI) calculated for [M+H]^+^ C_23_H_22_F_3_N_4_O_3_S 491.1359, found 491.1351.

**^1^H NMR**

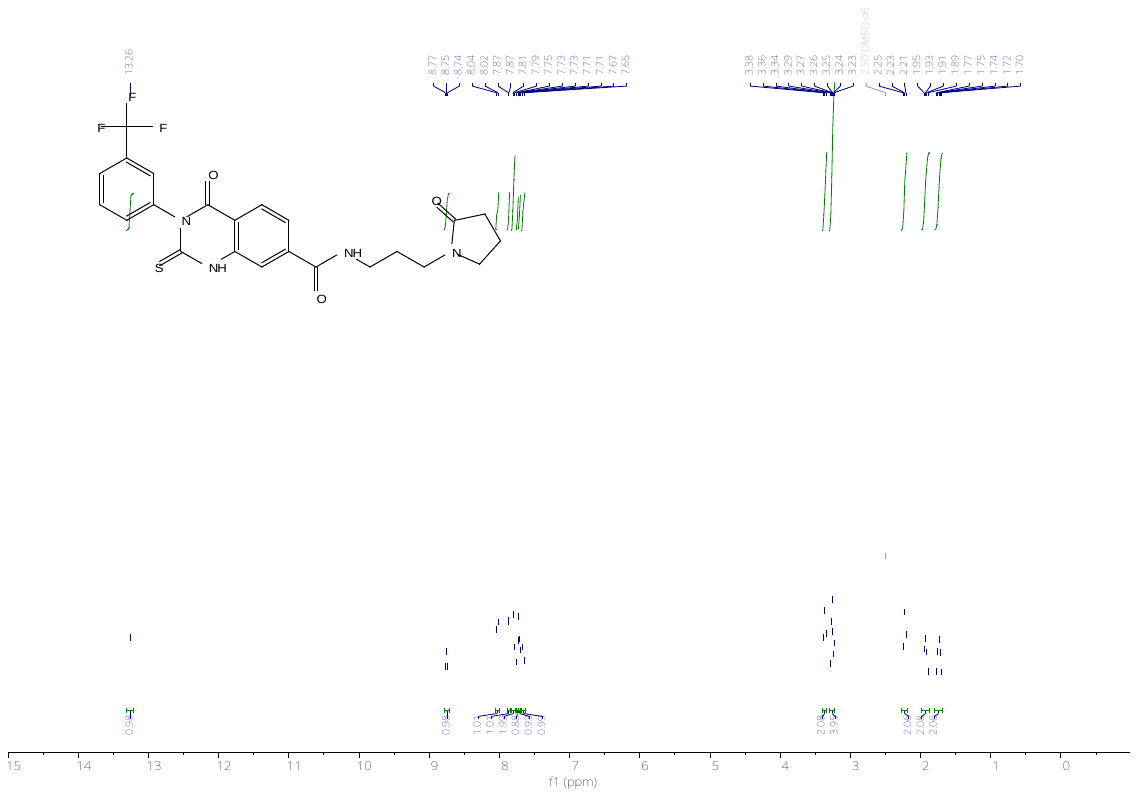

**^13^C NMR**

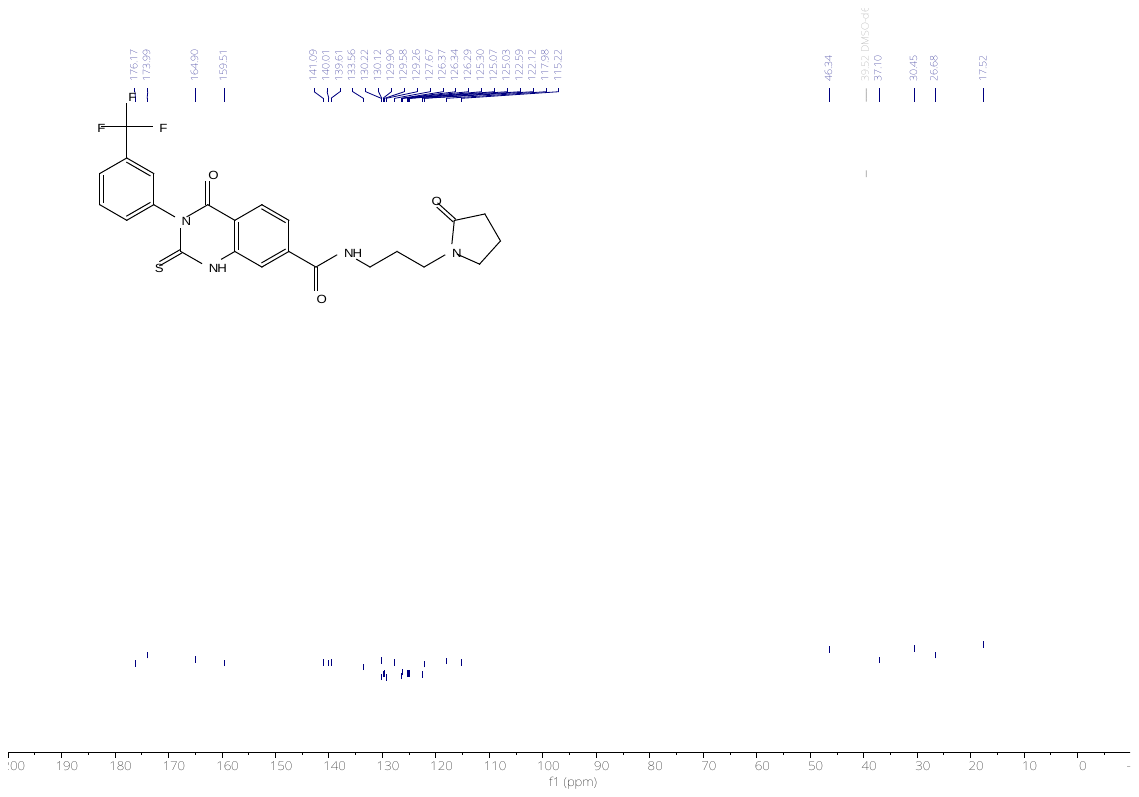

**^19^F NMR**

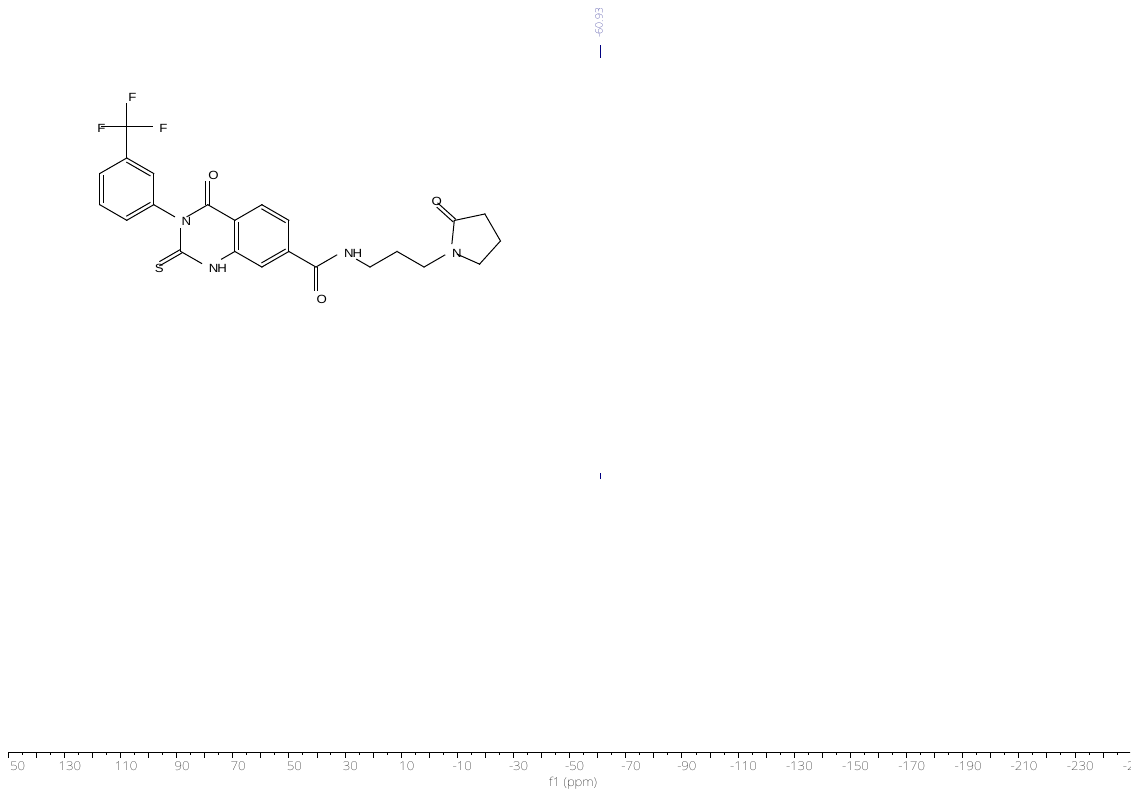

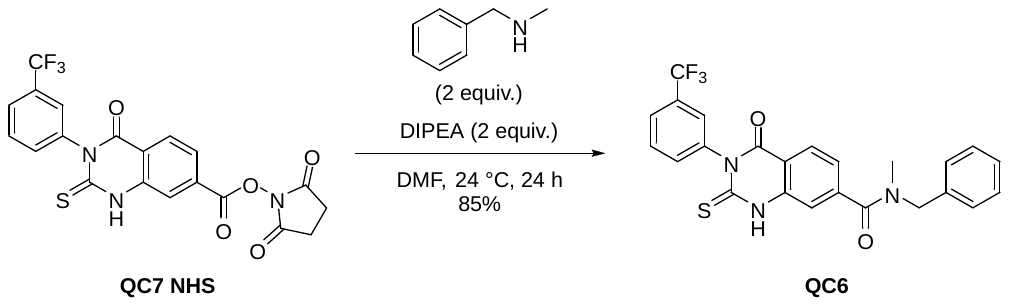

***N*-benzyl-*N*-methyl-4-oxo-2-thioxo-3-(3-(trifluoromethyl)phenyl)-1,2,3,4-tetrahydroquinazoline-7-carboxamide (QC6).** Prepared according to general procedure I: 85% yield (0.08755 g, 0.18648 mmol). Yellow solid: Mixture of amide rotamers, **^1^H NMR** (400 MHz, DMSO-*d*_6_) δ 13.18 (s, 1H), 8.08 – 7.95 (m, 1H), 7.86 – 7.69 (m, 3H), 7.69 – 7.57 (m, 1H), 7.52 – 7.26 (m, 6H), 7.20 (d, *J* = 7.5 Hz, 1H), 4.78 – 4.41 (m, 2H), 2.98 – 2.79 (m, 3H). **^13^C NMR** (101 MHz, DMSO) δ 176.14, 169.25, 168.67, 159.45, 143.06, 142.92, 139.97, 139.63, 137.00, 136.32, 133.53, 130.15, 129.91, 129.59, 129.27, 128.79, 128.69, 128.00, 127.71, 127.54, 127.38, 126.77, 126.27, 125.29, 125.07, 122.58, 122.38, 122.08, 116.77, 113.81, 113.49, 54.90, 53.82, 49.82, 36.70, 32.52. **^19^F NMR** (376 MHz, DMSO) δ -60.95, -60.97. **HRMS** (ESI) calculated for [M+H]^+^ C_24_H_19_F_3_N_3_O_2_S 470.1145, found 470.1136.

**^1^H NMR**

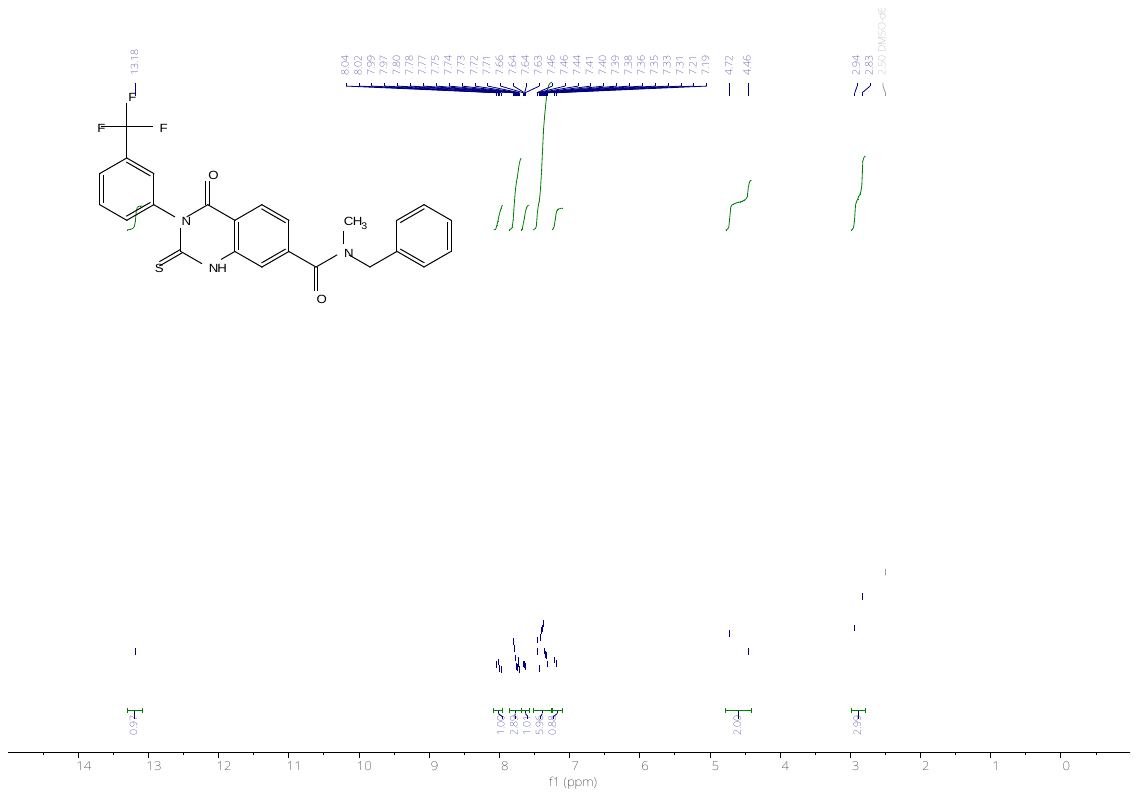

**^13^C NMR**

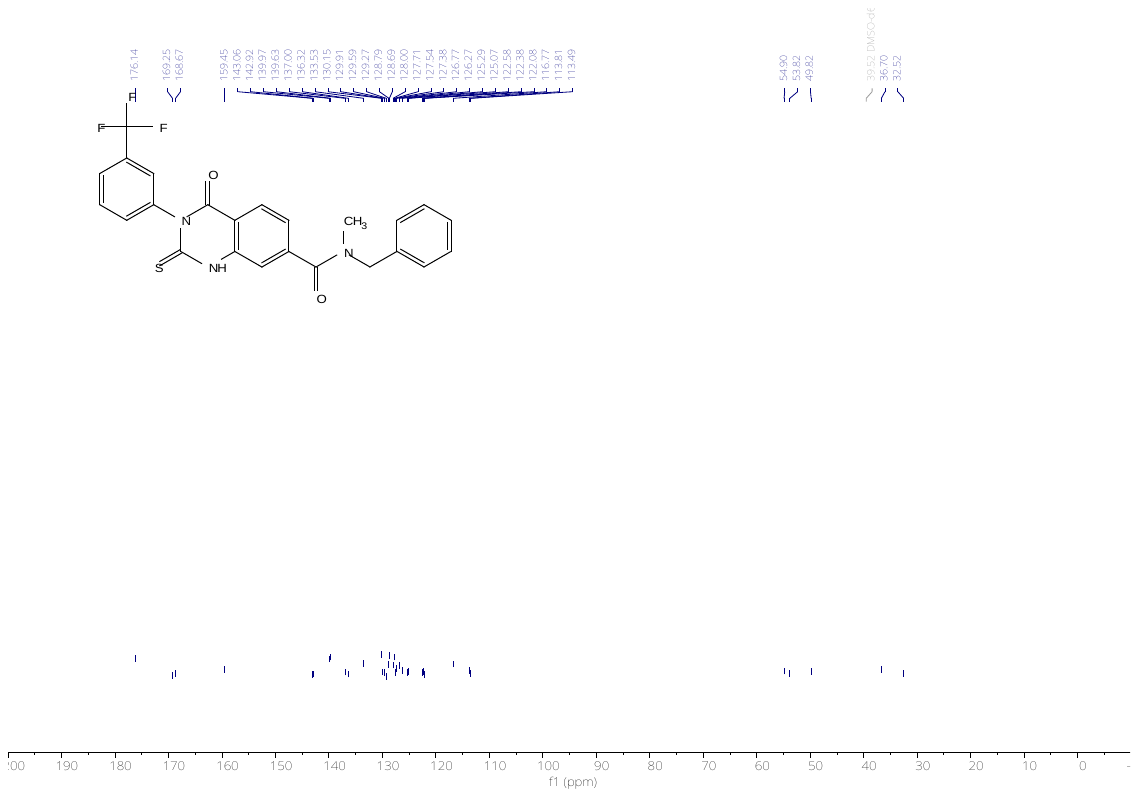

**^19^F NMR**

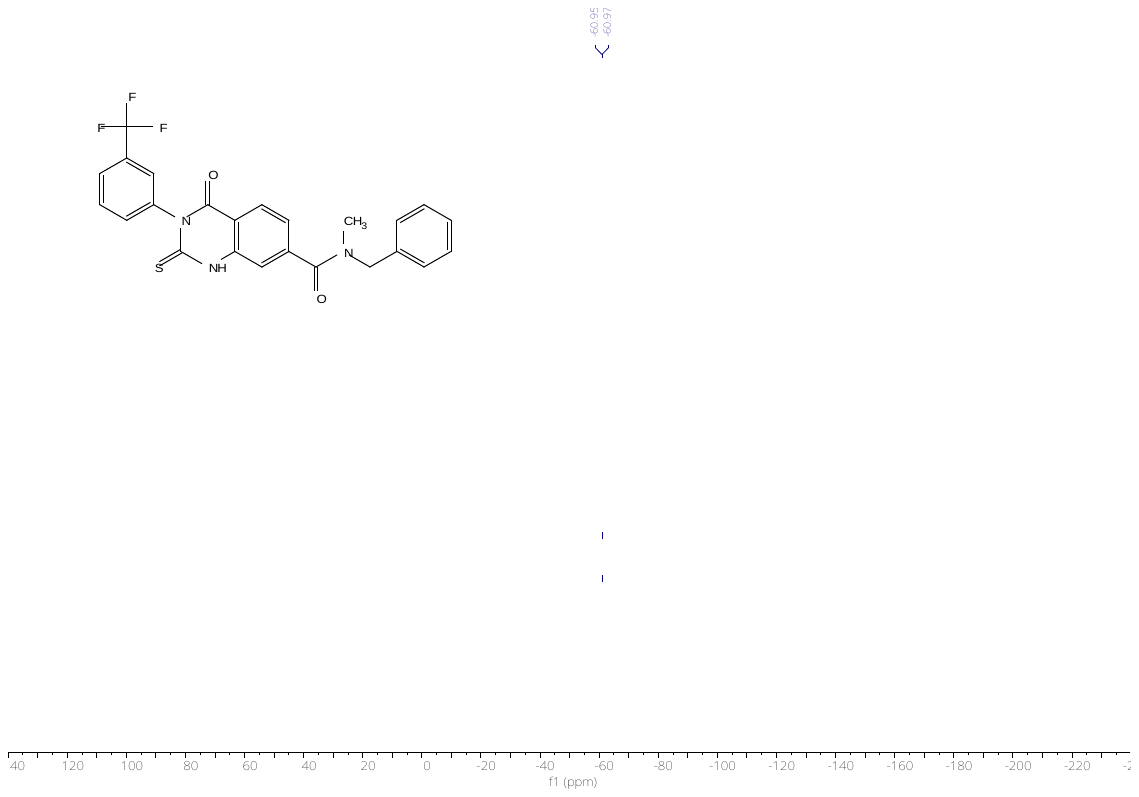
