## Supplementaries for "Investigation of the threonine metabolism of *Echinococcus multilocularis*: the threonine dehydrogenase as a potential drug target in alveolar echinococcosis": S3 File.txt

//This macro aims to automate spheroid size measurement in three-dimensional cell culture by approximation from measurements from images of a two-dimensional plane
//It requires an input folder with image files only
//It detects spheroids with manual input from the user and outputs images with the outline/s of the determined spheroid/s
//It measures the detected spheroid's area 
//It measures the detected spheroid's mean radial distance by measuring 360 radial distances from the detected spheroid’s centroid to its approximated border and calculates the mean value
requires("1.53d");
//Expandable arrays are used in the script
setOption("ExpandableArrays", true);

//Returns an asterisk progress bar from the ratio of p1 to p2
//Used in script for visualizing progress of multiple image processing
function getBar(p1, p2) {
       n = 20;
       bar1 = "--------------------";
       bar2 = "********************";
       index = round(n*(p1/p2));
       if (index<1) index = 1;
       if (index>n-1) index = n-1;
       return substring(bar2, 0, index) + substring(bar1, index+1, n);
}

//This function isolates a spheroid in an already open image by deleting everything except a circle selection with the given parameters
//The deleted space is filled with the mean pixel value of the original space so as to minimize incorrect detection in further processing if space was an absolute 0 or 1
function isolateSpheroid(xcentroid,ycentroid,diameter)
{
	run("Set Scale...", "distance=0");
	run("Select None");
	size = diameter/2 + 50;
//Determines limits for width and height in case diameter that is input exceeds distance from centroid to image borders and if so sets size to limit instead of diameter that was input
	widthLim = getWidth()-xcentroid;
	heightLim = getHeight()-ycentroid;
	if (size>xcentroid)
		size = xcentroid;
	if (size>ycentroid)
		size = ycentroid;
	if (size>widthLim)
		size = widthLim;
	if (size>heightLim)
		size = heightLim;
	makeOval(xcentroid-size, ycentroid-size, 2*size, 2*size);
	run("Make Inverse");
	getStatistics(area,mean,min,max,std);
	setBackgroundColor(mean, mean, mean);
	run("Clear", "slice");
	run("Select None");
};

//This is the function to measure the mean radial distance of the active image
//res is the angular resolution
//The image file must already be open and contain the selection or overlay to be measured
//Returns the value of the mean radial distance
function radialDistance(res)
{
	//Divides the 360 degree rotation into n steps
	n = 360/res;

	//Stores sum of squared measured radii for later area calculation
	sqsumr = 0;

	setOption("BlackBackground", false);
	run("Create Mask");
	run("Create Selection");

	//Gets coordinates of centroid of ellipse from which radii will be drawn
	run("Set Scale...", "distance=0 known=0 pixel=1 unit=pixel");
	xcentroid = getValue("X");
	ycentroid = getValue("Y");
	run("Clear Results");
	run("Select None");
	run("Remove Overlay");
	run("Find Edges");

	//Gets the px dimensions of the image
	width = getWidth();
	height = getHeight();

	//Calculates angles at which elongated radii would touch the image corners
	corner1 = atan((ycentroid)/(width-xcentroid))*180/PI;
	corner2 = 180-atan((ycentroid)/(xcentroid))*180/PI;
	corner3 = atan((height-ycentroid)/(xcentroid))*180/PI+180;
	corner4 = 360-atan((height-ycentroid)/(width-xcentroid))*180/PI;

	//Array for storage of radii
	radii = newArray(n);

	//Draws lines from centroid all around and measures radii
	for (i=0; i<n; i++)
	{ 
		angle = i*res;
		if(angle == 0||angle == 180)
		{
			y = ycentroid;
			x = width-width*angle/180;
		}
		else if(angle == 90||angle == 270)
		{
			x = xcentroid;
			y = height-height*(270-angle)/180;
		}
		else
		{
			if(angle < corner1 || angle > corner4)
			{
				x = width;
				y = ycentroid-(width-xcentroid)*tan(angle*PI/180);
			}
			else if(angle < corner2)
			{
				y = 0;
				x = xcentroid + ycentroid/tan(angle*PI/180);
			}
			else if(angle < corner3)
			{
				x = 0;
				y = ycentroid + xcentroid*tan(angle*PI/180);
			}
			else if(angle < corner4)
			{
				y = height;
				x = xcentroid-(height-ycentroid)/tan(angle*PI/180);
			};
		};
		makeLine(xcentroid,ycentroid,x,y);
		prof = getProfile();
tolPeak = 50;
		//tolPeak is the amplitude tolerance to consider separate peaks and can be modified depending on the images for the assay
		peaks = Array.findMaxima(prof, tolPeak);


		//Sets radius as not a number in case no maximum was found and the peaks array is empty
		radii[i] = 0/0;
		if (peaks.length != 0)
			radii[i] = peaks[0];
		
		sqsumr = sqsumr + radii[i]*radii[i];
	};

	//Gets statistics from array of radii to calculate minimum maximum and mean
	Array.getStatistics(radii, min, max, mean, std);

	//Approximates area by summation of n triangles
	sqsumr = sqsumr*tan((res*PI/180)/2);

	return mean;
};


//This is the actual spheroid detection and measurement applied to every image
//Adapted and expanded from Ivanov, DP /2015/ Assessment of drug-loaded nanoparticles in a 3D in vitro brain tumour model /Doctoral dissertation, University of Nottingham/
//Analyzes the already open input image file and ends up with an overlay of the detected spheroid outline on the edited image
//The thresholdMode parameter must specify Default or Manual
function action(inputFolder,filename,thresholdMode,enhanceCont,lowerThresh,upperThresh)
{
//minCirc is the minimum circularity parameter and can be modified depending on the assay implementation
	minCirc = 0.6;

if (thresholdMode == "Default")
	{
		run("Enhance Contrast", "saturated=" + enhanceCont + " normalize");
		//Uses Default thresholding algorithm 
		setAutoThreshold("Default");
		setOption("BlackBackground", false);
		run("Convert to Mask");
	}
	else if (thresholdMode == "ModifManual")
	{
		run("Revert");
		run("Set Scale...", "distance=0");
		run("8-bit");
		origImg = getImageID();
		percentSat = enhanceCont;
		enhanceCont = percentSat;
		run("Enhance Contrast", "saturated=" + percentSat + " normalize");

		ovalWidth = getWidth()/2;
		setBatchMode("exit and display");
		getCursorLoc(x, y, z, flags);
		
		x2=-1; y2=-1;
		run("Original Scale");
		zoom = getZoom();
		while (!isKeyDown("space") && !isKeyDown("shift"))
		{
			getCursorLoc(x, y, z, flags);
			zoom2= getZoom();
			if (zoom2 < zoom)
			{
				run("Original Scale");
				ovalWidth = ovalWidth - 100;
				wait(50);
			};
			if (zoom2 > zoom)
			{
				run("Original Scale");
				ovalWidth = ovalWidth + 50;
				wait(50);
			};
			if (x!=x2 || y!=y2 || zoom != zoom2)
			{
				makeOval(x-ovalWidth/2, y-ovalWidth/2, ovalWidth, ovalWidth);
				if (zoom == zoom2) wait(10);
			};
			x2=x; y2=y;
		};
		setKeyDown("none");		
	
		setBatchMode("hide");
	
		numPix = 10;
		imgWidth = getWidth(); imgHeight = getHeight();
		needToExpand = (x-ovalWidth/2<10 || x+ovalWidth/2>imgWidth-10 || y+ovalWidth/2>imgHeight-10 || y-ovalWidth/2<10);
		if (needToExpand)
		{
			oldImgWidth = getWidth();
			oldImgHeight = getHeight();
			run("Canvas Size...", "width=" + (oldImgWidth+ovalWidth) + " height=" + (oldImgHeight+ovalWidth) + " position=Center zero");
			imgWidth = getWidth(); imgHeight = getHeight();
			x = x + ovalWidth/2;
			y = y + ovalWidth/2;
			makeOval(x-ovalWidth/2, y-ovalWidth/2, ovalWidth, ovalWidth);
		};
		diagDists = newArray(sqrt(x*x+y*y),sqrt((imgWidth-x)*(imgWidth-x)+(imgHeight-y)*(imgHeight-y)),sqrt((imgWidth-x)*(imgWidth-x)+y*y),sqrt(x*x+(imgHeight-y)*(imgHeight-y)));
		Array.sort(diagDists);
		maxSize = diagDists[diagDists.length-1];

		run("Make Band...", "band=" + numPix);
		run("Copy");
		newImage("TempRing", "8-bit white", ovalWidth+2*numPix, ovalWidth+2*numPix, 1);
		run("Paste");

		run("Scale...", "x=- y=- width=" + (ovalWidth + 1.8*2*numPix) + " height=" + (ovalWidth + 1.8*2*numPix) + " interpolation=Bilinear average create title=TempRing2");
		makeOval(0.5*(getWidth()-ovalWidth)-0.9*numPix, 0.5*(getHeight()-ovalWidth)-0.9*numPix, ovalWidth + 0.9*2*numPix, ovalWidth + 0.9*2*numPix);
		run("Make Inverse");
		run("Copy");
		close("TempRing2");
		selectImage("TempRing");
		run("Select None");
		run("Canvas Size...", "width=" + (ovalWidth + 1.8*2*numPix) + " height=" + (ovalWidth + 1.8*2*numPix) + " position=Center zero");
		run("Paste");
		numPix = 1.8*numPix;

		while (getWidth() <= 2*maxSize)
		{
			makeOval(0.5*(getWidth()-ovalWidth), 0.5*(getWidth()-ovalWidth), ovalWidth, ovalWidth);
			run("Make Inverse");
			run("Scale...", "x=- y=- width=" + (ovalWidth + 1.8*2*numPix) + " height=" + (ovalWidth + 1.8*2*numPix) + " interpolation=Bilinear average create title=TempRing2");
			makeOval(0.5*(getWidth()-ovalWidth)-0.9*numPix, 0.5*(getHeight()-ovalWidth)-0.9*numPix, ovalWidth + 0.9*2*numPix, ovalWidth + 0.9*2*numPix);
			run("Make Inverse");
			run("Copy");
			close("TempRing2");
			selectImage("TempRing");
			run("Select None");
			run("Canvas Size...", "width=" + (ovalWidth + 1.8*2*numPix) + " height=" + (ovalWidth + 1.8*2*numPix) + " position=Center zero");
			run("Paste");
			numPix = 1.8*numPix;
		};
		
		makeRectangle(getWidth()/2-x, getHeight()/2-y, imgWidth, imgHeight);
		run("Crop");
		makeOval(x-ovalWidth/2-3, y-ovalWidth/2-3, ovalWidth+6, ovalWidth+6);
		run("Make Inverse");
		getStatistics(area, mean, min, max, std, histogram);
		BGColor = mean;
		run("Copy");
		close("TempRing");
		selectImage(origImg);
		run("Paste");
		run("Select None");
		
		res = 1;
		xcentroid = x;
		ycentroid = y;
		if (needToExpand)
		{
			//run("Canvas Size...", "width=" + (imgWidth+ovalWidth) + " height=" + (imgHeight+ovalWidth) + " position=Center zero");
			//xcentroid = x + ovalWidth/2;
			//ycentroid = y + ovalWidth/2;
			makeRectangle(ovalWidth/2, ovalWidth/2, oldImgWidth, oldImgHeight);
			run("Make Inverse");
			setBackgroundColor(BGColor, BGColor, BGColor);
			run("Clear", "slice");
			run("Select None");
		};
		//imgWidth = getWidth(); imgHeight = getHeight();
		xs = newArray();	ys = newArray();

		//Draws lines from centroid all around and stores coordinates of minima
		for (i=0; i*res<360; i++)
		{ 
			angle = i*res;
			x = xcentroid+0.5*ovalWidth*cos(angle*PI/180);
			y = ycentroid-0.5*ovalWidth*sin(angle*PI/180);
			makeLine(xcentroid,ycentroid,x,y);
			prof = getProfile();
			peaks = Array.findMinima(prof, 5);
			if (peaks.length != 0)
			{
				darkVal = prof[peaks[0]];
				prof2 = Array.slice(prof,peaks[0]);
				peaks2 = Array.findMaxima(prof2, 5);
				if (peaks2.length != 0)
				{
					lightVal = prof2[peaks2[0]];
					//threshVal = (darkVal+lightVal)/2;
					threshVal = darkVal+(lightVal-darkVal)*0.25;
					for (ii=peaks[0]; ii<prof.length; ii++)
					{
						if (prof[ii]>=threshVal) break;
					};
					if (ii < prof.length)
					{
						xs = Array.concat(xs,xcentroid+ii*cos(angle*PI/180));
						ys = Array.concat(ys,ycentroid-ii*sin(angle*PI/180));
						if (needToExpand)
						{
							xs[xs.length-1] = xs[xs.length-1]-ovalWidth/2;
							ys[ys.length-1] = ys[ys.length-1]-ovalWidth/2;
						};
					};
				};
			};
		};
		if (needToExpand)
		{
			makeRectangle(ovalWidth/2, ovalWidth/2, oldImgWidth, oldImgHeight);
			run("Crop");
		};
		makeSelection(1, xs, ys);
		setForegroundColor(0, 0, 0);
		run("Fill", "slice");
		setBackgroundColor(255, 255, 255);
		run("Clear Outside");
		run("Select None");
		run("Convert to Mask");
	}
	else
	{
		waitForUser("Error: This threshold mode has not been implemented yet.");
	};

	//Gets the ratio between black (spheroid) and white (background) pixels. Assuming a single spheroid, the black to white ratio allows us to estimate size of the spheroid
	getHistogram(0,hist,256);
	ratio = hist[255]/hist[0];

	minSize = PI*ovalWidth*ovalWidth/4/4;

	//If there are more pixels detected as spheroid(black) than background(white) then the spheroid has not been detected due to variations in background
	//if (ratio>1)
	//{
		// reverts the image, subtracts the background and proceeds as normal
		//run("Revert");
		//run("8-bit");
		//percentSat = enhanceCont;
		//enhanceCont = percentSat;
		//run("Enhance Contrast", "saturated=" + percentSat + " normalize");
		// Subtract Background is not used in the default function because it can lead to merging of spheroids and debris or it can remove the core of the spheroid leaving a very thin interrupted edge. In certain cases where the edges of a spheroid are very bright removing the background can give better results.
		//run("Subtract Background...", "rolling=50 light");
		//setAutoThreshold("Yen");
		//setOption("BlackBackground", false);
		//run("Convert to Mask");
		//run("Remove Outliers...", "radius=15 threshold=0 which=Dark");
		//getHistogram(0,hist,256);
		//ratio = hist[255]/hist[0];
	//};
	
	//The strategy here is to act differently according to spheroid size. The general pattern is to expand and then shrink back the spheroids in order to include all cells on the edges. Then a series of functions are used to remove noise and the Watershed function separates fused or superimposed particles. The Analyze particles function is targeted to the specific spheroid size according to the black/white pixel ratio.
	if (ratio<0.001)
	{ 
		run("Maximum...", "radius=8");
		run("Fill Holes");
		run("Minimum...", "radius=8");
		//small spheroids require a more "gentle" function to clean up noise
		//run("Median...", "radius=2");
		run("Maximum...", "radius=25");
		run("Minimum...", "radius=25");
		run("Fill Holes");
		//run("Watershed");
		run("Analyze Particles...", "size=" + minSize + "-Infinity circularity=" + minCirc + "-1.00 show=[Overlay Outlines] display exclude include summarize");
	};
	if (ratio >=0.001 && ratio<0.01)
	{
		run("Maximum...", "radius=8");
		run("Fill Holes");
		run("Minimum...", "radius=8");
		//slightly bigger spheroids and a more rigorous function to remove noise
		//run("Remove Outliers...", "radius=10 threshold=0 which=Dark");
		//run("Watershed");
		run("Analyze Particles...", "size=" + minSize + "-Infinity circularity=" + minCirc + "-1.00 show=[Overlay Outlines] display exclude include summarize");
	};
	if (ratio>=0.01 && ratio<0.2)
	{
		run("Maximum...", "radius=8");
		run("Fill Holes");
		run("Minimum...", "radius=8");
		//run("Remove Outliers...", "radius=15 threshold=0 which=Dark");
		//run("Median...", "radius=4");
		//run("Watershed");
		run("Analyze Particles...", "size=" + minSize + "-Infinity circularity=" + minCirc + "-1.00 show=[Overlay Outlines] display exclude include summarize");
	};
	if (ratio>=0.2 && ratio<1)
	{
		//Very big spheroids generally do not need to be expanded much to fill up the edges. 
		run("Maximum...", "radius=3");
		run("Fill Holes");
		run("Minimum...", "radius=3");
		//Outliers and noise are removed rigorously
		//run("Remove Outliers...", "radius=50 threshold=0 which=Dark");
		run("Minimum...", "radius=30");
		run("Maximum...", "radius=30");
		//run("Watershed");
		run("Analyze Particles...", "size=" + minSize + "-Infinity circularity=" + minCirc + "-1.00 show=[Overlay Outlines] display exclude include summarize");
	};
	if (Overlay.size!=1 || isNaN(getResult("Circ.")))
		return newArray(-1,-1,-1,-1);
	else
		return newArray(getResult("Circ."),enhanceCont,upperThresh,ratio);
};


//This part actually runs the program
call("java.lang.System.gc");
run("Clear Results");

//Asks for the input folder which should have only the images to be analyzed
Dialog.create("Please choose the input folder");
Dialog.addDirectory("Input folder:", File.directory);
Dialog.show();

inputFolder = Dialog.getString();
inputImages = getFileList(inputFolder);

//Checks that the input folder is not empty
if (inputImages.length == 0)
{
	waitForUser("Error: the input folder is empty.");
	exit;
};

//Checks that the input folder has no subfolders
allImages = String.join(inputImages);
if (allImages.contains("/"))
{
	waitForUser("Error: the input folder contains subfolders." + "\n" + "It should contain only images to be processed.");
	exit;
};

//Checks that the input folder has only image files
for(i=0; i<inputImages.length; i++)
{
	dd = toLowerCase(inputImages[i]);
	isImage = (dd.contains(".tif") || dd.contains(".gif") || dd.contains(".jp") || dd.contains(".bmp") || dd.contains(".png"));
	if(!isImage) {
		waitForUser("Error: the input folder should contain only image files.");
		exit;
	};
};

//Detects whether the first image in the input folder has a scale
//Gets its pixel dimensions to compare to others
setBatchMode(true);
open(inputFolder + inputImages[0]);
//knownScale = 13.5*850/1317;
knownScale = 2000/462;
onePix = 1;
oneUnit = onePix;
toScaled(oneUnit);
fieldViewWidth = getWidth();
fieldViewHeight = getHeight();
close("*");

//Starts building the process overview dialog
Dialog.create("Process overview");
Dialog.addMessage(inputFolder);
Dialog.addMessage(inputImages.length + " images to be processed." + "\t" +" Image dimensions: " + fieldViewWidth + " x " + fieldViewHeight + " px");

if(onePix == oneUnit)
	Dialog.addMessage("No scale was detected in the images.");
else {
	Dialog.addMessage("A scale of " + (oneUnit/onePix) + " µm/px was detected automatically.");
	if (oneUnit/onePix > 1 && oneUnit/onePix < 20)
		knownScale = oneUnit/onePix;
};

if (oneUnit/onePix < 1 || oneUnit/onePix > 20)
{
	Dialog.addMessage("WARNING: the detected scale is outside of the usual range.");
	Dialog.addMessage("The value of " + oneUnit/onePix + " µm/px results in a field of view of " + (knownScale*fieldViewWidth) + " x " + (knownScale*fieldViewHeight) + " µm.");
	Dialog.addMessage("Please input the desired scale manually.");
};

//Asks what the scale of the images is
scaleChoices = newArray("Use the following global scale: ","Detect scale for each image  ","Do not use a scale  ");
Dialog.addRadioButtonGroup("", scaleChoices, 1, 3, "Use the following global scale: ")
Dialog.addNumber("Scale:", knownScale, 8, 12, "");
items1 = newArray("µm/px","px/µm");
Dialog.addChoice("", items1, "µm/px");

//Asks for known spheroid size
Dialog.addCheckbox("The approximate spheroid size range is known: (diameter)", false);
Dialog.addNumber("From", 500, 0, 12, "");
Dialog.addNumber("To", 10000, 0, 12, "");
items2 = newArray("µm","px");
Dialog.addChoice("", items2, "µm");

//Asks whether to output the processed images as TIFs with the outline of detected spheroids as overlays
Dialog.addCheckbox("Save output detected spheroids (as TIFs with overlays)", true);

//Asks whether to also output the processed images as JPGs with the flattened outline of the detected spheroids
Dialog.addCheckbox("Visualize output detected spheroids (generated as JPGs)", true);

//Asks whether the cross sectional area of the spheroids should be measured
Dialog.addCheckbox("Measure spheroid cross-sectional area", true);

//Asks whether the mean radial distance of the spheroids should be measured
Dialog.addCheckbox("Measure spheroid mean radial distance with the following angular resolution:", true);
Dialog.addNumber("Resolution:", 1, 2, 5, "deg");

Dialog.show();

//Gets all the choices the user selected from the dialog window
globalScale = Dialog.getRadioButton();
knownScale = Dialog.getNumber();
scaleUnits = Dialog.getChoice();
sizeIsKnown = Dialog.getCheckbox();
minSize = Dialog.getNumber();
maxSize = Dialog.getNumber();
sizeUnits = Dialog.getChoice();
//noOverlays = Dialog.getCheckbox();
outputTIFs = Dialog.getCheckbox();
outputFlatJPGs = Dialog.getCheckbox();
measureArea = Dialog.getCheckbox();
measureMRD = Dialog.getCheckbox();
res = Dialog.getNumber();
//visualizeProcess = Dialog.getCheckbox();
//quickenProcess = Dialog.getCheckbox();
//downScaleWidth = Dialog.getNumber();

//Creates output folder for the processed images with the editable outlines as overlays in TIFs
outputFolder = inputFolder + "/ProcessedImages/";
File.makeDirectory(outputFolder);

if (scaleUnits == "px/µm")
{
	knownScale = 1/knownScale;
	scaleUnits = "µm/px";
};

if (sizeIsKnown && sizeUnits == "µm" && (globalScale == "Do not use a scale  " || (globalScale == "Detect scale for each image  " && onePix == oneUnit)))
{
	showMessage("Note: Spheroid size range was input in µm but no scale was selected or detected." + "\n" + "Known size range will be set to false.");
	sizeIsKnown = false;
};
if (sizeUnits == "µm" && sizeIsKnown)
{
	minSize = minSize/knownScale;
	maxSize = maxSize/knownScale;
	sizeUnits = "px";
};

setBatchMode(false);

if (outputTIFs) {
	outputTIFFolder = inputFolder + "/ProcessedImages/outputTIF/";
	File.makeDirectory(outputTIFFolder);
};

//Creates output folder for the processed images with the flattened spheroid outlines as JPGs
if (outputFlatJPGs)
{
	outputJPGFolder = inputFolder + "/ProcessedImages/outputJPG/";
	File.makeDirectory(outputJPGFolder);
};

//Sets the measurements that are recorded for each spheroid
run("Set Measurements...", "area centroid shape feret's display add redirect=None decimal=4");

//Sets up the column headers for the results file
numColumns = 9;
//numColumns = 11;
if (!measureMRD)
	numColumns = numColumns - 4;	
if(!measureArea)
	numColumns = numColumns - 2;
header = "File" + "\t";
if(measureArea)
	header = header + "Cross-sectional Area (µm^2)" + "\t";
if(measureMRD)
	header = header + "Mean Diameter (µm)" + "\t" + "Mean Radial Distance (µm)" + "\t";
if(measureArea)
	header = header + "CSA (px^2)" + "\t";
if(measureMRD)
	header = header + "Mean Diameter (px)" + "\t" + "MRD (px)" + "\t";
header = header + "Circ." + "\t" + "Round";

//Gets date and time for naming results text file
getDateAndTime(year, month, dayOfWeek, dayOfMonth, hour, minute, second, msec);
month++;
if (month < 10) month = "0" + month;
if (dayOfMonth < 10) dayOfMonth = "0" + dayOfMonth;
if (hour < 10) hour = "0" + hour;
if (minute < 10) minute = "0" + minute;
dateTimeText = "" + year + "" + month + "" + dayOfMonth + "_" + hour + "" + minute;

//Generates text file to store all measurements
f = File.open(outputFolder + "Results_" + dateTimeText + ".txt");
print(f, "Scale:" + "\t" + knownScale + "\t" + "µm/px");
print(f, header);
print(f, "");
File.close(f);

//Generates text file to store enhancecontrast and threshold
gg = File.open(outputFolder + "Thingzs_" + dateTimeText + ".txt");
print(gg, "File:" + "\t" + "BestEnhanceCont" + "\t" + "BestUpperThresh" + "\t" + "Best b/w ratio");
print(gg, "");
File.close(gg);

waitForUser("To detect spheroid manually on image, adjust size with (-) and (+)" + "\n" + "Confirm with Shift or SPACE");
startTime = getTime();

//This is the cycle that runs through all images
for (i=0; i<inputImages.length; i++)
{ 
	redo = true;
	//xcentroid = 0;
	//ycentroid = 0;
	//autoCrop = false;
	detectSpheroid = newArray;
	enhanceCont = newArray(0.35,0.5,1,2.5,5,7.5,10);
	lowerThresh = 0;
	upperThresh = newArray;

	while (redo == true)
	{
		close("*");
		open(inputFolder + inputImages[i]);

		run("8-bit");
		
		actionOutput = action(inputFolder,inputImages[i],"ModifManual",0.35,0,0);
		detectSpheroid = Array.concat(detectSpheroid,actionOutput[0]);
		//setBatchMode("exit and display");
		//Table.setLocationAndSize(wide/8,high*5/16,wide/4,high/4,"Results");
		//Table.setLocationAndSize(wide/8,high*5/16,wide*0.75,high*9/16,"Summary");
		
		//Checks if there was a spheroid detected
		if (Overlay.size > 0)
		{
			if (Overlay.size==1)
			{
				//Sends particle detected to the ROI manager
				currentCirc = getResult("Circ.");
				currentRound = getResult("Round");
				currentArea = getResult("Area");
			}
			else if (Overlay.size>1)
			{
				//Since there were multiple particles detected it will select the one with the highest circularity
				sizeOverlay = Overlay.size;
				run("To ROI Manager");
				run("Remove Overlay");
				xValues = newArray();
				yValues = newArray();
				circValues = newArray();
				feretValues = newArray();
				roundValues = newArray();
				areaValues = newArray();
				selectRois = newArray();
				for (m=0; m<sizeOverlay; m++)
				{
					roiManager("Select", m);
					roiManager("Measure");
					xValues = Array.concat(xValues,getValue("X"));
					yValues = Array.concat(yValues,getValue("Y"));
					circValues = Array.concat(circValues,getValue("Circ."));
					feretValues = Array.concat(feretValues,getValue("Feret"));
					roundValues = Array.concat(roundValues,getValue("Round"));
					areaValues = Array.concat(areaValues,getValue("Area"));
					selectRois = Array.concat(selectRois,m);
				};
				run("Clear Results");
				n = Array.findMaxima(circValues,0);
				selectRois = Array.deleteIndex(selectRois, n[0]);
				roiManager("Select", selectRois);
				roiManager("Delete");
				run("From ROI Manager");
				roiManager("reset");
				close("Roi Manager");			
			
				currentCirc = circValues[n[0]];
				currentRound = roundValues[n[0]];
				currentArea = areaValues[n[0]];
				//if (attemptNum<enhanceCont.length+2)
				//{
				//	xcentroid = xValues[n[0]];
				//	ycentroid = yValues[n[0]];
				//	maxFeret = feretValues[n[0]];
				//};
			};
		
			//Sends particle detected to the ROI manager
			run("To ROI Manager");
		
			run("Revert");
			imgID = getImageID();
			run("From ROI Manager");
			Overlay.measure;
			currentCirc = getResult("Circ.");
			currentRound = getResult("Round");
			currentArea = getResult("Area");
			run("Clear Results");
			//File.append("" + inputImages[i] + "\t" + bestEnhanceCont + "\t" + upperThresh[upperThresh.length-1] + "\t" + ratio + "\t",outputFolder + "Thingzs_" + dateTimeText + ".txt");
			selectImage(imgID);
			//selectWindow("name");
			setBatchMode("exit and display");
			redo = getBoolean("Preview", "Redo", "Continue");
			setBatchMode("hide");
			//redo = false;
		}
		else if (Overlay.size == 0)
		{
			redo = getBoolean("Nothing detected", "Redo", "Skip");
			//redo = true;
		};
	};

	//Checks whether there was a single spheroid detected
	measurable = false;
	if (Overlay.size == 1)
		measurable = true;

	//Saves image as TIF with spheroid outline as overlay
	run("Overlay Options...", "stroke=none width=0 fill=none set apply show");
	//sets scale to predetermined values
	scaleText = "distance=100 known=" + knownScale*100 + " pixel=1 unit=µm";
	run("Set Scale...", "" + scaleText);
	if (measurable && outputTIFs)
		saveAs("Tiff", outputTIFFolder + inputImages[i]);
	else if (!outputTIFs)
	{
		run("Overlay Options...", "stroke=none width=5 fill=none set apply show");
		saveAs("Jpeg", outputJPGFolder + inputImages[i]);
	};
	
	call("java.lang.System.gc");

	//Prints the spheroid measurements in the previously created results text file
	if (measurable)
	{
		if (measureMRD)
			mrd = radialDistance(res);
		spheroidStats = "";
		spheroidStats = spheroidStats + inputImages[i] + "\t";
		if (measureArea)
			spheroidStats = spheroidStats + currentArea*knownScale*knownScale + "\t";
		if (measureMRD) {
			spheroidStats = spheroidStats + 2*mrd*knownScale + "\t";
			spheroidStats = spheroidStats + mrd*knownScale + "\t";
		};
		if (measureArea)
			spheroidStats = spheroidStats + currentArea + "\t";
		if (measureMRD) {
			spheroidStats = spheroidStats + 2*mrd + "\t";
			spheroidStats = spheroidStats + mrd + "\t";
		};
		spheroidStats = spheroidStats + currentCirc + "\t";
		spheroidStats = spheroidStats + currentRound + "\t";
		File.append(spheroidStats,outputFolder + "Results_" + dateTimeText + ".txt");
	}
	else
	{
		spheroidStats = "";
		spheroidStats = spheroidStats + inputImages[i] + "\t";
		if (measureArea)
			spheroidStats = spheroidStats + NaN + "\t" + NaN + "\t";
		if (measureMRD)
			spheroidStats = spheroidStats + NaN + "\t" + NaN + "\t" + NaN + "\t" + NaN + "\t";
		spheroidStats = spheroidStats + NaN + "\t" + NaN + "\t";
		File.append(spheroidStats,outputFolder + "Results_" + dateTimeText + ".txt");
	};
	
	run("Clear Results");
	close("Roi Manager");
	
	//Closes image if still open
	close("*");
	
	//Saves image as JPG with flattened spheroid outline
	if (outputFlatJPGs && outputTIFs && measurable)
	{
		outputTIFFiles = getFileList(outputTIFFolder);
		tifExtension = outputTIFFiles[outputTIFFiles.length-1];
		tifExtension = split(tifExtension,".");
		tifExtension = tifExtension[tifExtension.length-1];
		open(outputTIFFolder + File.getNameWithoutExtension(inputImages[i]) + "." + tifExtension);
		run("Overlay Options...", "stroke=none width=5 fill=none set apply show");
		saveAs("Jpeg", outputJPGFolder + inputImages[i]);
		close("*");
	};
	
	close("Summary");
	
	showProgress(i, inputImages.length);
	//waitForUser("Just finished image" + "\n" + "All Circs: " + String.join(detectSpheroid," ") + "\n" + "All Upper Threshs: " + String.join(upperThresh," ") + "\n" + "Best Enhance Cont: " + bestEnhanceCont);
	
};

close("Roi Manager");
close("Results");
close("Summary");
	
currentTime = getTime();
elapsedTime = (currentTime-startTime)/60000;
wait(5000*inputImages.length);
//Displays end message
waitForUser("Process completed in " + elapsedTime + " minutes.");

//filedel = File.delete(outputFolder + "Thingzs_" + dateTimeText + ".txt");

setBatchMode(false);
