## Supplementaries for "Investigation of the threonine metabolism of *Echinococcus multilocularis*: the threonine dehydrogenase as a potential drug target in alveolar echinococcosis": S4 File.docx

**Analysis of amino acids in supernatant samples**

For quantitative amino acid analysis, the supernatant samples were diluted 1:20 with MilliQ water and 10 µL thereof were pipetted into a glass tube (5x60 mm). After addition of 5µL EDTA-solution (Merck Titriplex III, 2 mg/mL in water), the samples were dried on a vacuum concentrator. 10 µLof a methanol/water/triethylamine (2/2/1) solution were added, followed by drying for 10 min. The sample was reconstituted in 10 µL of a methanol/water/triethylamine/phenylisothiocyanate (70/10/10/1) solution and incubated for 20 min at room temperature before being dried for 45 min.

The amino acid standard H (Thermo Fisher Scientific) was diluted with 0.1% trifluoroacetic acid in water to a concentration of 125 nmol/mL. 10 μL of this standard solution were treated as described above for the samples.

Analysis was performed by high-performance liquid chromatography (HPLC) using a Nova-Pak C18 (4µm, 60Å, 3.9mm x 125 mm) reversed-phase column (Waters, Baden-Dättwil, Switzerland) on an UltiMate 3000 HPLC system (Thermo Fisher Scientific, Reinach, Switzerland). The column was held at 50°C and a gradient ranging from 4% eluent B to 50% eluent B in 12 min was applied with a flow rate of 1 mL/min. Eluent A consisted of 96% ammonium acetate (0.14 M) and 575 µl triethyl amine in water (adjusted with acetic acid to pH 6.4) and 4% acetonitrile. Eluent B was 40% Milli-Q water and 60% acetonitrile. Derivatized samples and standard were reconstituted in 50 μL eluent B and 20 μL were loaded on the column.
