## Supplementaries for "Investigation of the threonine metabolism of *Echinococcus multilocularis*: the threonine dehydrogenase as a potential drug target in alveolar echinococcosis": S5 File.docx

**S5 File. Cloning and recombinant expression of EmTDH and MmTDH**

***emtdh* (EmuJ_000511900, sequence obtained from WormBase ParaSite (https://parasite.wormbase.org))**

ATGCTGGTTCGTCAGGCAACCCTCGCCTGCCAAAAGACCCTCTCCTCCTTCGCTGCCCGTGCTGCAGCTGTCGGTGTCAATTTGAGAGGCCATCATGGCACTGGAGCATGTTCCCACCAAACTCCGAAAATCCTAATCACTGGCGGTCTTGGTCAACTTGGAATGCCCCTAGCACGCGTCTTTCGTTCCAAATATGGCCAGGACTCTGTCATCCTGACTGACATTAAGAAACCCGATCCTGCTGTGAAGGATGTCGGCCACTTTGAGTTCGCTGACATCATGGATGTTGATGGATTGAGAGAACTGGTGGTGGAGCATCGTATTGACTGGCTCATCCACTTCTCAGCTCTGCTCTCCGCTGTTGGGGAACAAAACACCCCGTTGGCAATGCAGGTGAATATTGGAGGTCTTCATAACATTTTCAACGTGGCGCGAGAATTCAATCTCCGTCTGTTCATTCCCAGCACCATCGGAGCCTTTGGACCCAATTCACCGCGTAATCCGACACCTGATGATTGTATTCAACAACCGAATACCATATATGGTGTGAGTAAGGTTCATGCTGAACTACTCGGCATGTATTTGAACCACAGATACGGACTTGACTTCAGATCTCTCCGTCTTCCCGGTGTCATCTCTGCTGATGTCGAACCCGGCGGTGGCACAACAGACTATGCCATTCACATTTTCAAGTACGCAATGCGTGGTGAACCCTATCCTTGCTTTCTGAGCCAGAACACGAGGCTGCCGATGATTTGGATCGATGATGTTATTCGAGCCATGGTCGAGGTGATGGAATGCCCAAGAGAGAAGTTGAAGAGGTGTGTTTACAATTTGGCGGGTATTTCGTTCACCCCGAAGGAGATTAATGAAGCCATGCTTCGTGGAATGCACCGGCATGGATTGGCGAAGCAGTTGAAGACGTATGAGCTCAAATACGACGTTGACTTCCGTCAAAAGATTGCTGATAGTTGGCCAGAAGTTTTGGATGATGCCAATGCCAGACGAGACTGGGGCTGGAGGCACGAGGTCGACATCGATGGCATGGTGGACAAGATGTTTGAGTACCTCAAGGCCCACTCAACCTAA

***mmtdh* (ENSMUST00000022522.15, sequence obtained from the National Library of Medicine (https://www.ncbi.nlm.nih.gov)**

ATGCTGGTTCGTCAGGCAACCCTCGCCTGCCAAAAGACCCTCTCCTCCTTCGCTGCCCGTGCTGCAGCTGTCGGTGTCAATTTGAGAGGCCATCATGGCACTGGAGCATGTTCCCACCAAACTCCGAAAATCCTAATCACTGGCGGTCTTGGTCAACTTGGAATGCCCCTAGCACGCGTCTTTCGTTCCAAATATGGCCAGGACTCTGTCATCCTGACTGACATTAAGAAACCCGATCCTGCTGTGAAGGATGTCGGCCACTTTGAGTTCGCTGACATCATGGATGTTGATGGATTGAGAGAACTGGTGGTGGAGCATCGTATTGACTGGCTCATCCACTTCTCAGCTCTGCTCTCCGCTGTTGGGGAACAAAACACCCCGTTGGCAATGCAGGTGAATATTGGAGGTCTTCATAACATTTTCAACGTGGCGCGAGAATTCAATCTCCGTCTGTTCATTCCCAGCACCATCGGAGCCTTTGGACCCAATTCACCGCGTAATCCGACACCTGATGATTGTATTCAACAACCGAATACCATATATGGTGTGAGTAAGGTTCATGCTGAACTACTCGGCATGTATTTGAACCACAGATACGGACTTGACTTCAGATCTCTCCGTCTTCCCGGTGTCATCTCTGCTGATGTCGAACCCGGCGGTGGCACAACAGACTATGCCATTCACATTTTCAAGTACGCAATGCGTGGTGAACCCTATCCTTGCTTTCTGAGCCAGAACACGAGGCTGCCGATGATTTGGATCGATGATGTTATTCGAGCCATGGTCGAGGTGATGGAATGCCCAAGAGAGAAGTTGAAGAGGTGTGTTTACAATTTGGCGGGTATTTCGTTCACCCCGAAGGAGATTAATGAAGCCATGCTTCGTGGAATGCACCGGCATGGATTGGCGAAGCAGTTGAAGACGTATGAGCTCAAATACGACGTTGACTTCCGTCAAAAGATTGCTGATAGTTGGCCAGAAGTTTTGGATGATGCCAATGCCAGACGAGACTGGGGCTGGAGGCACGAGGTCGACATCGATGGCATGGTGGACAAGATGTTTGAGTACCTCAAGGCCCACTCAACCTAA

Potential mitochondrial target signal sequences were predicted via TargetP 2.0 (1). *Emtdh*, without the predicted mitochondrial target signal, was amplified from cDNA obtained from *in vitro* cultured *E. multilocularis* metacestode vesicles using primers EmTDH_FW (5’-CACCCACCAAACTCCGAAAATCC-3’) and EmTDH_RV (5’-TTAGGTTGAGTGGGCCTTGAG-3’) in final concentrations of 0.5 µM via PCR using Q5 Polymerase (New England Biolabs). recMmTDH was amplified from 200 ng of pUCIDT-AMP plasmid containing a 1007 bp fragment of MmTDH without the predicted mitochondrial target sequence (ordered from LubioScience, Lucerne, Switzerland) using primers MmTDH_FW (5’- CACCCACTCCACCTCGATCTCGG-3’) and MmTDH_RV (5’-TTAGTTCACTTGGGCAACTCTGG-3’). The PCR reaction consisted of an initial denaturation step at 98°C for 2 min, followed by 35 cycles of 98°C for 10 seconds, 60°C for 30 seconds and 72°C for 2 minutes. A final elongation step at 72°C for 5 min was included. PCRs were run on a 1% agarose gel for 30 minutes at 100 V with TBE buffer (90 mM Tris-HCl pH 8.0, 90 mM boric acid, 2 mM EDTA) and ethidium bromide (200 ng/mL). Then, gels were imaged using a Vilber CX.5.TS Edge E-Box (Île-de-France, France) gel imaging system. The PCR for EmTDH resulted in one clear band (S5 File Fig 1) and was purified using the High Pure PCR Product Purification Kit (Roche, Basel, Switzerland) according to the manufacturer’s protocol. The PCR for MmTDH still contained parts of the vector and was thus cut out of the gel and purified using the Zymoclean™ Gel DNA Recovery Kit (Zymo Research, Irvine, USA) according to the manufacturer’s protocol.

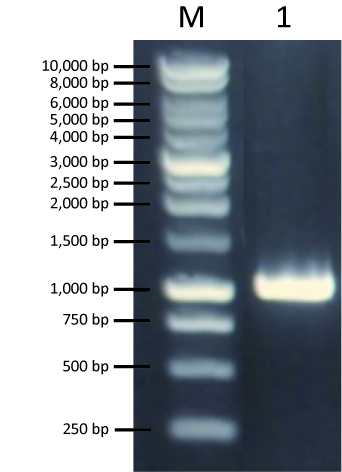

**S5 File Fig 1: Amplification of EmTDH from cDNA obtained from *in vitro* grown *E. multilocularis* metacestode vesicles.** As a Marker (M), Bentchtop 1 KB ladder (Promega, Dübendorf, Switzerland) was used.

Both EmTDH and MmTDH were cloned into the pET151/D-TOPO® vector using the Champion™ pET151 Directional TOPO™ Expression Kit (Fisher Scientific AG, Reinach, Switzerland) and transformed into *Escherichia coli* Top10 according to the manufacture’s protocol. Clones were picked and tested via colony PCRs for the expected fragment size with the vector-specific forward primer T7_FW (5’- TAATACGACTCACTATAGGG-3’) and an insert-specific reverse primer. The PCR reaction consisted of GoTaq® Green Master Mix (Promega, Dübendorf, Switzerland), 1 μL of a water suspension in which single clones were dissolved and primers at final concentrations of 1 µM. The PCR program consisted of an initial denaturation step at 94°C for 3 min, followed by 35 cycles of 94°C for 30 seconds, 58°C for 30 seconds and 72°C for 2 minutes. A final elongation step at 72°C for 5 min was included. PCRs were run onto a 1% agarose gel for 30 minutes at 100V (S5 File Fig 2).

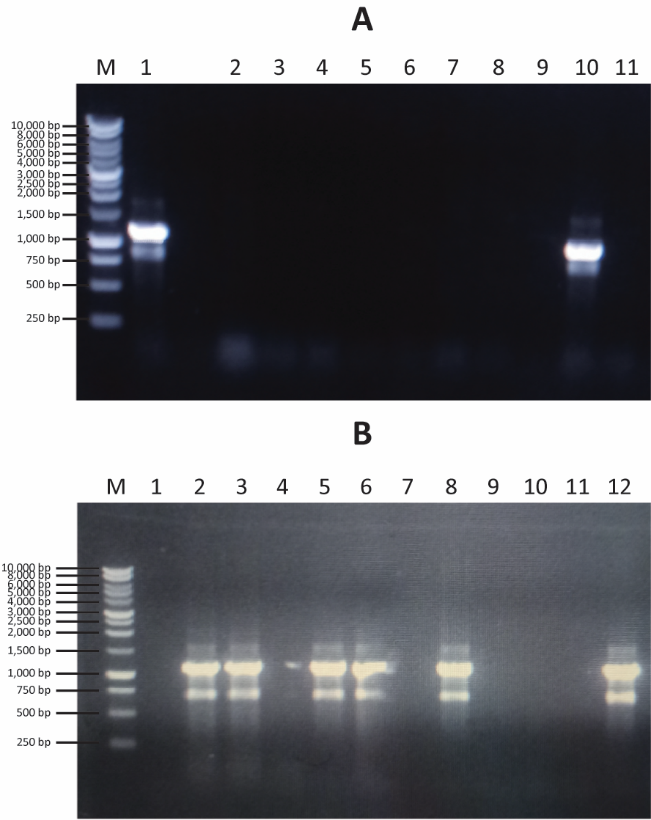

**S5 File Fig 2: Colony PCRs.** Shown are colony PCRs with *E. coli* Top10 transformed with pET151/D-TOPO®_EmTDH (A) or pET151/D-TOPO®_MmTDH (B). 1% agarose gels were run for 30 minutes at 100V. As a Marker (M), Bentchtop 1 KB ladder was used. Numbers stand for the respective clones.

Plasmid DNA was isolated from positive clones (clones 1 and 10 for EmTDH and clones 2 and 8 for MmTDH) using the ZymoPURE™ Plasmid Miniprep Kit (Zymo Research, Irvine, USA) and Sanger sequencing was conducted at Microsynth AG (Balgach, Switzerland). Sequences were compared using BioEdit (2) The EmTDH insert sequence was almost identical to the reference sequence (EmuJ_000511900) with the exception of transitions T->A at position 975 and a transversion A->G at position 977 which did not alter the amino acid sequence. An additional 8 amino acids were present between the end of the EmTDH sequence and the stop codon (S5 File Fig 3). The MmTDH sequence on the other hand was completely identical to the reference sequence (ENSMUST00000022522.15) (S5 File Fig 4).

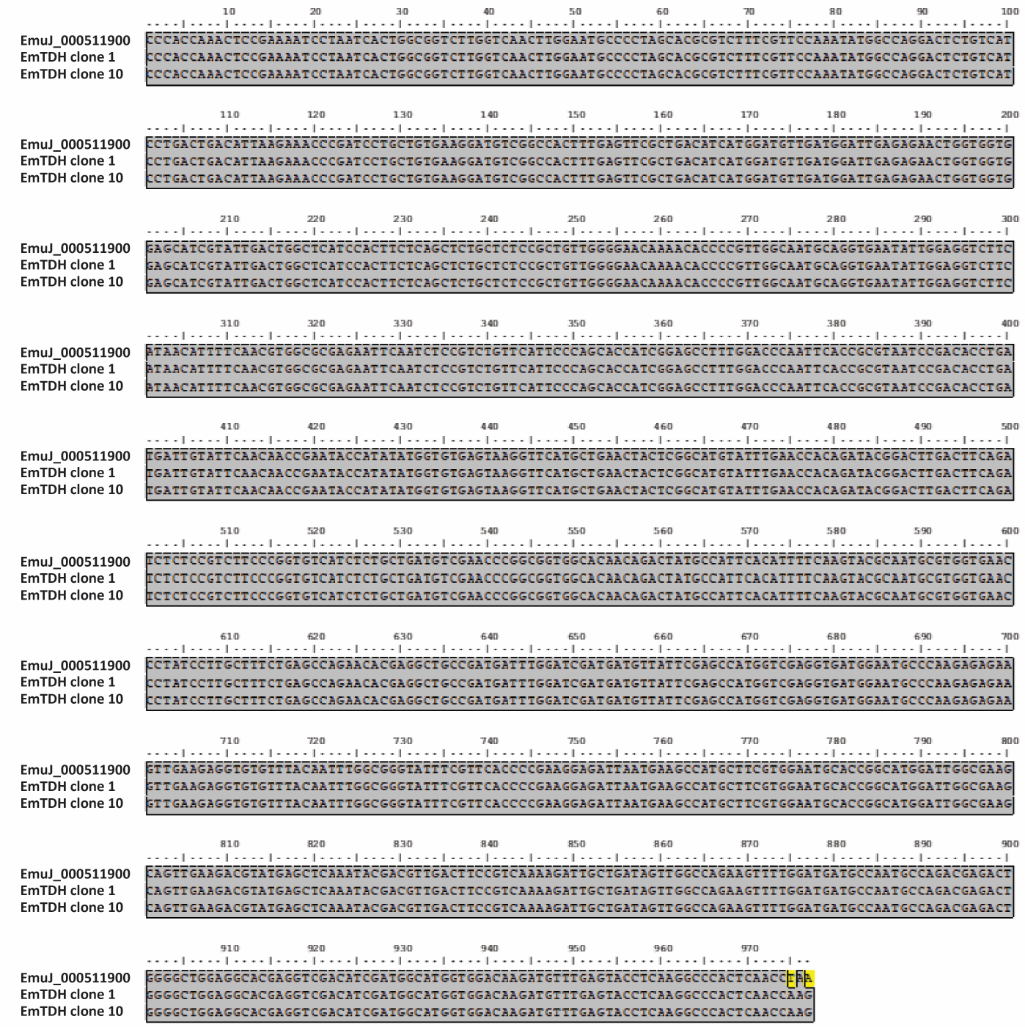

**S5 File Fig 3: Alignment of recEmTDH to the reference sequence EmuJ_000511900.** The alignment was prepared in BioEdit (2).

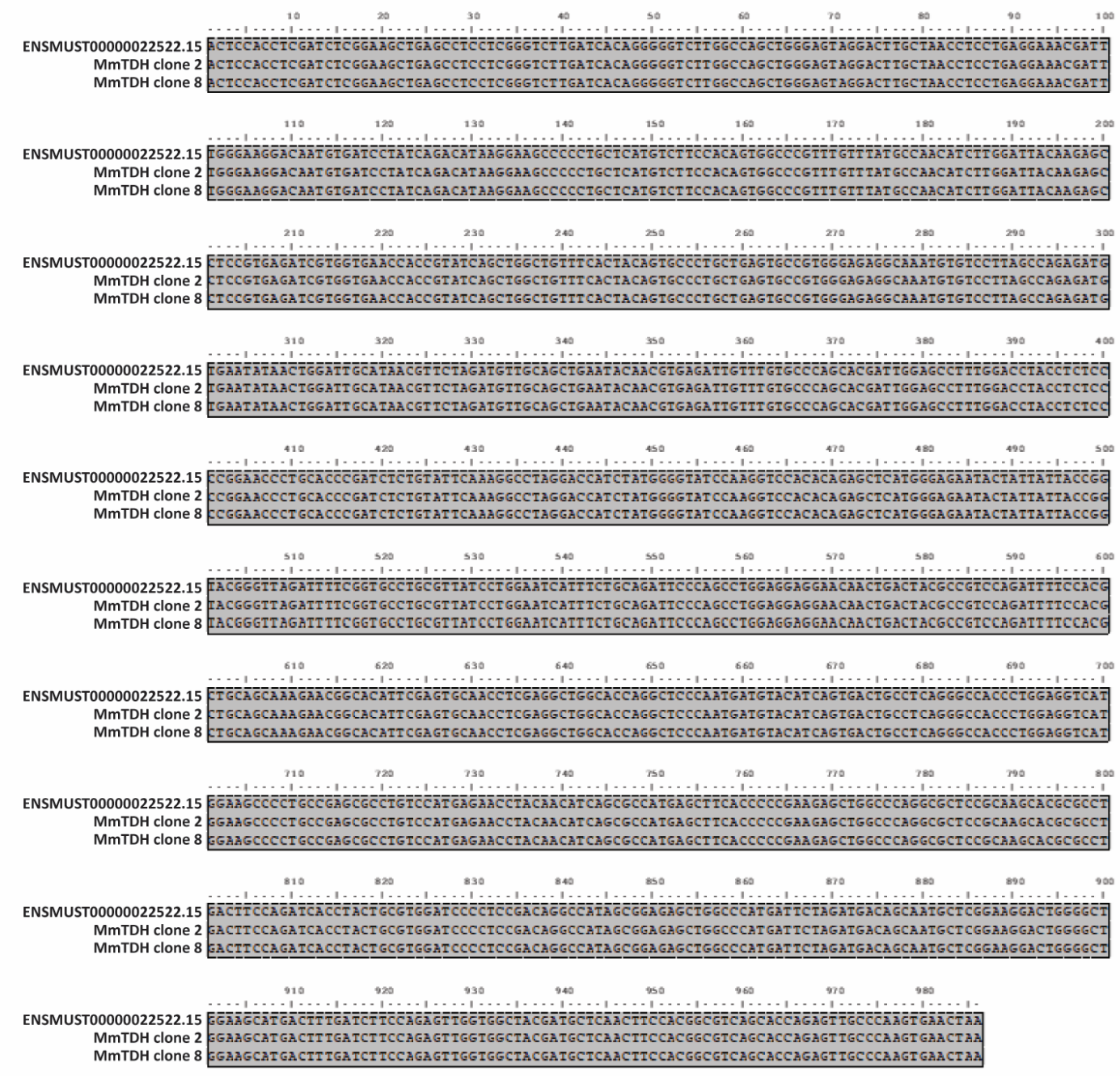

**S5 File Fig 4: Alignment of MmTDH clones 2 and 8 to the reference sequence ENSMUST00000022522.15.** The alignment was prepared in BioEdit (2).

*E. coli* BL21 were transformed with 100 ng of plasmids containing either EmTDH or MmTDH, plated onto LB agar plates and incubated overnight at 37°C. Single clones were incubated overnight at 37°C under mild shaking in Luria broth (LB) (10% (w/v) bactotryptone, 5% (w/v) bacto yeast extract, 5% (w/v) NaCl) containing 100 μg/mL carbenicillin. The next day, liquid cultures were diluted 1:20 with new LB medium containing 100 μg/mL carbenicillin and allowed to incubate for three hours at 37°C under mild shaking. Once the optical density reached a value of 0.6 to 0.9, protein expression was induced by adding IPTG to a final concentration of 1 mM, a non-induced culture was done simultaneously. The cultures were then incubated for another four hours at 37°C under mild shaking. RecEmTDH and recMmTDH were purified via the Macherey-Nagel™ Protino™ Ni-TED-IDA 1000 Kit (Fisher Scientific, Schwerte, Germany) according to the manufactures protocol and eluates were loaded onto a 12% sodium dodecyl sulfate polyacrylamide gel. Correct protein size of 41.4 kDa for recEmTDH and 40.3 kDa for recMmTDH was confirmed by western blot using a mouse monoclonal anti-HIS tag antibody and an anti-mouse IgG (h+l) ap conjugate as secondary antibody (Promega, Dübendorf, Switzerland) (S5 File Fig 5).

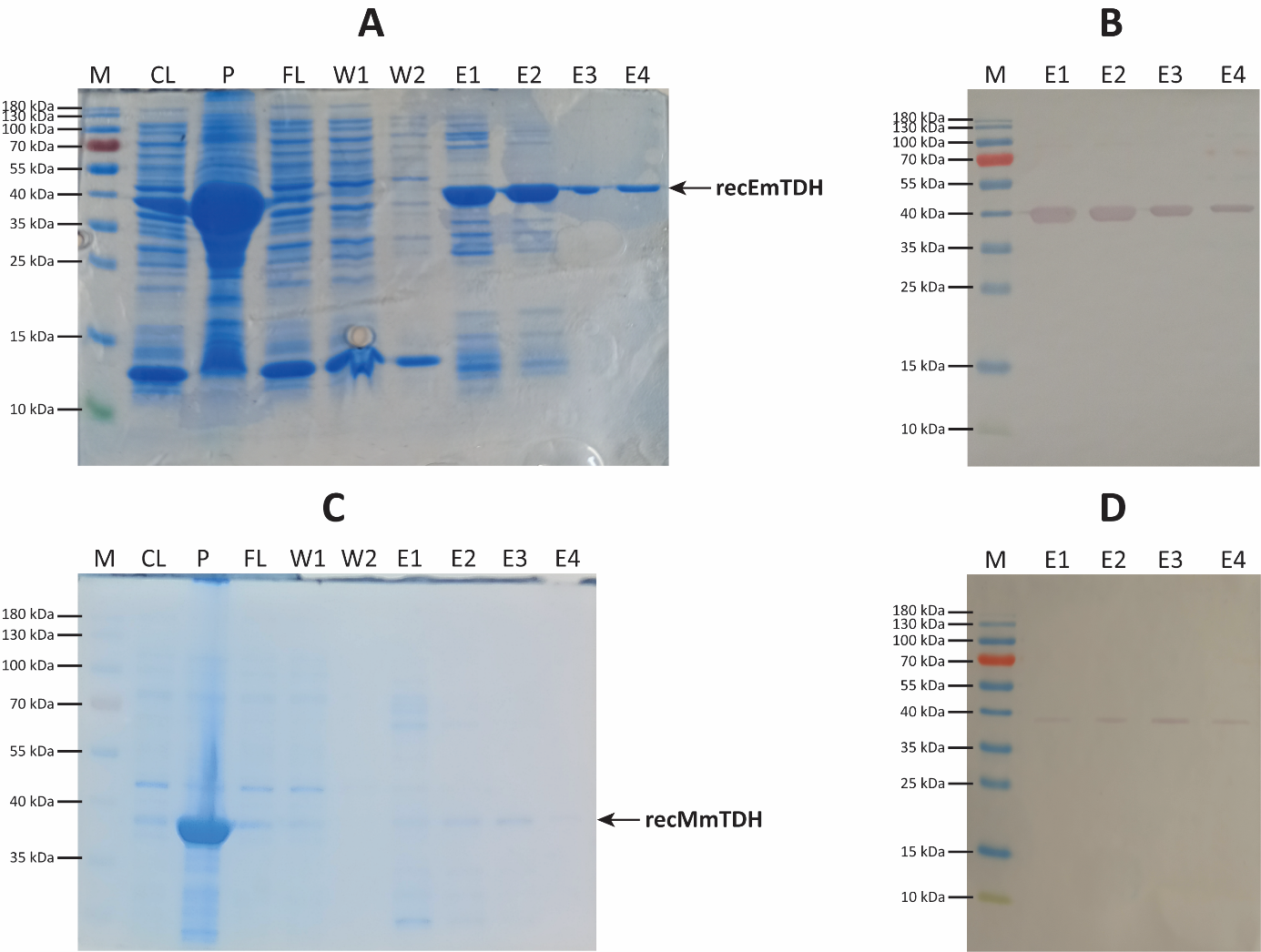

**S5 File Fig 5: HIS-tag purification of recEmTDH and recMmTDH visualized by SDS gel electrophoresis, followed by subsequent staining by colloidal coomassie G-250 and protein identity validation via Western Blot.** A, Coomassie stained 12% SDS containing samples extracted from *E. coli* BL21 transformed with pET151/D-TOPO®_EmTDH). B, Western Blot with eluates of 1 to 4 of purified recEmTDH using a monoclonal anti-HIS tag antibody and an anti-mouse IgG (h+l) ap conjugate as secondary antibody. C, Coomassie stained 12% SDS containing samples extracted from *E. coli* BL21 transformed with pET151/D-TOPO®_MmTDH. D, Western Blot with eluates of 1 to 4 of purified recMmTDH using a monoclonal anti-HIS tag antibody and an anti-mouse IgG (h+l) ap conjugate as secondary antibody. M: marker, CL: cleared lysate, P: pellet, FL: flow through, W: wash (1-2), E: eluate (1-4).

Finally, enzymatic activity was tested for recEmTDH and recMmTDH (see main manuscript body).
