## Supplementary figures and images for "Investigation of the threonine metabolism of *Echinococcus multilocularis*: the threonine dehydrogenase as a potential drug target in alveolar echinococcosis"

### S1 Fig.tif

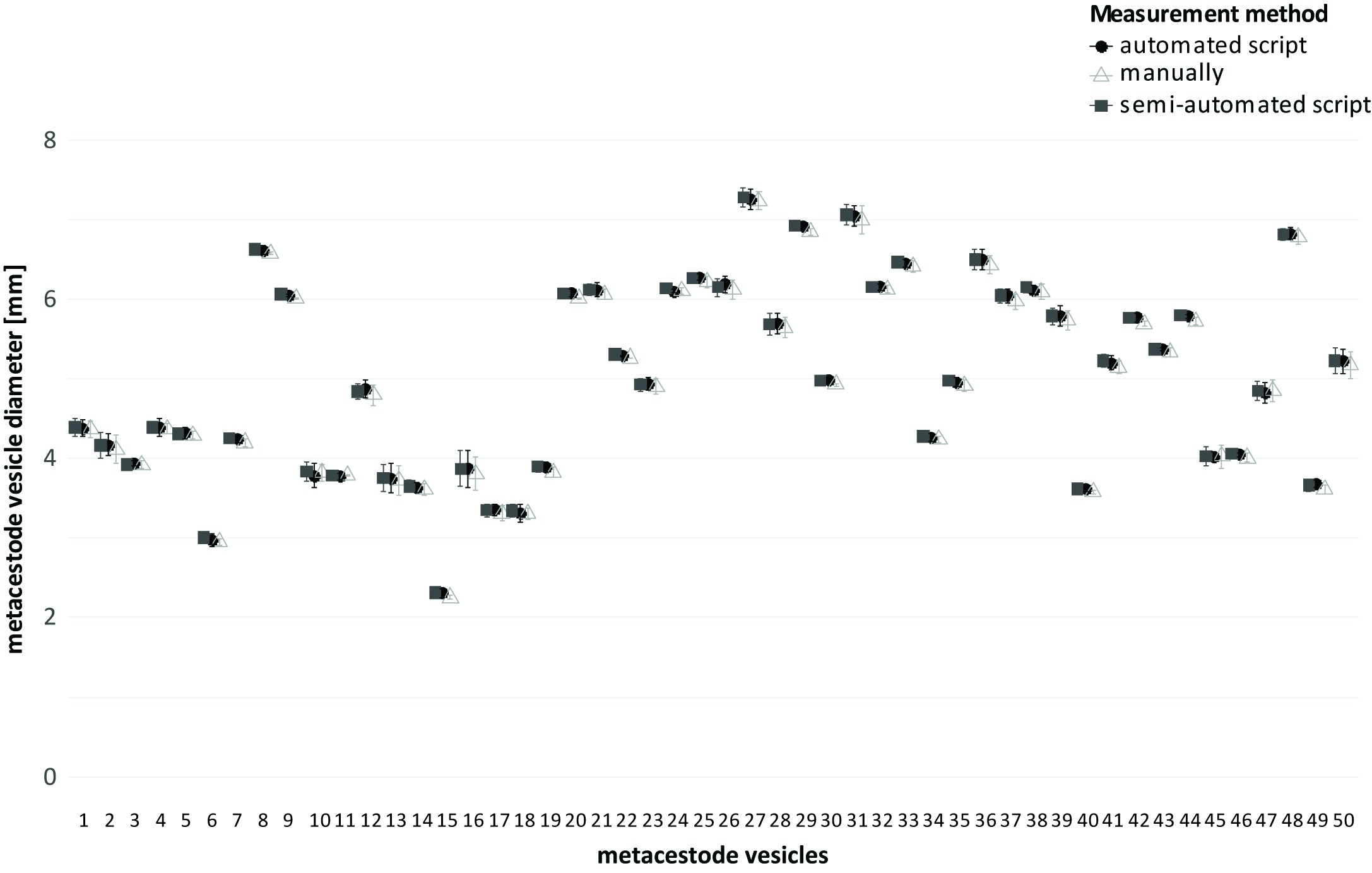

### S2 Fig.tif

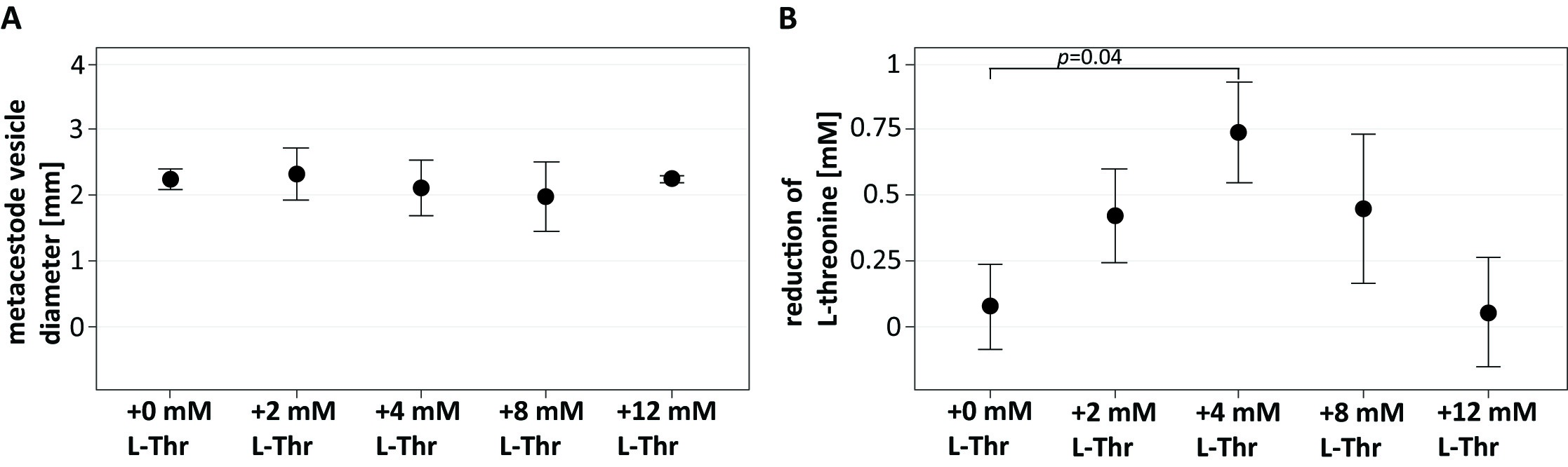

### S3 Fig.tif

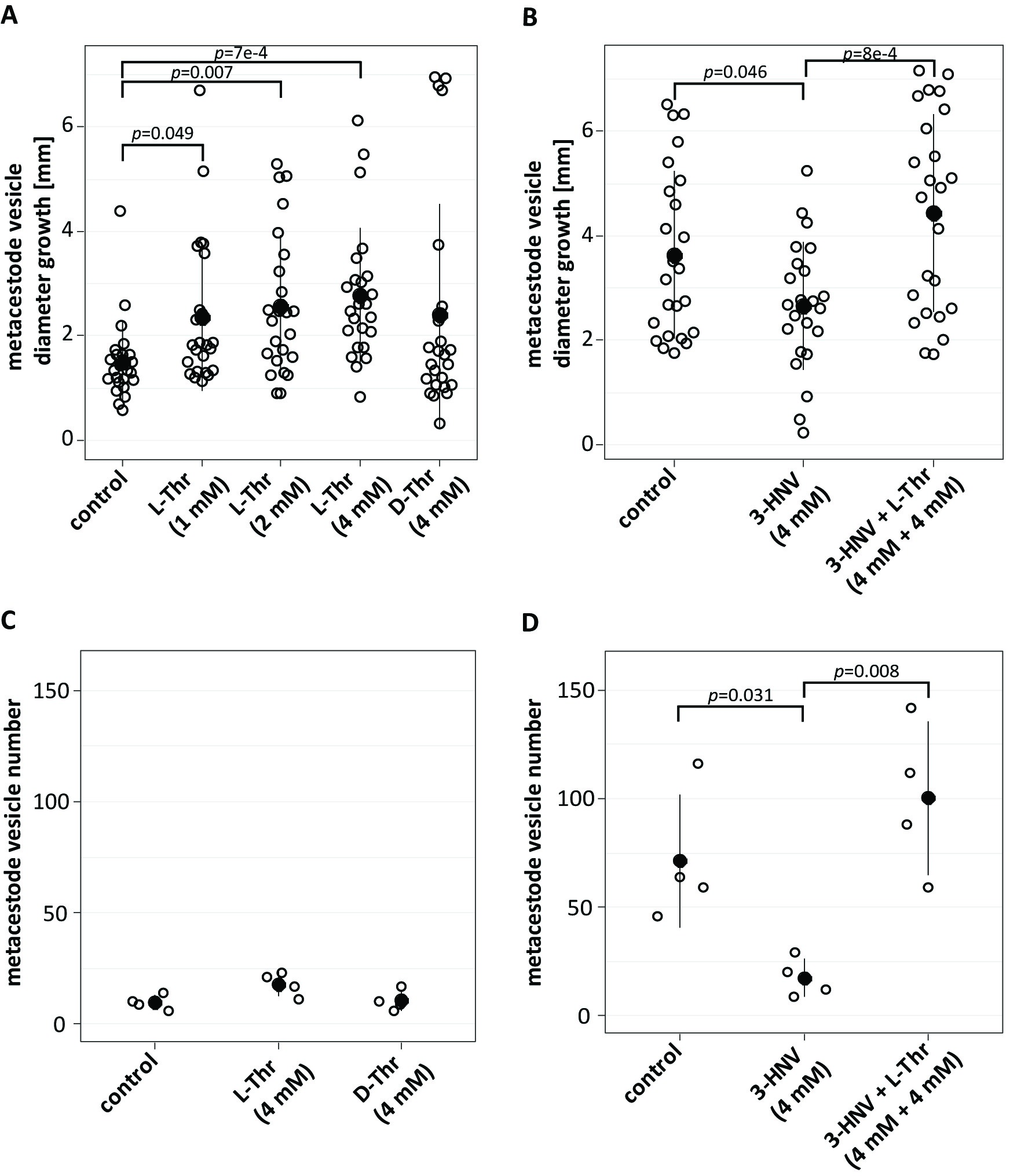

### S4 Fig.tif

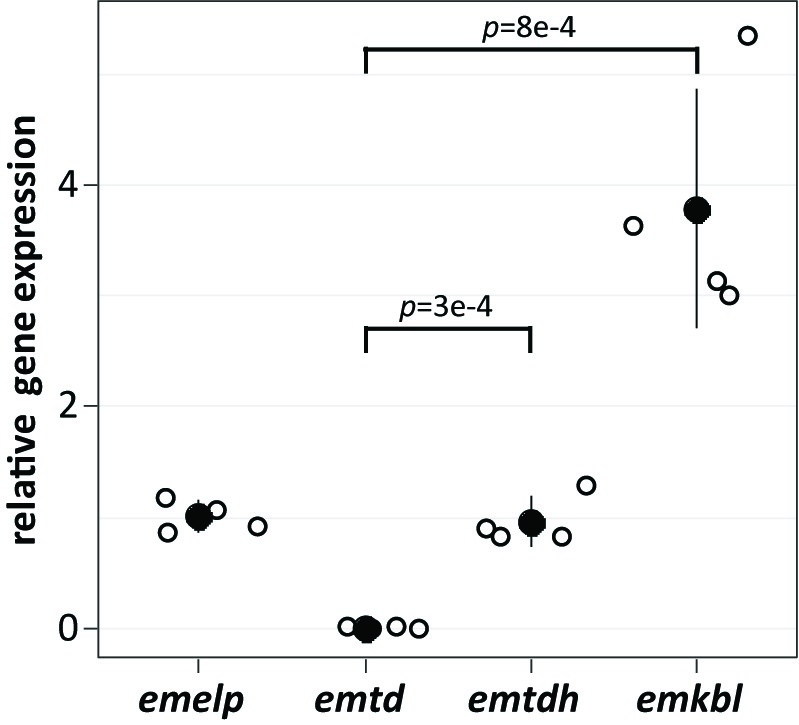

### S5 Fig.tif

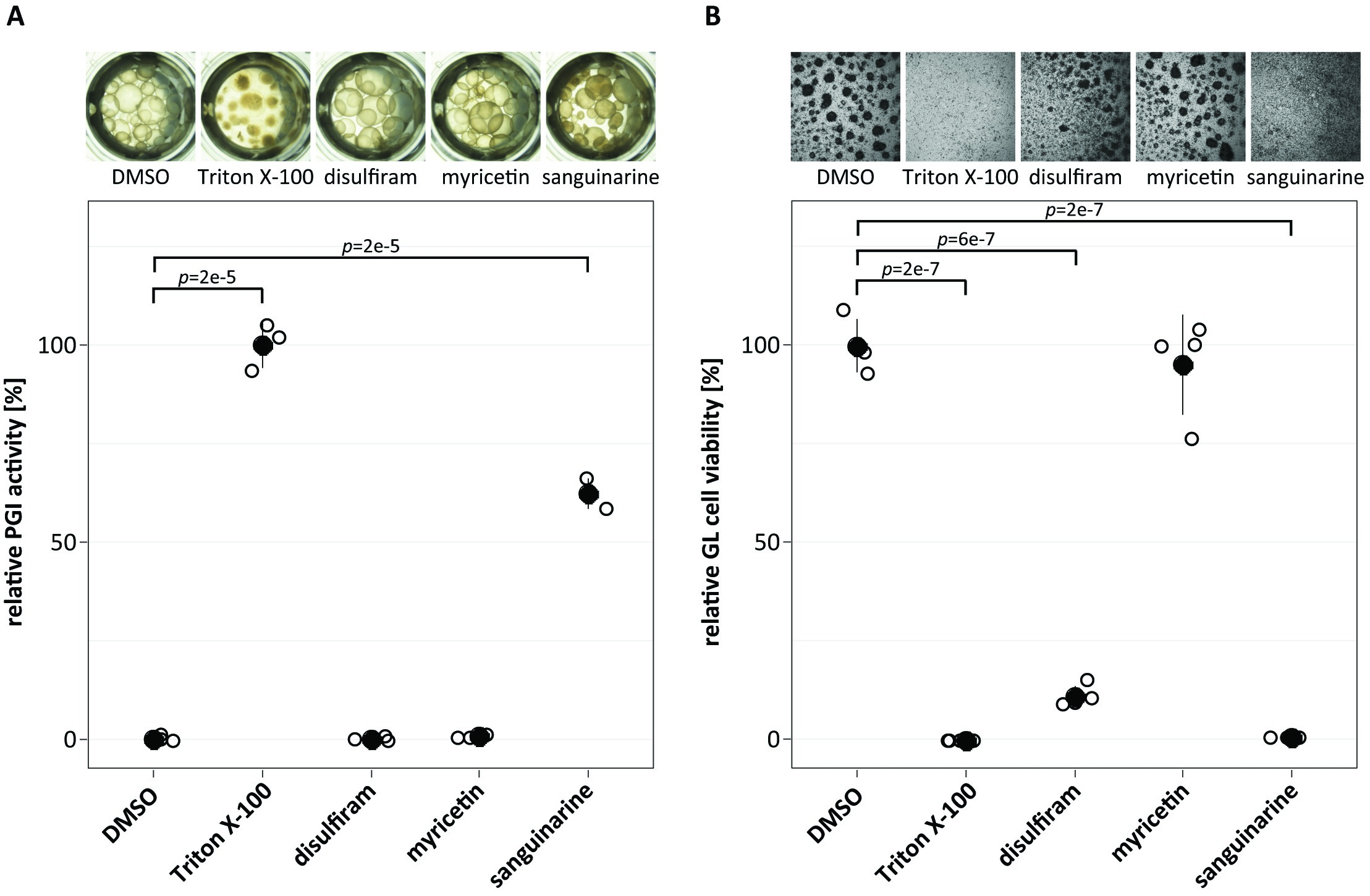
